# A modified cyclosporine enhances lentivector transduction *ex vivo* and *in vivo* by degrading IFITM3

**DOI:** 10.1101/2025.08.26.669098

**Authors:** Dara Annett, Kate L. Morling, Bethan J. Critchley, Ben Graham, Valeria Pingitore, Thomas E. Whittaker, Nada Kurdi, Justin Warne, Diego León-Rico, Nihansu Kuru, Sara Toros, Tasnim Nurullah, Aura Hare, Matthew V. X. Whelan, Edith A. W. Chan, Richard S. B. Milne, Lydia S. Newton, Hashim Ali, Kate Powell, Matteo Rizzi, Adrian J. Thrasher, Giorgia Santilli, Lucy G. Thorne, David L. Selwood, Greg J. Towers

## Abstract

Intrinsic innate immune barriers have evolved to suppress viral infection and can reduce effective gene delivery in gene therapy. We have developed BG147, a novel cyclosporine A analogue, optimised via structure-guided design to prevent inhibition of HIV cofactor Cyclophilin A and to specifically inhibit interferon-induced transmembrane proteins (IFITM1-3). BG147 enhances VSV-G pseudotyped lentiviral vector transduction *ex vivo* in hematopoietic stem and progenitor cells (HSPCs) and in *in vivo* ocular gene therapy of photoreceptor cells in mice. Upon BG147 treatment, IFITM proteins are mislocalised and degraded through lysosomal acidification-dependent pathways. IFITM3 levels functionally return in cells 96 h after BG147 washout. BG147 promises to transform *ex vivo* and *in vivo* eye gene therapies by transiently inhibiting intrinsic immune barriers mediated by IFITM proteins to enhance a wide range of protocols.

**One Sentence Summary:** Modified cyclosporine, BG147, enhances lentivector gene therapy transduction, *ex vivo* in HSPC and *in vivo* in mouse photoreceptors, by degrading IFITM3.

## INTRODUCTION

Monogenic disorders can be treated using gene therapy approaches to correct, replace, or add therapeutic genes. Inborn errors of immunity (IEIs) are tractable targets because hematopoietic stem and progenitor cells (HSPCs), isolated from the bone marrow, or mobilized into peripheral blood, are accessible for *ex vivo* approaches and autologous reinfusion of transduced cells, carries lower risks than allogeneic HSPC transplantation. Lentiviral vectors (LV) are popular and effective tools for gene delivery into HSPCs given their safe integration profile, improved from first generation gammaretroviral vectors, and their ability to transduce quiescent cells. Self-inactivating LVs are used clinically for several IEIs(*1, 2*) as well as for metabolic disorders that benefit from cross-correction by blood-derived cells(*3, 4*). However, for many conditions, high multiplicity of infection and extended protocols (e.g., two-hit approaches) are required resulting in high costs and potentially impacting graft quality(*5–7*).

The future of gene therapy lies in *in vivo* approaches which are cheaper and less invasive. Ocular gene therapy has been particularly successful but the large sizes of some therapeutic genes, such as ABCA4 to treat Stargardt disease, undermines the use of otherwise effective Adeno-associated viral (AAV) vectors(*8*). LV provide large capacity, stable integration but poorly transduce photoreceptor cells(*9–15*). Improving LV-based transduction is therefore a key goal. One barrier to LV transduction is the IFITM (interferon-induced transmembrane protein) family(*16–19*). IFITM1-3 cycle through the plasma membrane, endosomal, and lysosomal compartments and inhibit divergent viruses by blocking viral fusion with host membranes. IFITM3 is the most effective IFITM against the VSV-G envelope. It is predominantly endosomal and restricts viruses utilising endosomal fusion entry pathways(*20, 21*). Mutations that mis-localise IFITM3 reduce antiviral activity-consistent with a location specific inhibition mechanism(*22*). The molecular details are not well understood but IFITM3 may alter endosome pH or membrane composition and/or rigidity to impede fusion leading to virus degradation in lysosomes(*23–28*).

Cyclosporine A (CsA) causes IFITM3 degradation, and enhances LV entry(*18, 29*), but also causes downstream inhibition of LV infectivity by inhibiting the HIV-1 cofactor cyclophilin A (CypA), undermining its use as an LV transduction enhancer (TE). CypA inhibition prevents CypA protecting LV capsids from restriction by the antiviral protein tripartite motif protein 5 (TRIM5)(*30*). TRIM5 binding to capsids stimulates ubiquitin chain synthesis, proteasome recruitment and capsid destruction as well as innate immune signalling(*31–33*). CypA also influences capsid-binding cofactor activity, likely through dynamic allosteric effects on capsid, controlling the timing and location of uncoating and thus integration targeting (*34–36*). We showed previously that the non-immunosuppressive macrocycle, cyclosporine H (CsH), increased HSPC transduction efficiency more effectively than CsA because it does not inhibit CypA(*18, 37*). Like CsA, CsH enhances transduction by causing IFITM3 degradation(*18*). However, CsH is a minor product of CsA production from *Tolypocladium inflatum,* making it expensive and reducing suitability for clinical use(*38*). In this study, we have taken a structure-activity-relationship (SAR) approach to developing a CsA-derived TE, BG147, that causes IFITM trafficking to endolysosomes and degradation, enhancing LV transduction in HSPC without measurable CypA inhibition. We found that BG147 is additive with protamine sulphate (PS), a common TE reported to promote LV attachment to target cell membranes(*39, 40*). Unexpectedly, we found that Lentiboost (LB), a commercial TE reported to work through enhancing LV attachment to target cells, also inhibits IFITM3. We have also shown that BG147 can improve photoreceptor cell transduction when co-administered with LV in a mouse ocular gene therapy model. We propose BG147 as a highly effective TE for *ex vivo* HSPC and *in vivo* gene therapy and as a probe for IFITM biology.

## RESULTS

### Subhead 1: Modified cyclosporine analogues enhance LV transduction

CsA has immunosuppressive activity through acting as a molecular glue to create a ternary inhibitory complex between CsA, CypA and calcineurin (CN) (*41*). This inhibits T-cell activation by preventing de-phosphorylation and nuclear localisation of transcription factor NFAT(*41*). Importantly, chemical modification of CsA can selectively abrogate calcineurin inhibition and immunosuppression(*42, 43*). By exploring tractable chemistry at the 1[Bmt] and the 3[Sar] residues of CsA, we synthesised a SAR series of thirty CsA analogues (Fig. 1A, Table 1). These molecules are expected to vary in biological activities allowing for selection of transduction enhancement properties through further chemical modification.

**Fig. 1:**
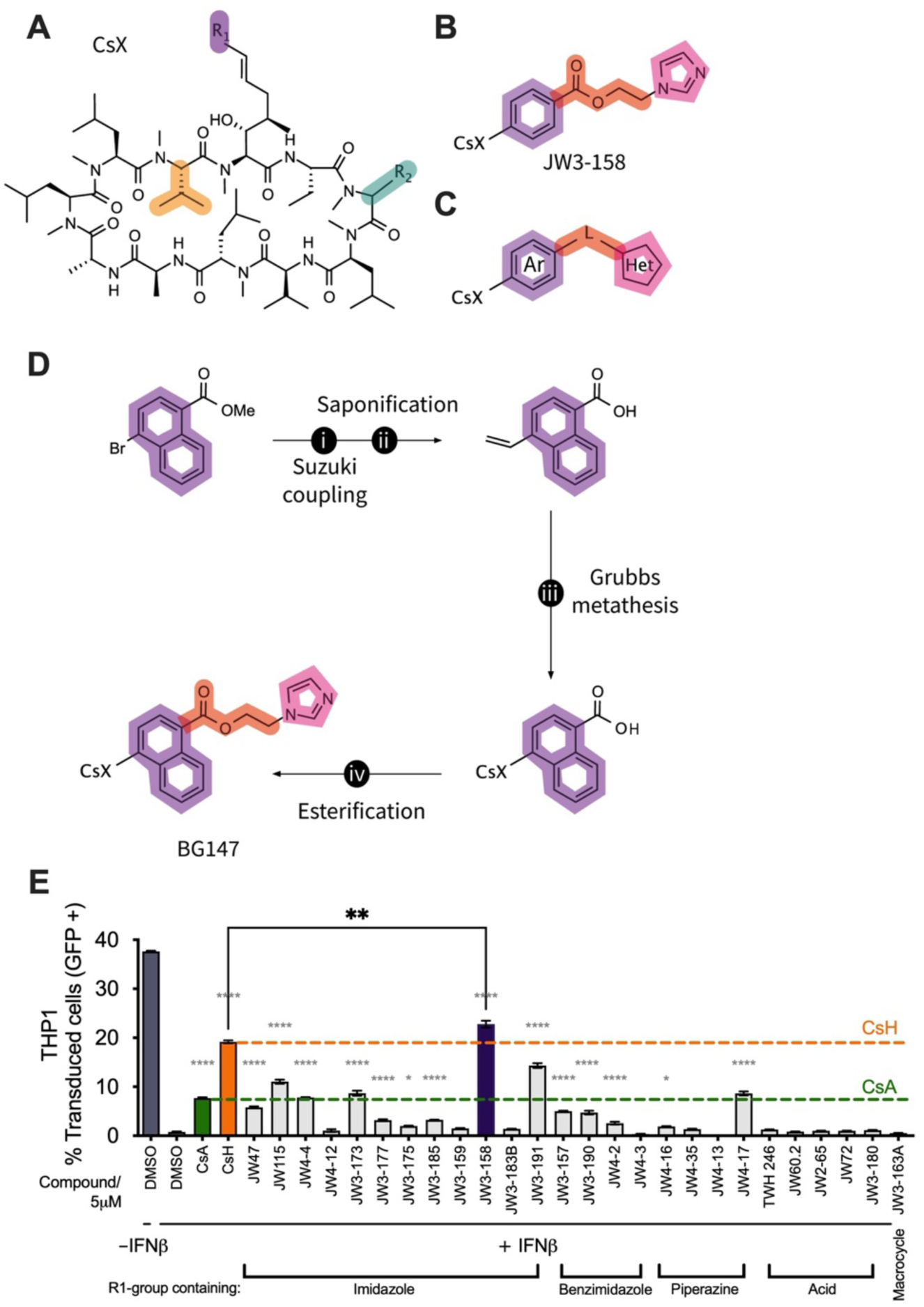
CsA chemical screening library and derivatives. **(A)** Modified Cs structure CsX. CsA, R_1_=CH_3,_ R_2_=H. CsH, R_1_=CH_3,_ R_2_=H, inversion of stereometry at the orange centre, position 11. Modifications exemplified in the screening library (JW series); **(B)** Structure of the prototype molecule JW3-158; **(C)** Variation to the general structure of JW3-158, aromatic group (Ar), linker (L), heterocycle (Het); **(D)** Structure and synthetic route to BG147 : i) K+BF3-CH=CH_2_.Pd[dppf]Cl_2_, Cs_2_CO_3_, THF, water, (89%), ii) LiOH, water, THF (97%), iii) Hoyveda-Grubbs (G2), CH_2_Cl_2_, MW, 70°C, 28%, iv) HATU, DIPEA, DMF(20%). **(E)** LV-GFP infection of THP-1 pretreated with +/-10 ng/ml IFNβ, with 5µM CsA analogues added at the time of infection. % GFP positive cells measured at 48 h post infection (hpi), n=3, +/-SEM. Statistic in black, unpaired t-test, CsH vs JW3-158, statistic in grey ordinary one-way ANOVA with Tukey’s test compared to DMSO + IFNβ.

**Table 1.**
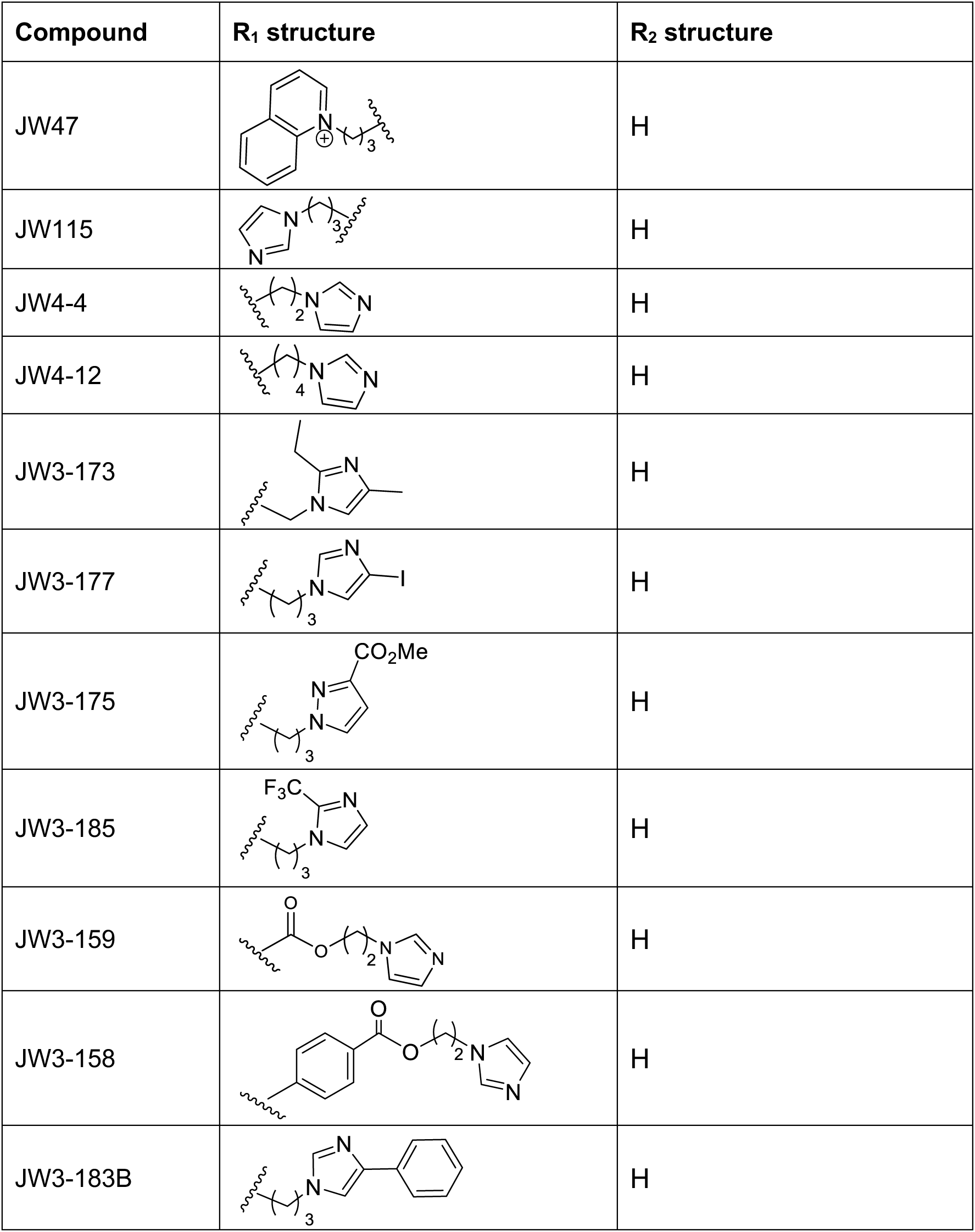
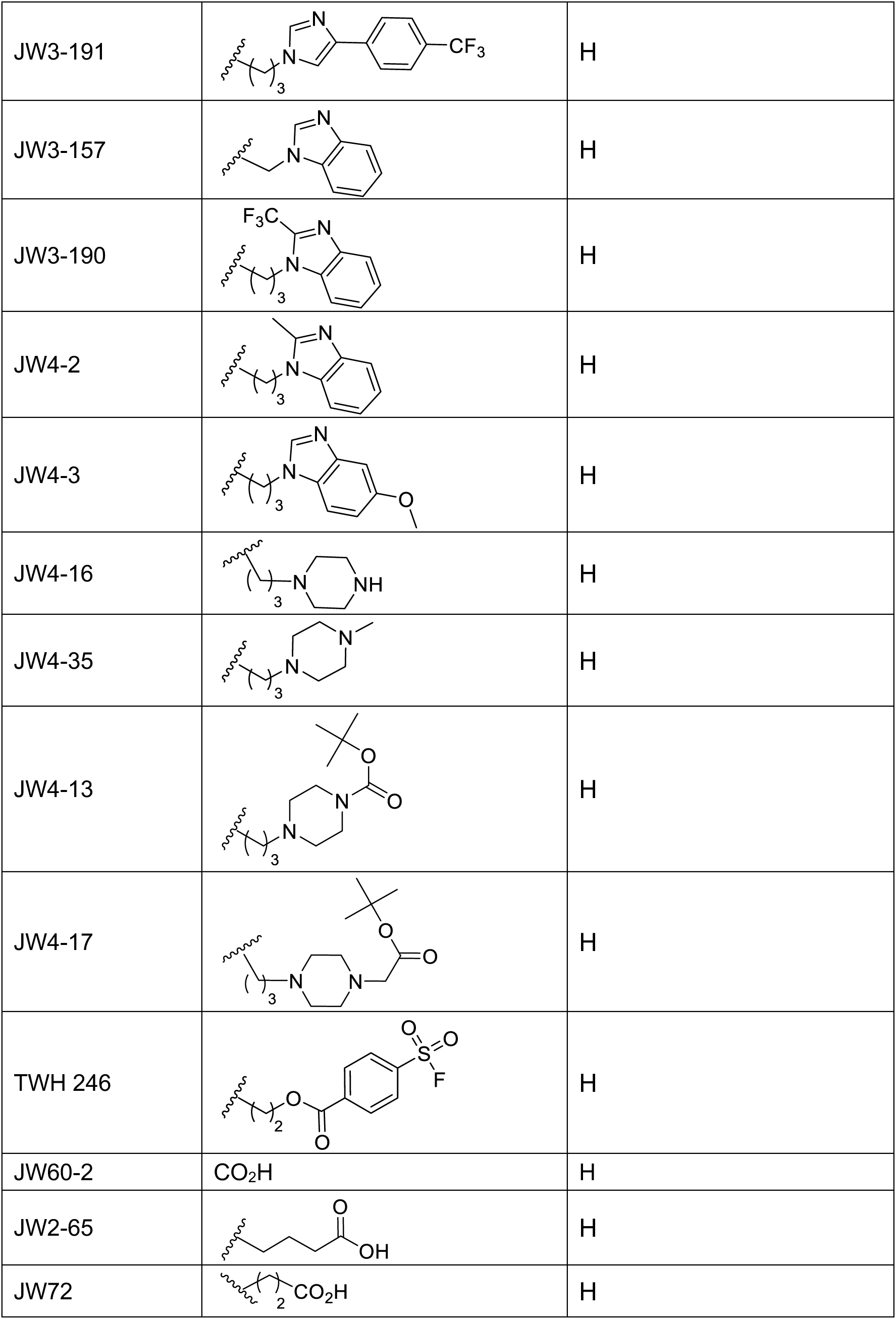
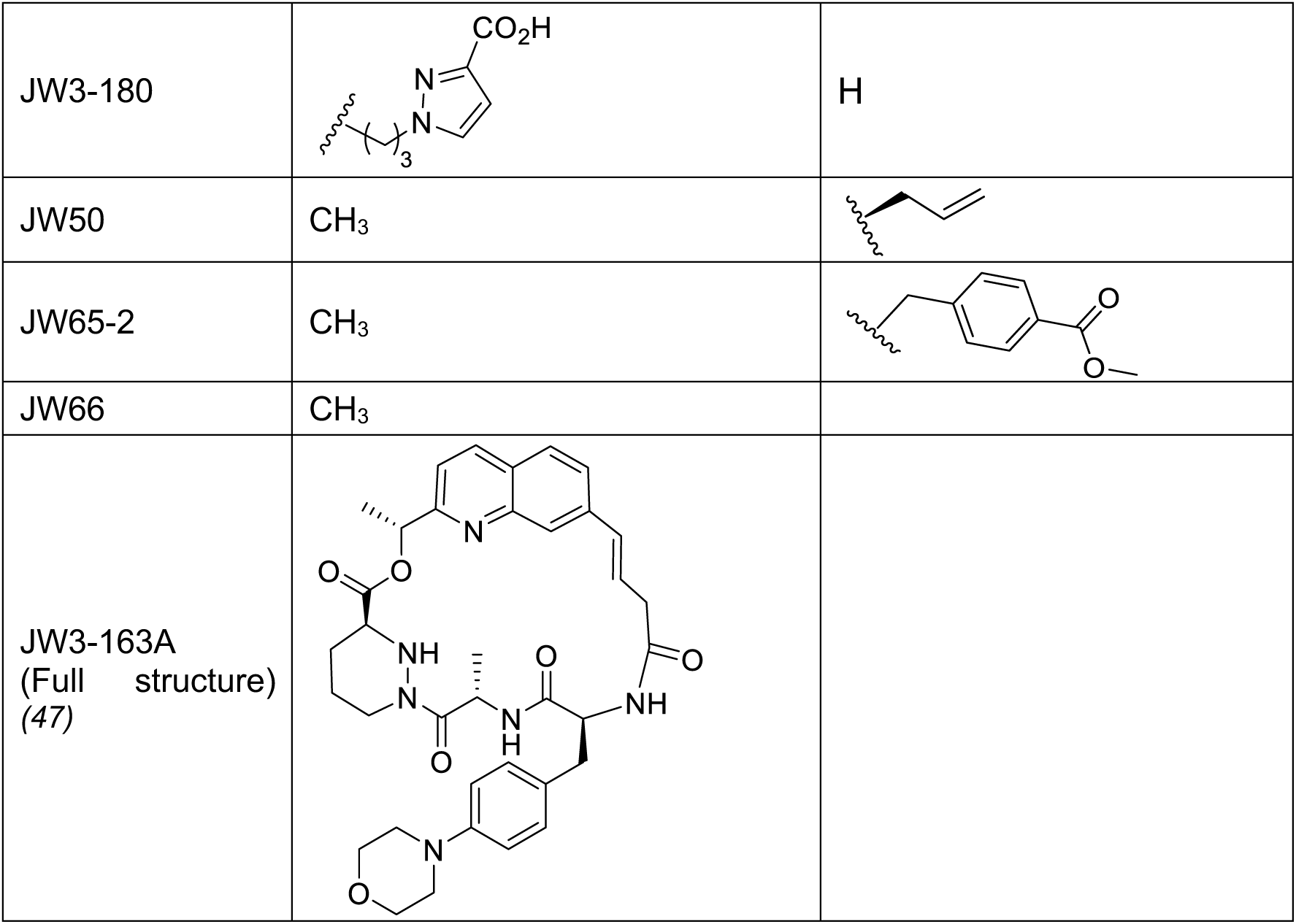
First generation of transduction enhancers (JW series). CsA-R (Fig. 1A).

| Compound | R<sub>1</sub> structure | R<sub>2</sub> structure |
| --- | --- | --- |
| JW47 |  | H |
| JW115 |  | H |
| JW4-4 |  | H |
| JW4-12 |  | H |
| JW3-173 |  | H |
| JW3-177 |  | H |
| JW3-175 |  | H |
| JW3-185 |  | H |
| JW3-159 |  | H |
| JW3-158 |  | H |
| JW3-183B |  | H |
| JW3-191 |  | H |
| JW3-157 |  | H |
| JW3-190 |  | H |
| JW4-2 |  | H |
| JW4-3 |  | H |
| JW4-16 |  | H |
| JW4-35 |  | H |
| JW4-13 |  | H |
| JW4-17 |  | H |
| TWH 246 |  | H |
| JW60-2 | CO<sub>2</sub>H | H |
| JW2-65 |  | H |
| JW72 |  | H |
| JW3-180 |  | H |
| JW50 | CH<sub>3</sub> |  |
| JW65-2 | CH<sub>3</sub> |  |
| JW66 | CH<sub>3</sub> |  |
| JW3-163A<br>(Full structure)<br>(47) |  |  |

We identified the THP-1 monocyte-derived tumour cell line as a model for measuring IFITM3-dependent transduction enhancement because it highly expresses IFITM1-3 upon IFNβ treatment, without expressing other anti-LV factors. We therefore used IFNβ-treated THP-1 to test the CsA analogues for IFITM3 inhibition and transduction enhancement of VSV G-pseudotyped HIV-1 LV encoding GFP, referred to throughout as LV-GFP, measuring GFP expressing cells by flow cytometry (Fig. 1E). The most effective TE, JW3-158, modified at the 1[Bmt] (R1 position, Fig.1A-B), gave better transduction enhancement than CsH at 5µM, with a 29-fold increase compared to DMSO-treated control (Fig. 1E). CsA-SmBz is a previously described non-immunosuppressive CsA derivative with a methylphenyl-4-carboxylic acid group on the 3[Sar] position (R2 position, Fig. 1A)(*44, 45*). CsA-SmBz retains CypA binding and five CsA-SmBz-like analogues were tested but none enhanced transduction to the level of CsA or CsH, consistent with previously reported anti-HIV activity(*45*) (Table 1, Fig. S1B).

### Subhead 2: JW3-158 enhances LV transduction through IFITM3 degradation, but interacts with HIV-1 cofactor CypA

In IFNβ-treated THP-1, JW3-158 decreased IFITM3 levels after 6-24 h treatment (Fig. 2A-B). JW3-158, CsA and CsH showed dose-dependent enhancement of LV transduction in IFNβ-treated THP-1 as expected (Fig. 2C-D). JW3-158 was more potent than CsA and CsH up to 1.25μM (Fig. 2C-D), above which JW3-158 showed some toxicity (Fig. S2A-B). CsA-SmBz did not show transduction enhancement, as expected (Fig. 2C-D). To confirm that transduction enhancement was IFITM3-dependent, THP-1 over-expressing IFITM3 were prepared (Fig. S2C), and over-expression was found to phenocopy the effect of IFNβ treatment with LV-GFP transduction enhanced by JW3-158, CsH and CsA, but not CsA-SmBz (Fig. 2E).

**Fig. 2:**
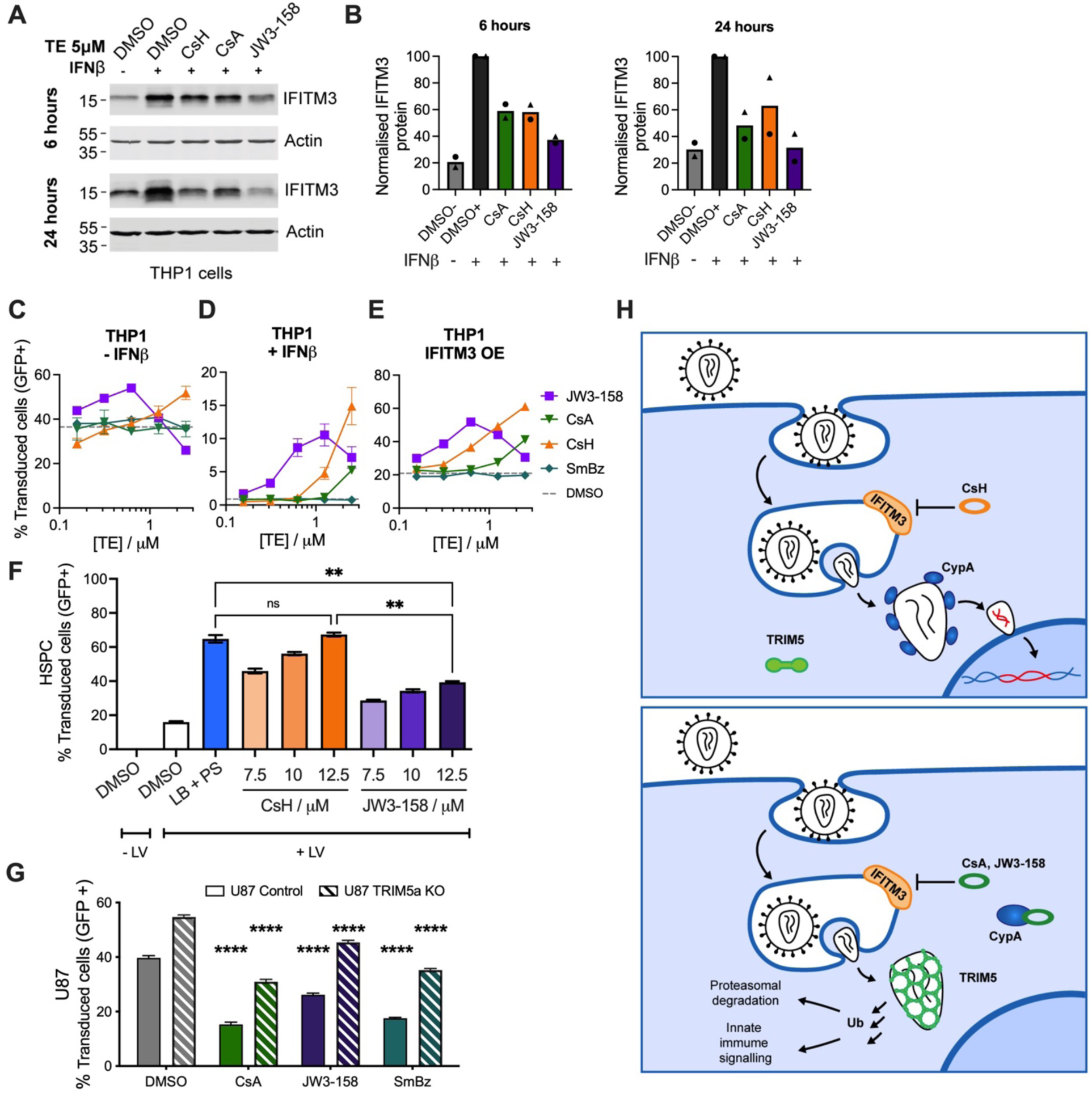
Cyclosporin-like transduction enhancers degrade IFITM3 and inhibit Cyclophilin. **(A)** IFITM3 levels at 6 and 24 h post treatment with 5 µM TEs/DMSO in THP-1 pretreated with +/- 10 ng/ml IFNβ. **(B)** IFITM3 protein quantification from (A), normalised to actin and expressed as a percentage of DMSO + IFNβ. n=2. **(C-E),** Titration of TEs on THP-1 pretreated with +/- 10 ng/ml IFNβ, or in THP-1 over-expressing IFITM3, with LV-GFP for 48 h, n = 3, +/-SEM. **(F)** HSPC treated with 1% LB and 4 μg/ml PS, or novel TEs, with LV-GFP at an MOI of 10 genome copies (GC) /cell, % GFP positive cells measured at 48 hpi, n=2, +/- SEM. One-way ANOVA with Tukey’s test compared to DMSO with LV. **(G)** Unmodified and TRIM5-KO U87 cells treated with 5 μM TEs, with LV-GFP 48 h, n=3, +/- SEM. Statistical significance compared to DMSO (control or TRIM5 KO as appropriate) two-way ANOVA with Tukey’s test. p ≤ 0.0001, ****; p ≤ 0.0002, ***; p ≤ 0.0021, **; p < 0.0332, *; p < 0.1234 ns. **h,** Model of TE mechanism.

We next tested JW3-158 for transduction enhancement in HSPCs. JW3-158 was well-tolerated by HSPCs but failed to increase LV-GFP transduction to the same levels as CsH or established TE combination 1% Lentiboost plus 4 μg/ml protamine sulphate (PS) (Fig. 2F, Fig. S2D). We hypothesised that residual CypA inhibition by JW3-158 caused LV-GFP restriction by TRIM5 and reduced transduction enhancement. qRT-PCR confirmed TRIM5 expression in HSPC (Fig. S2E). Fluorescence polarisation assay detecting CypA-JW3-158 interaction revealed that although JW3-158-CypA binding was reduced as compared to CsA, JW3-158 retained CypA binding activity (CsH IC_50_= undetected, CsA IC_50_ = 37 nM, JW3-158 IC_50_ = 3926 nM, Fig. S1A). The THP-1 used to screen TEs do not restrict TRIM5-sensitive viruses and so to test whether JW3-158 functionally inhibits CypA we used U87 cells. U87 cells allow us to separate TRIM5-dependent inhibition of transduction from IFITM3 dependent inhibition because IFITM3 is not efficiently expressed in unstimulated U87 cells (Fig. S2F). We confirmed TRIM5 activity in unstimulated U87 cells by comparing infection by TRIM5-sensitive N-tropic murine leukaemia virus (MLV-N) with TRIM5-insensitive MLV-B(*46*) (Fig. S2G-H). CsA, JW3-158 and CsA-SmBz treatment all decreased LV-GFP infection in U87, consistent with CypA-sensitive TRIM5 activity, and inhibition reflected the different TE-CypA binding affinities (Fig. 2G, Table 2). Importantly, transduction inhibition by JW3-158 was partially rescued by TRIM5 knock out in U87 (Fig. 2G), confirming a role for TRIM5 in inhibition of LV-GFP transduction by CypA binding CsA analogues. These data underline the importance of avoiding CypA binding and TRIM5 activity for transduction enhancement, as shown in Fig. 2H.

**Table 2.**
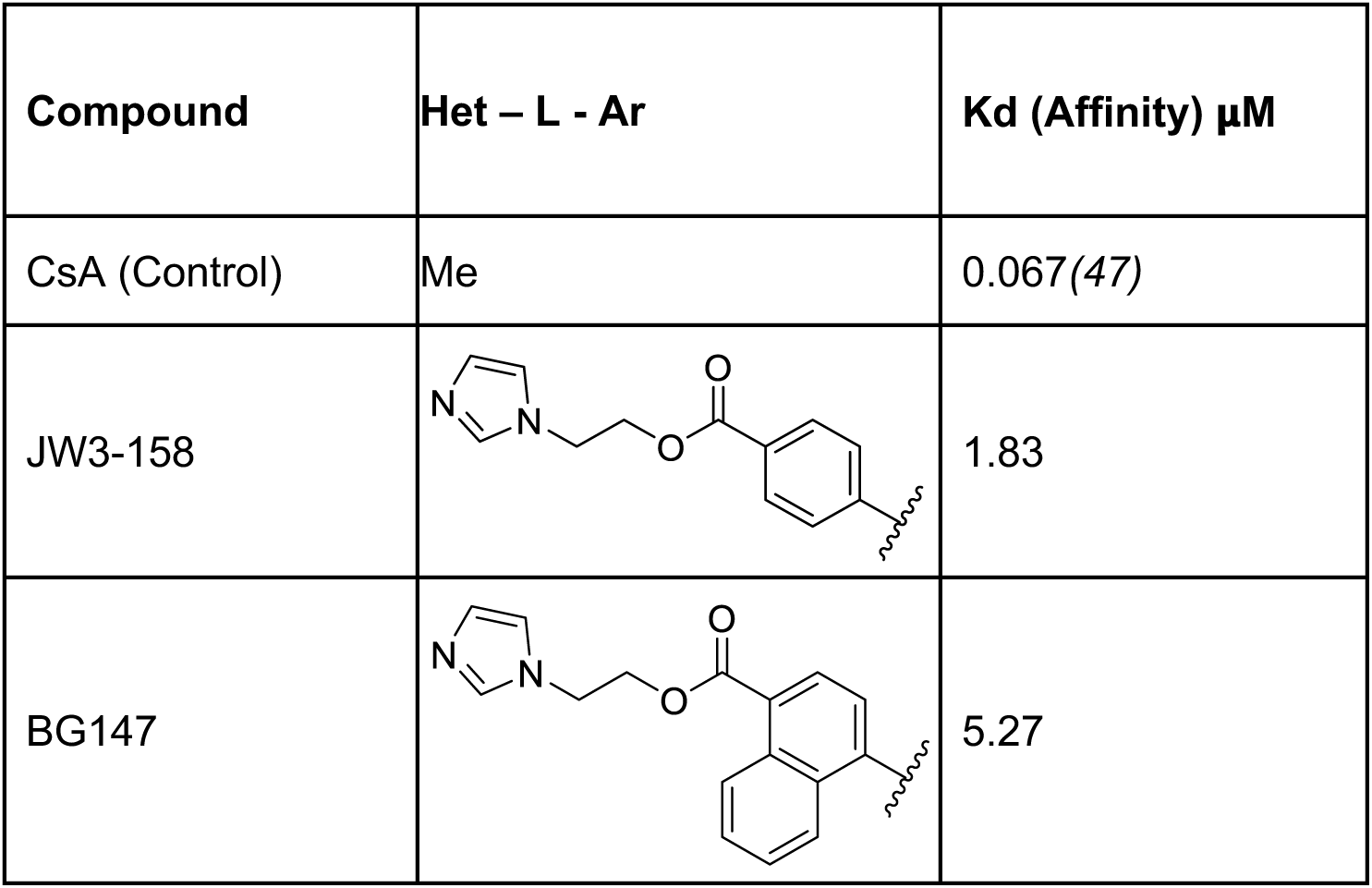
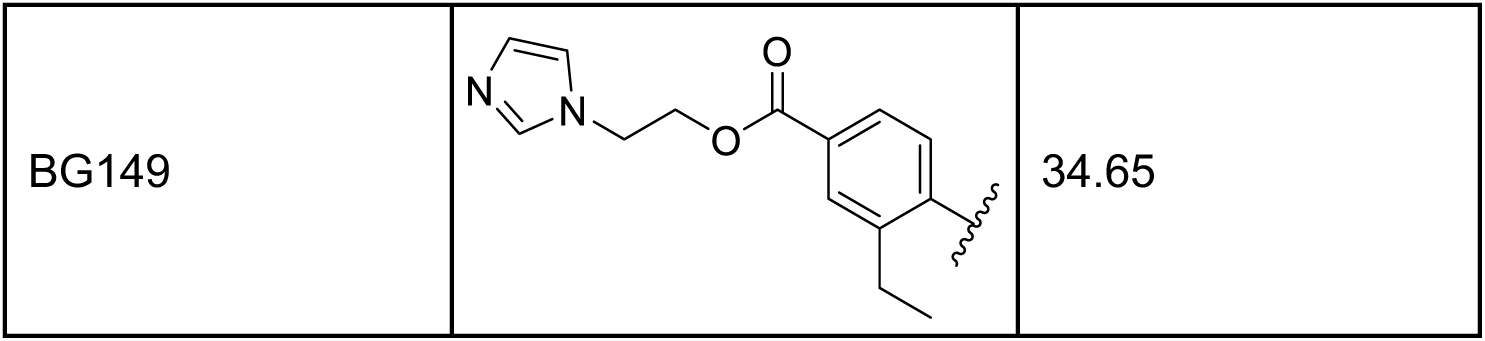
JW3-158 analogues (BG series). Variation to the Ar group of JW3-158 (Fig.1C).

| Compound | Het – L - Ar | Kd (Affinity) μM |
| --- | --- | --- |
| CsA (Control) | Me | 0.067(47) |
| JW3-158 |  | 1.83 |
| BG147 |  | 5.27 |
| BG149 |  | 34.65 |

### Subhead 3: A structure activity relationship (SAR) approach reduced CypA binding and increased transduction efficiency in cell models

Modelling JW3-158 bound to CypA revealed that CypA Ala103 is close to the JW3-158 phenyl group (Fig. 3A). We hypothesised that substituting this aryl group with a bulkier substituent would cause steric clash to reduce CypA binding (Fig. 1B-C). Exploration of substituted aryl groups proved synthetically challenging, but ethyl (BG149) and naphthyl (BG147) analogues were tractable. A typical route is shown for BG147 (Fig. 1D). Initial Suzuki coupling on the bromoaryl **1** and saponification provided the substituted naphthyl acid **2**. Cross-metathesis using the Hovyeda-Grubbs second-generation catalyst provided the CsA-naphthyl acid **3** as a convenient intermediate easily purified from the CsA starting material. Esterification gave the final products. Both analogues displayed reduced CypA binding by surface plasmon resonance (SPR) (CsA Kd = 67 nM(*47*), JW3-158 Kd = 1.8 μM, BG147 Kd = 11.5 μM, BG149 = 34.65 μM, Fig. 3B, Fig. S3A-B, Table 2). Critically, BG147 produced the best transduction enhancement at 5 µM in both IFNβ-treated THP-1 and IFITM3 over-expressing THP-1 (Fig. 3C, E-G). BG147 also retained activity in U87, consistent with it not causing TRIM5 restriction (Fig. 3D). BG147 was well-tolerated in THP-1 and U87 cells up to 5 µM (Fig. S3C-E).

**Figure 3:**
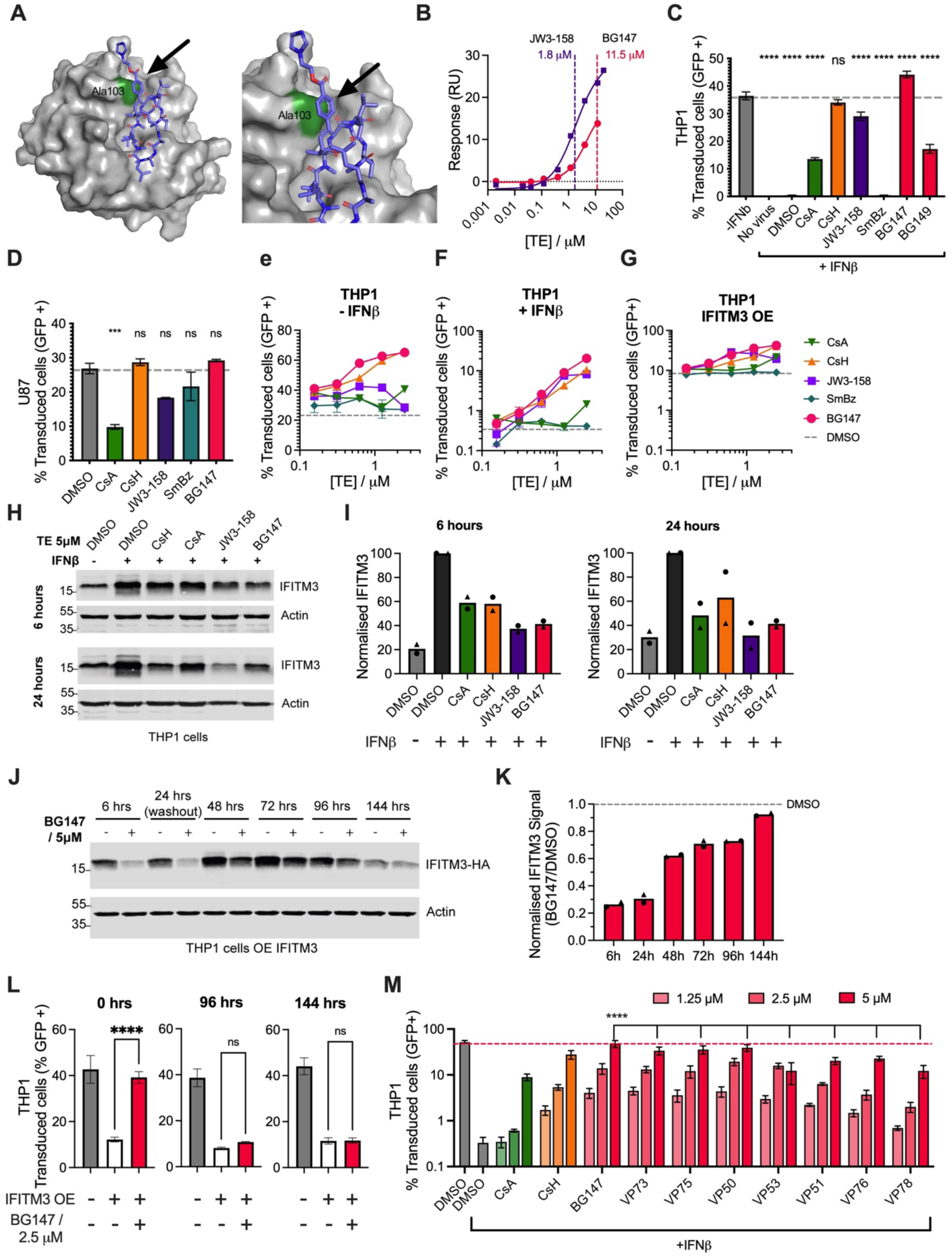
Structure-based design of BG147. **(A)** Model of Cyclophilin A (grey) binding to JW3-158 (blue), showing proximity to residue Ala103 (green). **(B)** Dose-dependent binding (SPR) JW3-158 or BG147 and CypA (immobilised on chip). Steady state response plotted against concentration, (Kd) calculated from the sigmoid fit. **(C-D)** THP-1 cells (pretreated with +/- 10 ng/ml IFNβ) (C) and U87 cells (D) treated with 5μM novel TEs and infected with LV-GFP (48 hpi), n=3, +/-SEM. One-way ANOVA with Tukey’s test compared to -IFNβ (C) or DMSO (D). **(E-G)** Titration of TEs on THP-1 pretreated with +/- 10 ng/ml IFNβ, or in THP-1 over-expressing IFITM3, infected with LV-GFP for 48 h, n = 3, +/-SEM, **(H)** IFITM3 levels at 6 and 24 h post treatment with 5 µM TEs/DMSO in THP-1 pretreated with +/- 10 ng/ml IFNβ. **(I)** IFITM3 protein quantification from (H) normalised to actin and expressed as a percentage of DMSO + IFNβ, n=2. **(J)** THP-1 cells overexpressing IFITM3-HA were treated with 5 μM BG147 or DMSO. After 24 h media was changed, and cells were washed 3 times with PBS. Samples were taken to measure IFITM3-HA levels at 6, 24, 48, 72, 96 and 144 h. **(K)** Quantification of IFITM3-HA levels from (J), normalised to actin and expressed as a percentage of DMSO treatment at each time point, n=3, +/- SEM. **(L)** THP-1 cells overexpressing IFITM3-HA were treated with 5μM BG147 or DMSO for 24 h followed by washout. LV-GFP was added at the same time as BG147 treatment (0 h) or at 96 h or 144 h after treatment, and % GFP cells measured 48 hpi, n=3, +/- SEM. One way ANOVA with Tukey’s test to compare to + IFITM3 - BG147**. (M)** THP-1 cells pretreated with +/- 10 ng/ml IFNβ treated with 1.25, 2.5 or 5 μM TE or BG147 derivatives (VP series) and infected with LV-GFP, infection measured at 48 hpi, n=2, +/-SEM. Two-way ANOVA with Dunnett’s test compared to BG147. p ≤ 0.0001, ****; p ≤ 0.0002, ***; p ≤ 0.0021, **; p < 0.0332, *; p < 0.1234 ns.

Similar to CsA and JW3-158, BG147 led to IFITM3 degradation in IFNβ treated THP-1 at 6 and 24 h (Fig. 3H-I). BG147, JW3-158, CsA and CsH also all led to degradation of exogenously expressed IFITM1 and IFITM2 as well as IFITM3 (Fig. S3F,H). Notably, JW3-158 is better at IFITM1 degradation than BG147, indicating IFITM protein specificity among this SAR series. THP-1 over-expressing IFITM1 or IFITM3 are both reduced in permissivity for LV-GFP infection, but only IFITM3 activity is fully suppressed by 2.5 µM BG147, again consistent with inhibitor specificity (Fig. S3G). To examine the return of IFITM3 after BG147 treatment, we washed THP-1 over-expressing IFITM3 after 24 h treatment and found that IFITM3 levels, and restriction of transduction, substantially returned after 96 h (Fig. 3J-L, Fig. S3I). The short-term effect, and restoration of antiviral IFITM3 activity, suggests that patient cells can be re-infused after functional IFITM3 restoration.

In a final investigation of SAR, we made a series retaining the BG147 naphthalene moiety and replacing the ethyl-imidazole with a variety of groups comprising simple esters, amides and weak bases (Table 3, Fig. 1C). However, none significantly improved transduction enhancement over BG147 in IFNβ treated THP-1 (Fig. 3M, Fig. S3J).

**Table 3.**
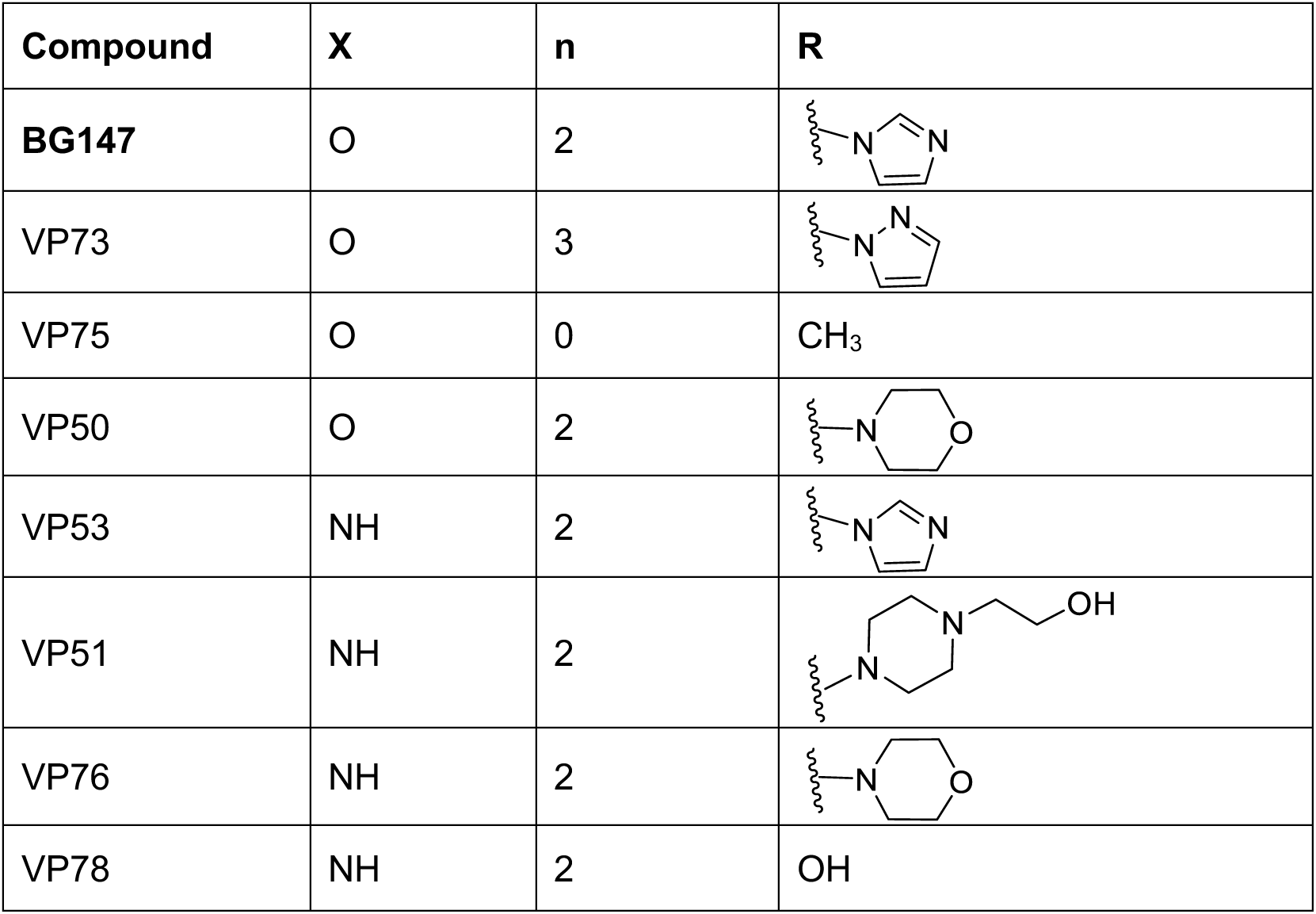
BG147 analogues (VP series). Variation to the L/Het of BG147 (Fig. 1C), R group (Fig. 3K).

| Compound | X | n | R |
| --- | --- | --- | --- |
| BG147 | O | 2 |  |
| VP73 | O | 3 |  |
| VP75 | O | 0 | CH<sub>3</sub> |
| VP50 | O | 2 |  |
| VP53 | NH | 2 |  |
| VP51 | NH | 2 |  |
| VP76 | NH | 2 |  |
| VP78 | NH | 2 | OH |

### Subhead 4: BG147 enhances HSPC transduction more effectively than CsH and comparably to classical TEs PS and Lentiboost

We next showed that BG147 significantly enhanced HSPC transduction compared to JW3-158, achieving transduction levels comparable to CsH but at a lower dose (2.5 μM BG147 compared to 7.5 μM CsH) (Fig. 4A). Higher concentrations of BG147 were not tested due to toxicity at >5 µM in THP-1 (Fig. S3D, S4A). BG147 boosted transduction similarly to gold-standard TE Lentiboost (1%) + PS (4 μg/ml) (Fig. 4A) but combining BG147 with Lentiboost and PS gave no significant additive effect (Fig. 4A).

**Fig. 4:**
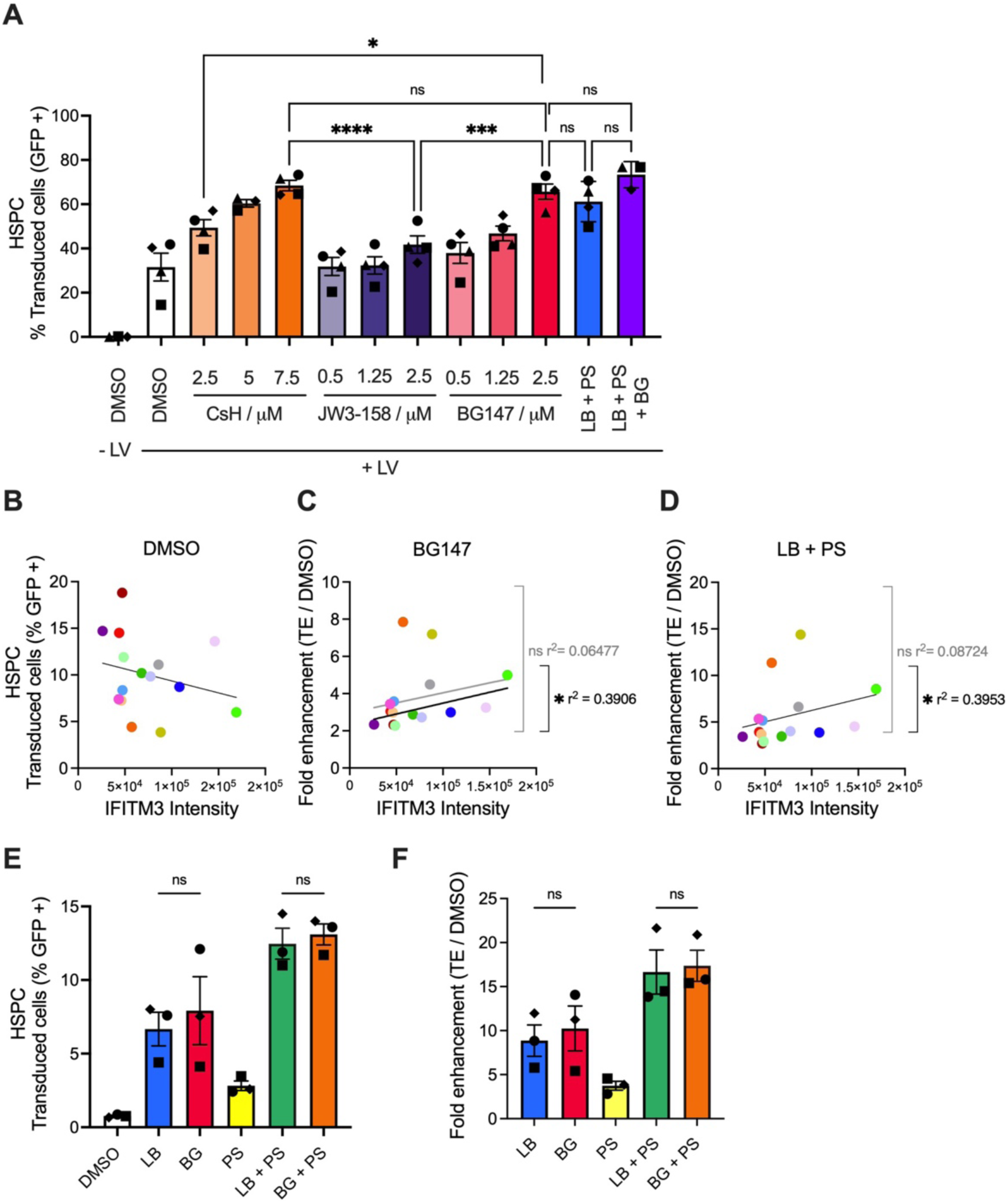
BG147 is novel, efficient transduction enhancer in HSPCs. **(A)** HSPC treated with 1% LB and 4 μg/ml PS, or novel TEs as shown, and transduced with LV-GFP (LV) at an MOI of 10 GC/cell, % GFP positive cells measured at 48 hours post infection (hpi), n=4 HSPC donors, represented by different symbols, +/- SEM. 2-way ANOVA with Tukey’s test. **(B-D)** HSPCs treated with 1% LB and 4 μg/ml PS or 2.5 μM BG147 were transduced with LV-GFP (LV) at an MOI of 2.5 IU/cell, % GFP positive cells measured at 9 dpi. Correlation of IFITM3 protein levels in donors with base level transduction (B), fold enhancement in transduction with 2.5μM BG147 (C), or 1% LB, 4 μg/ml PS (D). IFITM3 levels were measured by WB for each donor, normalized against actin. Simple linear regression, Pearson correlations. n=14 HSPC donors, represented by different coloured symbols. **(E-F)** HSPCs treated with 1% LB, 4 μg/ml PS, 2.5 μM BG147 (BG), or a combination, were transduced with LV-GFP (LV) at an MOI of 2.5 IU/cell, % GFP positive cells measured at 9 dpi (E) and fold enhancement over DMSO calculated (F). n=3 HSPC donors. One way ANOVA. p ≤ 0.0001, ****; p ≤ 0.0002, ***; p ≤ 0.0021, **; p < 0.0332, *; p <0.1234 ns.

Transduction variability between donors is linked to IFITM3 levels(*18*) so we next obtained HSPC from 15 donors and compared transduction enhancement using a lower MOI (2.5 IU/cell), measuring IFITM3 levels by immunoblot. BG147 gave an average of 3.75-fold enhancement, which was lower than LB + PS (5.59-fold) (Fig. 4B-D). Tracking donors across treatments revealed a striking similarity in sensitivity to BG147 and to LB + PS, suggesting related mechanisms, see Figure 5. IFITM3 levels trended negatively with baseline transduction, and positively with fold-enhancement (Fig. 4B-D, Fig. S5A). However, outliers confounded a statistically supported relationship suggesting factors beyond IFITM3 contribute to donor variability. Notably, among donors with 2 to 6-fold enhancement (13/15 donors) there was a Pearson significant correlation (r^2^=0.3906). There was no statistically supported correlation between IFITM3 mRNA expression (qRT-PCR) and baseline transduction or transduction enhancement (Fig. S5B-D). Seeking further explanation for transduction variability, we detected ISGs IFITM1, tetherin and MxB by immunoblot. IFITM1 suppresses VSV-G pseudotype infection and is affected by BG147 (Fig. S3G-H) and MxB suppresses LV nuclear transport(*48*). Both were expressed variably across donors and neither correlated with outliers in the transduction plot confounding a simple explanation, for example, interferon exposure of poorly transduced donor cells (Fig. S4E-F, H-I). Tetherin is known to be expressed in HSPCs, and although we did not expect it to restrict incoming LV(*49*) we were surprised to find that high tetherin protein levels correlated significantly with high baseline transduction levels (Fig. S4G). There was no correlation between tetherin and BG147 induced transduction enhancement (Fig. S4J).

**Fig. 5:**
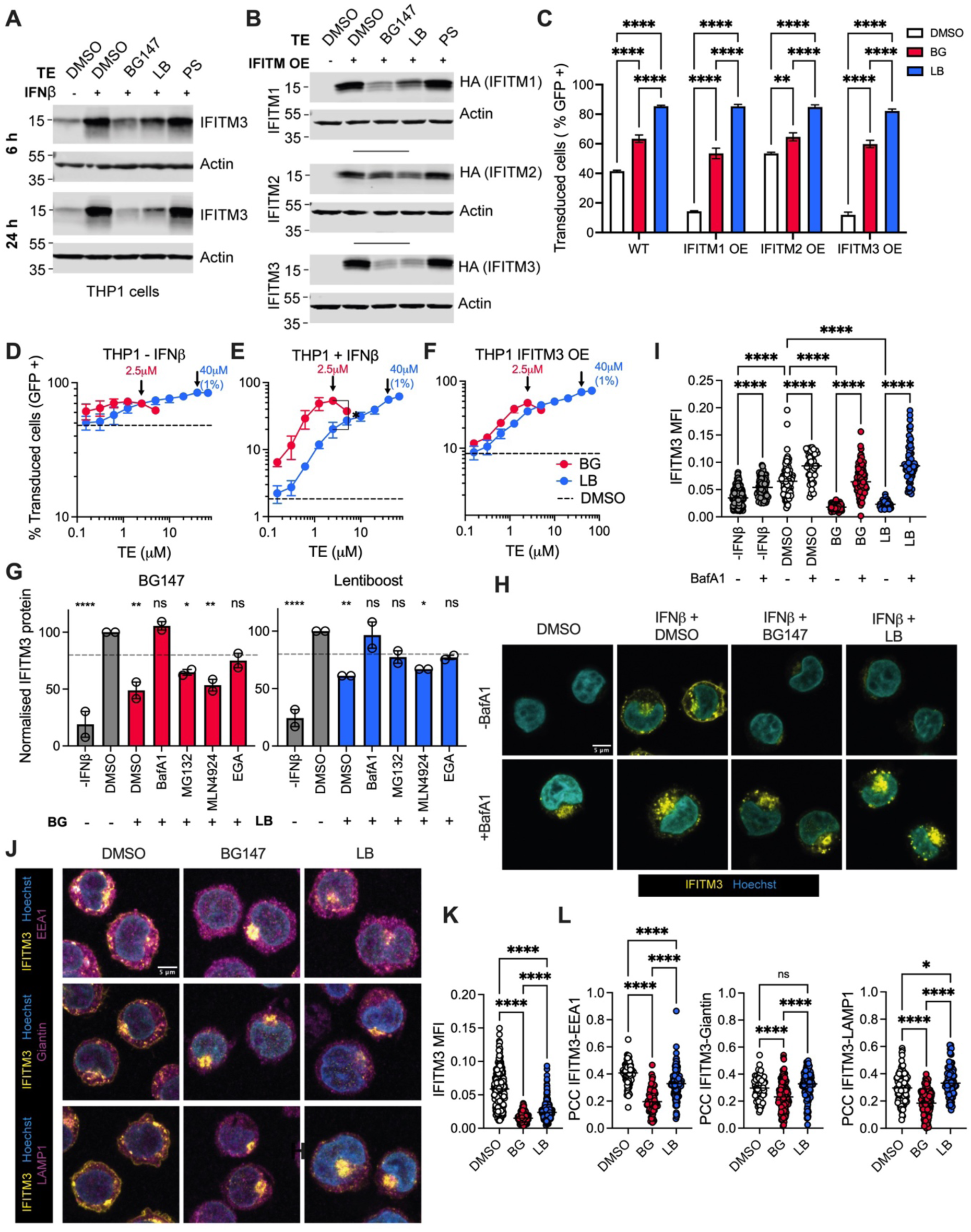
BG147 and Lentiboost cause IFITM3 depletion through aberrant trafficking. **(A)** IFITM3 levels at 6 and 24 h post treatment with 1% Lentiboost (LB) and 4 μg/ml PS, 2.5 μM BG147 (BG), or a combination, in THP-1 pretreated with +/- 10 ng/ml IFNβ. n=2. **(B)** Levels of IFITMs in THP-1s overexpressing IFITM1-HA, IFITM2-HA or IFITM3-HA treated with 1% LB and 4 μg/ml PS, 2.5 μM BG147 (BG), or a combination, for 24 h. **(C)** THP-1 cells overexpressing IFITM1-HA, IFITM2-HA or IFITM3-HA treated with 1% LB, 2.5μM BG147 or DMSO at the time of infection with LV-GFP. % GFP positive cells measured at 48 hpi, +/- SEM, 3 replicates, n=2. Statistical significances compared to DMSO was determined by two-way ANOVA. **(D-F)** Titration of LB and BG on THP-1 pretreated with +/- 10 ng/ml IFNβ, or in THP-1 over-expressing IFITM3, infected with LV-GFP for 48 h, +/-SEM 3 replicates, n = 2. (E) Paired t test. p ≤ 0.0001, ****; p ≤ 0.0002, ***; p ≤ 0.0021, **; p < 0.0332, *; p <0.1234 ns. **(G)** IFITM3 levels at 6 h post treatment with 1% LB or 2.5 μM BG147 (BG), and cellular process inhibitors (MG132 proteasome inhibitor 20 µM, MLN4924 NEDDylation inhibitor 1 µM, BafA1 lysosome inhibitor 0.5 µM, EGA late endosome trafficking inhibitor 20 µM) in THP-1 pretreated with +/- 10 ng/ml IFNβ. (Fig. S7A). IFITM3 protein density normalised to actin and plotted relative to THP-1 + IFNβ + DMSO, n=2. Two-way ANOVA compared to THP-1 +IFNβ + DMSO. **(H)** IFITM3 levels measured by immunofluorescence at 6 h post treatment with DMSO, 1% LB or 2.5 μM BG147 (BG) and with and without BafA1 (0.5 µM) in THP-1 pretreated with +/- 10 ng/ml IFNβ. N=2. **(I)** Mean fluorescence intensity (MFI) of IFITM3 from experiment shown in (D). Ordinary one-way ANOVA. N=2**. (J)** IFITM3 levels measured by immunofluorescence at 6 h post treatment with DMSO, 1% LB or 2.5 μM BG147 (BG) in THP-1 pretreated with +/- 10 ng/ml IFNβ. Markers used for Golgi (anti-Giantin), early endosome (anti-EEA1) and lysosome (anti-LAMP1). N=2. **(K)** Mean fluorescence intensity (MFI) of IFITM3 from pooled samples from experiment shown in (J). **(L)** The Pearson correlation coefficient (PCC) was calculated from (J) to quantify the colocalisation of IFITM3 with EEA1, Giantin or LAMP1. N=2. One-way ANOVA Ordinary one-way ANOVA. N=2.: p ≤ 0.0001, ****; p ≤ 0.0002, ***; p ≤ 0.0021, **; p < 0.0332, *; p < 0.1234 ns.

Previous work has shown that PS boosts transduction enhancement by Lentiboost(*50*). We found that Lentiboost and BG147 enhanced transduction similarly when used alone and that PS boosted each similarly (Fig. 4E-F).

### Subhead 5: Lentiboost causes IFITM degradation

Next, we tested whether Lentiboost or PS cause IFITM3 degradation. Immunoblots revealed that Lentiboost induced degradation of endogenous IFITM3 in IFNβ–treated THP-1 at 6 and 24 h after treatment (Fig. 5A, Fig. S6A-B). 1% Lentiboost was less effective than 2.5μM BG147 and PS had no effect on IFITM3 levels. IFITM2 levels were also reduced by Lentiboost although not as effectively as by BG147, and again PS had no effect (Fig. S5C). IFITM1 was not detected in IFNβ-treated THP-1 (Fig. S5D). In THP-1 over-expressing IFITM1, 2 or 3, Lentiboost and BG147 also induced degradation of each IFITM and PS had no effect (Fig. 5B-C, Fig. S6E-F). Over-expression of IFITM1 and 3, but not IFITM2, in THP-1 restricts LV-GFP (Fig. 5C). Like BG147, 1% Lentiboost enhanced transduction in each of the IFITM expressing THP-1 lines, as well as in the unmodified THP-1 (Fig. 5C). The modest enhancement observed in unmodified and IFITM2-expressing cells is likely due to degradation of basal endogenous IFITM3, although Lentiboost could have additional transduction enhancing effects. We next assessed potency of the TEs in THP-1 treated with IFNβ, or over-expressing IFITM3, with a fixed dose of LV. We found that both BG147 and Lentiboost depend on IFITM3 expression for transduction enhancement (Fig. 5D-F). Furthermore, BG147 was significantly more potent than Lentiboost in IFNβ-treated THP-1, although the effect of BG147 plateaus before Lentiboost (Fig. 5E). 2.5 μM BG147 has activity comparable to 1% Lentiboost (the clinically relevant concentration, approximately 40 μM) and 2.5 μM Lentiboost had significantly lower activity than 2.5 μM BG147 (Fig. 5E). Lentiboost enhancement is biphasic (Fig. 5D-F), suggesting additional activities. PS did not enhance transduction in THP-1 (Fig. S6H). We conclude that transduction enhancement by Lentiboost is mediated in part through the inhibition of IFITM3-mediated restriction, similar to BG147 and other TE(*17–19, 51*).

### Subhead 6: Transduction enhancers cause IFITM relocalisation and degradation via lysosomes

To investigate how BG147 (and Lentiboost) cause IFITM3 degradation, we inhibited cellular degradation pathways and measured the effect of TEs on IFITM3 levels in IFNβ-treated THP-1. Inhibition of lysosome acidification and autophagy using Bafilomycin A1 (BafA1) blocked depletion of IFITM3 by both TEs (Fig. 5G, Fig. S6A-C). Inhibition of proteasomes with MG132 or MLN4924 (NEDD8-activating E1 enzyme inhibitor), or late endosome trafficking with EGA did not rescue IFITM3 degradation (Fig. 5G, Fig. S7A-C). These data suggest that IFITM3 is depleted due to TE-dependent trafficking to lysosomes. This was supported by immunofluorescence microscopy. IFITM3 levels decreased after 6 h TE-treatment and BafA1 disrupted the altered IFITM3 trafficking and rescued it from degradation (Fig. 5H-I, Fig. S7D-F). In untreated cells, endogenous IFITM3 colocalizes with cellular marker EEA1 (early endosomes) and to a lesser extent Giantin (Golgi) and LAMP1 (lysosomes) (Fig. 5J,L). After 6h treatment, BG147 caused greater loss of IFITM3 than Lentiboost (Fig. 5J-K, Fig. S7G-I). Both TEs decreased IFITM localisation with EEA1, a small increase in LAMP1 colocalisation with Lentiboost supports a slower/less complete degradation (Fig. 5J-L, Fig. S7G-I). In TE-treated samples, IFITM3 localised to nuclear-proximal spherical bodies, possibly the trans-Golgi network (TGN) or ER-Golgi intermediate compartment (ERGIC) transport hubs (Fig. 5J). These results are similar to those seen with rapalog TEs (*17, 51*) and support TE-driven IFITM3 degradation dependent on lysosomal acidification.

### Subhead 7: BG147 as a transduction enhancer in *ex vivo* and *in vivo* gene therapy

In further optimisation experiments, we found that BG147-mediated transduction enhancement is boosted by a 24 h pretreatment facilitating 80% transduction of IFN-treated THP-1 cells (Fig. 6A-B). Conversely, Lentiboost displayed no increase in activity on pretreatment.

**Figure 6:**
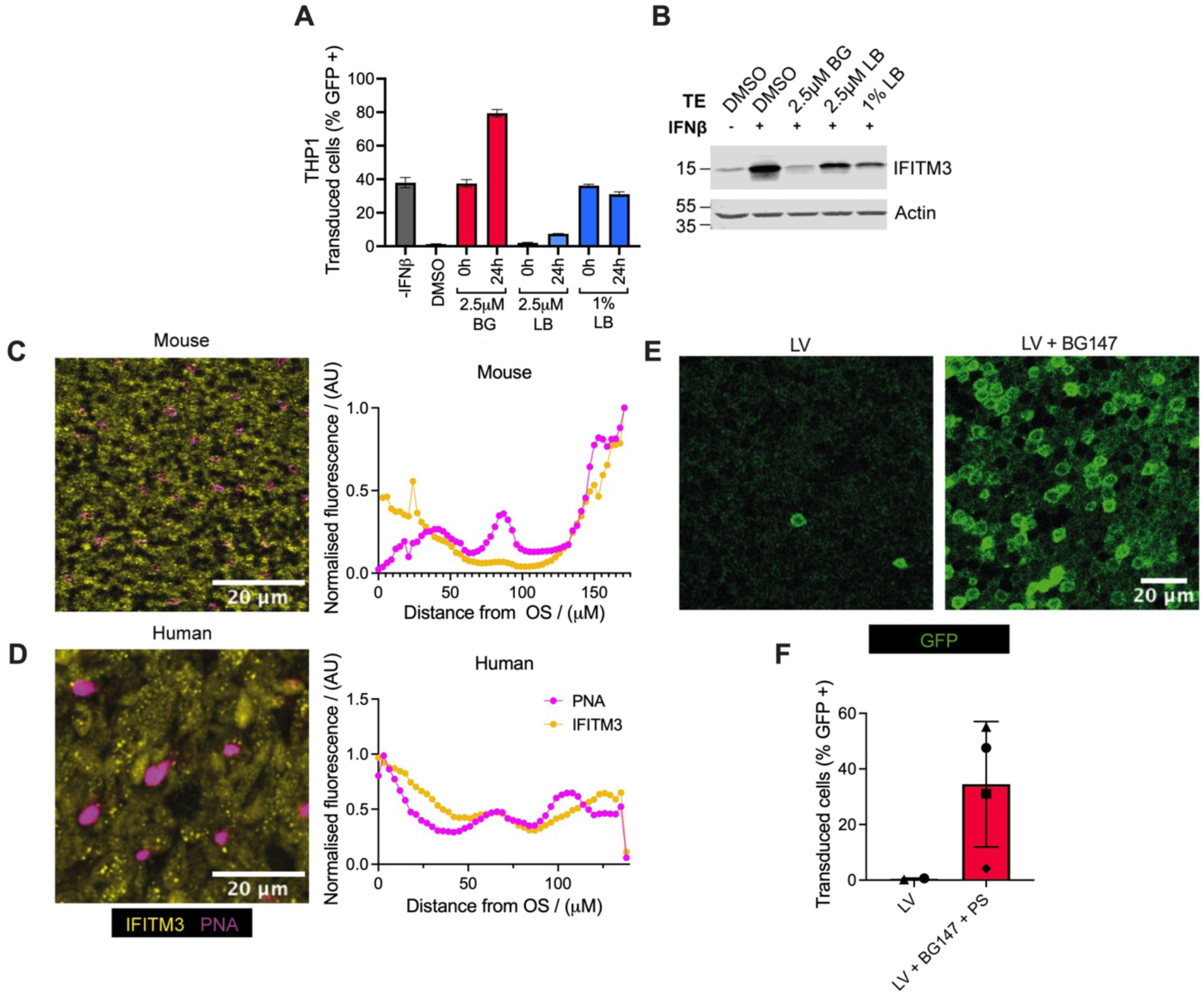
In vivo transduction enhancement. **(A-B)** THP-1s were pretreated with 1% LB or 2.5 μM BG147 (BG) for 24 or 0 h then infected with CMV-GFP (LV) and at an MOI of 2.5 IU/cell with fresh 1% LB or 2.5 μM BG, % GFP positive cells measured at 48 hpi (A), samples were taken for WB at 24 h (B), n=2. **(C-D)** Confocal microscopy of IFITM3 levels (yellow) in the photoreceptor segments of mouse (C) and human (D) retina, represented by the distance from the outer segment (OS), from ganglion to photoreceptor cells. Photoreceptor cone cells are detected by Peanut agglutinin lectin (PNA) marker (pink). **(E)** CMV-GFP (2 μl) with or without BG147 (0.1 μl, 100 μM) was administered to mice by subretinal injection. 1 month post injection wholemount retina was imaged by confocal microscopy. N=4, Representative images from 1 mouse. **(F)** Quantification of GFP signal from (E).

Finally, we investigated enhancement of *in vivo* transduction, using murine ocular gene delivery as a model. LVs work poorly in photoreceptor cells when injected into the subretinal space (*9–15*), to deliver ABCA4 to treat Stargardt disease(*52*)and Lentiboost is unsuitable *in vivo* due to the high concentrations required. We first used immunofluorescent labelling and confocal microscopy of inner and outer segments of both mouse and human retinal samples to confirm IFITM3 expression in photoreceptor cells (Fig. 6C-D). IFITM1 and 2 were also detected (Fig. S8A-B).

To assess whether BG147 enhances LV transduction *in vivo*, we co-administered a VSV-G CMV-GFP (CMV-GFP) and BG147 to mice via sub-retinal injection. Photoreceptor transduction, assessed one month after injection by counting GFP expressing cells, was increased from less than 0.5% (with CMV-GFP alone), to an average of 34.5% with BG147 treatment, with the maximum transduction being 55.0% (Fig. 6E,F). Thus, BG147 facilitates efficient LV transduction in ocular *in vivo* gene therapy. This finding is transformative as it overcomes a major hurdle for the deployment of integrating, well-defined and safe LV gene delivery systems for *in vivo* photoreceptor transduction and, suggests that BG147 may be invaluable in additional *in vivo* gene therapy protocols.

## DISCUSSION

We have developed a novel cyclosporine A analogue, BG147, optimised via structure-guided design to enhance LV transduction *ex vivo* and *in vivo* by targeting IFITM3 for degradation. Autologous stem cell gene therapy for genetic diseases, often curative with a single treatment, promises revolutionary treatment improvements in almost all branches of medicine. Clinical trials using VSV-G pseudotyped vectors have demonstrated benefits in diverse disorders, including SCID-X1, SCID-ADA, CGD, Wiskott-Aldrich syndrome, leukocyte adhesion deficiency, and several metabolic conditions(*1, 3–5*). Advantages of LV over, for example, AAV–based vectors include large capacity, stable integration that avoids epigenetic silencing and broad cell tropism. Non-integrating LVs can also deliver Cas9 systems or larger donor templates used in genomic correction(*53*). However, high production cost remains a significant limitation.

Natural products, like CsA, are evolved molecules typically with complex activities. We took an SAR approach to reduce unwanted properties (CypA binding inhibits HIV infection, CN inhibition disrupts T-cell function) and retain desirable properties (transduction enhancement). We showed that BG147 drives IFITM3 relocalisation and lysosome-dependent degradation, evidenced by IFITM3 rescue by BafA1, but not proteasome inhibitors (Fig. 6). The weak base imidazole group on BG147 may help target it to acidic endosomal compartments (Fig. 1). IFITM3 dependence and BafA1 sensitivity has been reported for other TEs including caraphenol (resveratrol trimer), rapamycin (and analogues), implicating IFITM3 as a central barrier to LV transduction(*17–19, 51*). The TE Lentiboost, widely used in clinical studies. was thought to act by promoting virus attachment (*39, 40*) or cell permeability(*54*). Here we show that Lentiboost drives IFITM3 degradation and rescues THP-1 transduction in an IFITM3-dependent manner (Fig. 5, Fig. 6). Lentiboost is a polyaxamer, used at high concentrations (1% is approximately 40 μM), which may affect cell membrane biology. BG147, used at 2.5 μM, is expected to target IFITM trafficking more specifically.

The molecular target of BG147, and the mechanism by which IFITMs are targeted for degradation, remain unclear. BG147 may interact with IFITMs directly, disrupting, for example, oligomerization or cholesterol binding. IFITM3 may inhibit viral fusion by disturbing cholesterol homeostasis to increase endosomal cholesterol(*23*) thus inhibition of IFITM3-cholesterol interactions could impede function. Recombinant IFITM3 binds to cholesterol through residues crucial for antiviral activity(*55*), however, neither artificially increasing cholesterol levels, nor cholesterol-sequestration, affected influenza A fusion in the presence of IFITM3(*24, 56, 57*). Conflicting findings may be due to over-expression of IFITMs, which alters localisation, and impacts antiviral activity(*58*).

Donor-to-donor variation in transduction efficiency during HSPC gene therapy remains problematic in clinic. High levels of IFITM3 have been associated with low transduction and large CsH-mediated enhancement(*18*). Although we also saw correlation between IFITM3 expression and BG147 mediated enhancement, outliers in which transduction was not rescued by BG147, or high transduction enhancement despite low IFITM3 levels, confounded a simple relationship (Fig. 4E). Surprisingly, levels of the ISG tetherin correlated with high base-line transduction, suggesting that regulation of LV permissivity in HSPCs is multifactorial (Fig. S4I). We envisage universally effective HSPC gene therapy through personalised TE cocktails, guided by proteomic or RNAseq profiling of HSPC targets, and thus inhibiting patient-specific combinations of anti-LV restriction factors. This will require identification of the restriction factors involved and specific mitigations. It will also be interesting to investigate whether synthetic lipid nanoparticle (LNP) systems benefit from TE as we expect they will.

BG147 is derived from CsA, a widely used and safe drug with over 2 million prescriptions in the US in 2022(*59*). It is well tolerated by HSPC (Fig. S4A) and we do not expect additional adverse effects because our SAR reduced activity against CypA and CN whilst retaining natural anti-IFITM activity. *Ex vivo* gene therapy reduces TE toxicity because TE can be washed off before re-infusion of modified cells. Indeed, we found that IFITM3 function returns after 96h (Fig. 3K, Fig. S3I-K).

The future of gene therapy lies in *in vivo* approaches which circumvent lengthy, expensive *ex vivo* protocols making therapies more accessible. TE are likely to play an essential role. To evidence this, we evaluated the effect of BG147 on *in vivo* LV transduction of mouse photoreceptor cells after sub-retinal injection. We showed that the photoreceptor layer in both mouse and human retina expresses IFITM3 (Fig. 6C-D), likely explaining reported poor LV transduction efficiency(*9–15*). Co-administration of BG147 with LV boosted transduction efficiency by 70-fold in our model (Figure 6).

In summary, we have synthesised and characterised a novel CsA-based TE, BG147, that enhances *ex vivo* HSPC transduction and *in vivo* mouse photoreceptor transduction by LV using clinical protocols. Our work supports previous reports that CsA-based TEs interfere with IFITM localisation/trafficking leading to enhanced transduction. We report for the first time that the commonly used TE, Lentiboost, also predominantly acts via this mechanism, informing its use in combination treatments. More broadly, our work brings LV, with their desirable characteristics of high capacity, low immunogenicity and stable integration, back to the forefront as viable *in vivo* delivery options, applicable to other emerging gene therapy protocols. This work illustrates how a sound understanding of host-virus interactions and infected-cell biology can drive therapeutic innovation.

## MATERIALS AND METHODS

### Study design

The aim of this study was to design and evaluate a synthetic cyclosporine-based molecule for the enhancement of LV transduction in HSPCs. We used THP-1 cells stimulated with IFNβ to produce ISGs such as IFITM3, as a model for HSPCs which express ISGs, including IFITM3, constitutively. U87 cells were used as a model to investigate interactions with CypA and TRIM5, which are not active in THP-1s. In THP-1 and U87 cells, WB was used to assess IFITM3 levels and flow cytometry to measure LV transduction of HIV-GFP. Each experiment was done at least twice on different days. For each flow cytometry experiment 3 technical replicates were done in parallel, derived from the same batch of cells. Statistical analyses were performed using GraphPad Prism (v10.3.1) as indicated in the figure captions.

### Tissue origin, Animal handling and ethics

For HSPC experiments donor cells were obtained from Lonza Biologics as mobilized CD34+ cells (#4Y-101D), or via isolation of CD34+ cells from a Peripheral Blood Stem Cell (PBSC) Donation courtesy of Anthony Nolan Charity, UK, following the method outlined in(*60*).

All animal procedures were conducted in accordance with the UK Animals (Scientific Procedures) Act 1986 and approved by the UK Home Office under Project Licence PPL number PP2185067. Experiments adhered to institutional guidelines for the care and use of laboratory animals and followed the Association for Research in Vision and Ophthalmology (ARVO) Statement for the Use of Animals in Ophthalmic and Vision Research (Rockville, MD, USA).

Human retinal tissue was obtained from the Moorfields Eye Hospital Biobank (London, UK) under the approval of the Moorfields Biobank governance framework (REC reference: 20/SW/0031; IRAS Project ID: 352490). All tissue was collected with informed consent from donors in accordance with the Declaration of Helsinki. The research was conducted in compliance with UK ethical and regulatory standards for research involving human tissue, and the Moorfields Biobank operates under a licence from the UK Human Tissue Authority (HTA).

### Chemical synthesis and compound characterisation

Descriptions of methods of chemical synthesis and characterisation are provided in the Supplementary Information. Compounds were dissolved in anhydrous DMSO (900645, Sigma Aldrich) for use in biological assays and stored in screw-top Eppendorf tubes at −20 °C with drying agent.

### Surface plasmon resonance (SPR) against CypA

SPR experiments were performed using Biacore T200 at 25 °C. Dual flow cells were used with a blank reference. For immobilisation, Biacore CM5 sensor chips were activated by flowing N-hydroxysuccinimide (NHS) and N-(3-dimethyl-aminopropyl)-N’-ethyl-carbodiimide hydrochloride (EDC) for 420 s with a flow rate of 10 μl/min in PBS-P+. CypA (provided by Morten Govasli) at 50 μg/ml in 10 mM sodium acetate pH 5 was flowed over the surface for 420 s before quenching for 420 s with ethanolamine. To improve binding site accessibility, CypA was preincubated with 60 μM bulky CsA analogue JW-47 for 10 min prior to injection. Transduction enhancers were diluted in HBS-EP+ buffer with 2% DMSO to 20 μM-4.88 μm and flowed over the surface for 250 s followed by 1200 s dissociation. After each injection, a wash of 50% DMSO 50% HBS-EP+ was performed. Solvent correction was carried out with six DMSO concentrations between 1% and 3%. All compound injections were run in duplicate.

### Fluorescence polarisation (FP)

The binding of TEs to CypA was measured by FP assay as previously described (*42*). Briefly, assays were conducted in 384-black low flange non-binding microtiter plates (Corning, Inc.). A total solution of 80 μl was used consisting of three components: fluorescent cyclosporine probe (FP-CsA) (45 nm), CypA (40 nm), and inhibitor (10–10000 nm). CypA and inhibitor were incubated in the OmegaFluostar at room temperature for 30 min, FP-CsA (40 μl) was added last. Measurements were taken after gain adjustment of control sample and 30 min incubation with a xenon flashlight with filter settings for 485 nm excitation and 520 nm emission. At least three replicates were used for each experiment.

### Cell lines

U87 and HEK 293T were maintained in Dulbecco’s Modified Eagle Medium (DMEM) (Gibco, 41966-029), 10% FCS, 1% penicillin/streptomycin (Gibco, 15140-122) 37 °C in 5% CO_2_ (U87) and 10% CO_2_ (293T). THP-1 cells were maintained at between 2×10^5^ and 1×10^6^ cells/ml in Roswell Park Memorial Institute medium (RPMI) (Gibco, 21875-034), 10% FCS, 37 °C, 5% CO2.

### Protein overexpression in THP-1

To generate cells overexpressing IFITM1/2/3, 4×10^5^ THP-1 cells in a volume of 0.75 ml were mixed with 1 ml viral vector with pSIN, or media for mock and 8 μg/ml polybrene and centrifuged at 1000 xg for 1h at room temperature (spinoculation). Supernatants were removed and cells resuspended in 2 ml media, transferred to 6-well plates and incubated until they reached a density of 1×10^6^ cells/ml. Cells were cultured with 1 μg/ml puromycin until all mock cells had died, monitored by trypan blue stain. Cells were periodically treated with 1 μg/ml puromycin to maintain overexpression.

### Production of lentiviral vector in HEK293T cells

A confluent plate of HEK 293T cells was split 1:4 into 10 cm dishes. 24 h later, 1 μg p8.91 + 1 μg pMDG + 1.5 μg pCSGW (LV-GFP) was mixed with 200 μl Opti-MEM (Gibco, 31985070) and 10 μl FuGENE 6 transfection reagent (Promega, E2692) and added dropwise to the cells. 24 h post-transfection, the media was replaced. 48- and 72-h post-transfection supernatants containing lentiviral vector were harvested, filtered (0.45 μm), combined and used at a multiplicity of infection (MOI) of 0.25 on THP-1 or U87 cells.

### Screening CsA analogues for transduction enhancement of lentiviral infection THP-1 and U87 cells

U87 cells were seeded 1.25×10^4^ cells/well in 250 μl in 48-well plates and allowed to settle overnight. Where necessary, THP-1 cells at 8×10^5^ cells/ml were incubated for 24 h with 10 ng/ml IFNβ (Peprotech, 300-02BC) to induce IFITM3 expression. For infection, THP-1 cells were typically seeded in 48-well plates at a density of 5.6×10^4^ cells/well. Inhibitors and LV-GFP (MOI 0.25) were added to a final volume of 200 μl for THP-1 or 250 μl for U87s. For mock infections, virus was replaced with media, each condition was performed in triplicate. After 48 h incubation, cells were fixed with 4% PFA and the proportion of GFP positive cells measured by flow cytometry using a Novocyte (Agilent).

### Testing of cellular process inhibitors

THP-1 cells were activated by IFNβ or DMSO control and treated with 5 µM CsH/BG147 or 1% LB and cellular process inhibitors: MG132 proteasome inhibitor 20 µM, MLN4924 NEDDylation inhibitor 1 µM (951950-33-7, Sigma Aldrich), BafA1 lysosome inhibitor 0.5 µM, EGA late endosome trafficking inhibitor 20 µM.

### Flow cytometry

For LV-GFP lentiviral infection of THP-1 U87, cells were fixed in 4% formaldehyde in PBS. The % GFP+ (infected) cells were determined using a NovoSampler Pro (Agilent) flow cytometer.

### Immunoblot

For immunoblot, 1×10^6^ cells were washed with PBS and lysed in 50 μl RIPA buffer (150 mM sodium chloride, 1.0% Triton X-100, 0.5% sodium deoxycholate, 0.1% SDS and 50 mM Tris pH 8.0 with phosphatase and protease inhibitors (cOmplete mini, EDTA-free, Roche) by incubation for 10 min on ice. Samples were centrifuged for 10 min at 13200 rpm at 4 °C. Total protein was quantified and adjusted using a Pierce™ BCA Protein Assay Kit (Thermofisher, 23225). Supernatants were mixed with 4X Laemmli sample buffer (200 mM Tris pH 6.8, 8% SDS, 0.4% bromophenol blue, 40% glycerol and 5% betamercaptoethanol). Samples were incubated at 95 °C for 10 min. Proteins were separated by SDS-PAGE on Mini-PROTEAN TGX 4-20% gels (Bio-Rad, 4561093) and transferred onto nitrocellulose membrane using the Trans-Blot Turbo Transfer System (Bio-Rad, 1704270). Membranes were blocked for 1 h in 5% non-fat dry milk PBS-Tween (PBS, 0.1% Tween-20). Primary antibodies were incubated for 1 h at rt or overnight at 4°C. Membranes were washed three times in PBS-Tween, incubated with IRDye® fluorescent secondary antibodies (LI-COR) for 1 h at rt. After washing with PBS-Tween and PBS, blots were visualised at 700 nm and 800 nm on a LICOR Odyssey CLx. Protein markers are indicated in kDa throughout. Densitometry was performed using Empiria Studio and protein densities were calculated by normalisation to β-actin loading control. Data are presented as the mean of two independent experiments.

### Antibodies

Immunoblot primary antibodies: Anti-IFITM1 (60074-1-Ig, Proteintech, 1:20000), Anti-IFITM2 (66137-1-Ig, Proteintech, 1:1000), Anti-IFITM3 (109429, Abcam, 1:1000), Anti-GAPDH (G9545, Sigma-Aldrich, 1:5000), Anti-HA (H6908, Sigma-Aldrich, 1:1000), Anti-Tetherin (BST2, sc-390719, Santa Cruz Biotechnology), Anti-MxB (HPA030235, Sigma-Aldrich, 1/1000). Anti-β-actin (clone AC-15, ab6276, Abcam, 1:10000). Immunoblot secondary antibodies: IRDye® 680LT goat anti-mouse (926-68020, LI-COR Biosciences, 1:15000). IRDye® 680LT goat anti-rabbit (926-68021, LI-COR Biosciences, 1:15000). IRDye® 800CW goat anti-mouse (926-32210, LI-COR Biosciences, 1:10000). IRDye® 800CW goat anti-rabbit (926-32211, LI-COR Biosciences, 1:10000). Immunofluorescence microscopy primary antibodies: Primaries, Anti-IFITM3 (109429, Abcam, 1:100), anti-EEA1 (610456, BD biosciences, 1:250), anti-LAMP1 (ab25630, Abcam, 1:250), anti-Giantin (ab37266, Abcam, 1:500). Immunofluorescence microscopy secondary antibodies: Goat anti-mouse 546 (A11003, Invitrogen, 1:450), and goat anti-rabbit 488 (A11034, Invitrogen, 1:450)

### Viability assay

3-(4,5-dimethylthiazol-2-yl)-2,5-diphenyltetrazolium bromide (MTT) reagent (10% total volume, 5 mg/ml in PBS) was added to TE-treated THP-1 or U87 cells in a 96-well plate, after 1-2 h at 37 °C cells crystal metabolites were solubilised by addition of 100 µL of 10% SDS, 0.01 M HCL and incubated for 4 at 37 °C. Spectrometer (Odyssey) used to measure absorbance at 570 nm.

### CD34+ Cell Culture

CD34+ Cells were cultured in “StemSpan Classic II”: StemSpan™ SFEM II (STEMCELL Technologies, #09655), supplemented with FLT-3 (#300-19), SCF (#300-07) and TPO (**#**300-18) at 100 ng/mL, and IL-3 (#200-03) at 60 ng/mL (all supplements from Peprotech), with 1% Pen/Strep (Gibco, 15140-122). Cells were cultured at 1×10^6^ cells/mL, and kept at 37 °C, 5% CO_2_.

### Transduction Enhancer Evaluation in CD34+

24 h post-thaw, cells were counted and resuspended in fresh StemSpan Classic II at 3.125×10^6^ cells/ml. 80 µL (250k) cell suspension was used per condition in a flat-bottomed 96-well plate. LentiBOOST ® (LB) was added at 1% and Protamine Sulphate (PS) at 4 ug/mL final concentration. Transduction enhancers and Lentivirus were added to final concentrations or MOIs stated in figures, to a final volume of 125 µL/well. Cells were incubated for an additional 18 h.

After incubation, cells were transferred to a V-bottomed 96-well plate, centrifuged for 10 mins at 300 x g. Supernatant was carefully aspirated, cells were resuspended in 1 mL fresh StemSpan Classic II and transferred to a 24-well plate. Cells were then incubated for an additional 2 or 9 days before GFP expression was measured. At the chosen timepoint, cells were washed once with 1 mL MACS buffer (PBS + 0.5% BSA, 0.45 µM-filtered) before staining with 1 µL of antibodies: anti-CD34 (BV421, Bio legend 343610), anti-CD38 (APC/Cy7, Biolegend 303534), anti-CD45RA (APC, Biolegend 304112), anti-CD90 (PE/Cy7, Biolegend 328124), in 50 µL of MACS buffer. Cells were pelleted again and resuspended in 300 µL MACS buffer, with DAPI added at 1:200 immediately before Flow Cytometry on a BD® LSR II Flow Cytometer. Results were analysed using FlowJo™ v10.10 Software (BD Life Sciences).

### Quantitative PCR (qPCR)

TRIM5 and IFITM3 expression in measured by qPCR TaqMan assay, normalised to the housekeeping gene OAZ1 and transformed to 2-ΔCt. THP-1 cells (with and without IFNβ), U87 cells and two HSC donors, were resuspended in 300 μl RLT buffer (74106, Qiagen,) with 1% betamercaptoethanol. RNA was extracted using RNeasy purification kit (74106, Qiagen), including on-column DNAse digest for 15 min at 25°C using an RNase-Free DNase Set (79254, Qiagen). Followed by TURBO DNase^TM^ (AM2238, Thermo) treatment, as per manufactures instructions. RNA was quantified using a nanodrop.

For cDNA synthesis reactions, 70 ng RNA, 1 μl dNTP and 1 μl OligoDT were made up to 13 μl in H_2_O. After step 1, reactions were incubated at 65°C for 5 min and then transferred to ice for 1 min. Reverse transcription was done with Superscript IV Reverse transcriptase (18090050, Invitrogen), reactions were incubated at 50 °C for 60 min then 70 °C for 15 min. cDNA was diluted 1/3. qPCR reactions were set up in 384-well plates, each condition was run in duplicate, negative controls without RNA template or reverse transcriptase were included. TaqMan qPCR set up using 2x TaqMan gene expression master mix (4369016, Applied Biosystems) and FAM/TAMRA probes for OAZ1 (Hs00427923_m1), TRIM5 (Hs01552559_m1), and IFITM3(Hs03057129_s1).

The plate was sealed with MicroAmp Optical Adhesive Film (4311971, Applied Biosystems), centrifuged at 1200 rpm for 2 min and qPCR performed using a QuantStudio5 (Thermofisher Scientific) using cycling parameters 50 °C 10 min extension, 85 °C 5 min, 4 °C cool down 30 cycles. Analysis was carried out using Thermofisher Design and Analysis software.

### U87 TRIM5 KO by CRISPR/Cas9

Lentiviral particles to generate CRISPR/Cas9-edited cell lines were produced by transfecting 10 cm dishes of HEK293T cells with 1.5 μg of TRIM5 CRISPR vector (kind gift of Adam Fletcher(*31*)), 1 μg of p8.91 packaging plasmid (*61*) and 1 μg of VSV-G glycoprotein-expressing plasmid pMDG (Genscript) using Fugene-6 transfection reagent (Promega). Virus supernatants were collected at 48 and 72 h post transfection and pooled. U87 cells were seeded at 1×10^5^ cells/well in a 6-well plate and 1 ml/well of vector added. Transduced cells were selected by culturing with 1 μg/ml puromycin (Merck Millipore) for 48 h.

### MLV

U87 treated with non-targeting guides (WT control) and U87 TRIM-5 KO were seeded at 5×10^4^ cells/well in a 48-well plate and left overnight. N-tropic murine leukaemia virus (MLV-N) and TRIM5-insensitive MLV-Bzotero, prepared as previously described (*46*), were titrated onto the cells in 250 μL. After 48 h incubation, cells were fixed with 4% PFA and the proportion of GFP positive cells measured by flow cytometry using a Novocyte (Agilent).

### Molecular modelling

Molecular docking of JW3-158 in CypA was performed with the Molecular Operating Environment (MOE) software, Chemical Computing Group.

### Immunofluorescent microscopy

THP-1 were treated with (or without) 10 ng/ml IFNϕ3 overnight (7×10^5^ cells/ml). THP-1 then treated with DMSO, CsH or BG147 (2.5 µM) and with and without BafA1 (0.5 µM) for 6 h (all) or 24 h (BafA1 untreated). Cells were taken in parallel for infection assay with HIV-LV-GFP and immunoblot as described above. Glass coverslips were prepared with poly-L-lysine (0.01% 37°C for 1 h), washed with ddH_2_0 and dried. Samples were transferred to coverslips (200 µl) and left to settle for 2 h before fixing with 6% PFA (400 µl, final 4% PFA). After 24 h slides were washed 3 x PBS. Slides treated with 150 mM ammonium chloride in PBS for 5 min to quench PF autofluorescence with followed by washing 3 x in PBS. Cells permeabilised with 0.1% Triton X-100 in PBS for 5 min followed by washing 3 x in PBS. Blocking step with 5% FCS in PBS (300 µl) for 30 min at room temp followed by addition of 50 µl 1° Ab mix for 45 min (Rb anti-IFITM3 1/100 and either Ms anti-EEA1 1/250, Ms anti-LAMP1 1/250 or Ms anti-Giantin 1/500). Slides were washed 3x with 5% FCS in PBS followed by incubation of 50 µl 2° Abs mix for 60 min in dark (goat anti-mouse 546 1/450, goat anti-rabbit 488 (green) 1/450). Cells were washed 2x with 5% FCS in PBS, 1x PBS, 2x H_2_O. Coverslips were mounted on slides with a 30 µl drop of Mowiol + DAPI, allowed to dry and then stored at 4 °C.

Images were recorded using the LSM_800 Zeiss LSM800 upright point scanning confocal, 60x with oil. Images presented are medial slices from a captured z-stack of 4-6 slices. For quantification z stacks were pooled over multiple cells in 6-8 captured fields, Pearson correlation coefficient (PCC), correlation was quantified by calculating the linear relationship between IFITM3 protein and the different cellular markers in multiple cells.

### Subretinal injections in mice

Subretinal injections were performed in mice to deliver viral vectors under direct visualization. Lentivirus was a pLenti CMV GFP vector (Addgene 17446), with titre >5×10^6^. Anaesthesia was induced using a mixture of ketamine (6 mg/ml) and medetomidine hydrochloride (Dormitor, 0.1 mg/ml) diluted in sterile water. Pupillary dilation was achieved with topical application of Tropicamide 0.5% and Phenylephrine 2.5%. Following gentle prolapse of the eye, the fundus was visualised using a contact lens system composed of Viscotears as a coupling medium and a glass coverslip. A 34-gauge, 10 mm needle mounted on a 5 µl Hamilton syringe was introduced tangentially through the superior equatorial sclera and advanced into the subretinal space under direct observation. Vector suspension was injected to produce a localized bullous retinal detachment, confirming successful subretinal delivery. The sclerotomy self-sealed, allowing vector retention within the subretinal space until absorption by the retinal pigment epithelium, typically within 48 h. In some cases, an additional injection was performed in the inferior hemisphere to create a secondary detachment. Patent retinal vasculature was confirmed in all injected eyes post-injection. Anaesthesia was reversed with intraperitoneal administration of 0.15 ml atipamezole hydrochloride (Antisedan, 0.10 mg/ml), and mice were monitored during recovery under mild warming.

### Immunohistochemistry and microscopy of mouse and human retina

Mouse eyes were rapidly enucleated and fixed in 4% paraformaldehyde at 4°C. Whole retinas were dissected from the eye and incubated with primary antibodies in 3% Triton X-100 at 4°C overnight. Retinas were washed three times in PBS and incubated with secondary antibodies and conjugated PNA in 3% Triton X-100 at 4°C overnight. Retinas were washed three times in PBS and counterstained with Hoechst 33342. Retinas were cleared using clearing solution for 2 h at RT (*62*). Retinas were flat-mounted in the same clearing solution between two coverslips and attached to an adhesion slide for imaging. Injected eyes were fixed in 4% PFA for 2 days, dissected, cleared for 2 h in the tissue clearing mentioned above at RT and flat-mounted for imaging.

Human eyes (Moorfields Lions Eye Bank) were fixed in 10% neutral buffered formalin at 4°C. 5 mm x 5 mm sections of human retinal tissue were dissected from the eye and prepared following the same process as the mouse retinas.

Primary antibodies used: IFITM1-Specific Monoclonal antibody 60074-1-IG (1:200), IFITM2-Specific Monoclonal antibody 66137-1-IG (1:200), Anti-Fragilis antibody EPR5242 (1:200). Secondary antibodies used: Donkey anti-mouse IgG (H+L), Alexa Fluor™ 647 A-31571 (1:200), Donkey anti-Rabbit IgG (H+L) Alexa Fluor™ 594 A-21207 (1:200), Donkey anti-Rabbit IgG (H+L) Alexa Fluor™ 647 A-31573 (1:200), Lectin PNA From Arachis hypogaea (peanut), Alexa Fluor™ 488 Conjugate L21409 (1:200), Lectin PNA From Arachis hypogaea (peanut), Alexa Fluor™ 594 Conjugate L32459 (1:200). Images were acquired using a confocal microscope (Leica Stellaris 8 with LAS X software), 40x with oil. Images presented (Fig. 6E, 6F) are single slices from captured z-stacks of the whole retina. For quantification z-stacks through the entire retina were used for z-profile of fluorescence intensity. Images (Fig. 6G) were acquired from merged tile scans using LAS X-integrated Navigator tool. Fluorescence quantification was performed using *Fiji* (ImageJ) with built-in analysis plugins. Fluorescence intensity was measured by drawing regions of interest (ROIs) around individual cell bodies. Signal intensity was normalized to background fluorescence within each image. Background measurements were obtained from at least five randomly selected regions at different locations and depths within each image stack. For each retina, a random sample of up to 35 transduced cells was analysed. If fewer than 35 transduced cells were present in the stack, all available cells were included. The percentage of transduced cells was quantified in *Fiji* by an experimenter blinded to the experimental condition (Lenti alone vs. Lenti + BG147). Transduced and non-transduced cells were distinguished based on fluorescence intensity: non-transduced cells exhibited only low levels of autofluorescence, which, when enhanced, revealed a negative contrast of the nucleus. Quantification was performed on three randomly selected sections per retina, each taken from a different depth of the outer segments (OS).

### Statistical analysis

Statistical analyses were performed using GraphPad Prism (v10.3.1) as indicated in the figure captions.

## Supporting information

Supplementary figures and Chemistry Materials and Methods

## Acknowledgments

The authors would like to acknowledge Anthony Nolan (Charity no. 803716/SC038827) for the provision of Peripheral Blood Stem Cell (PBSC) donations.

## Funding

Wellcome Collaborative award (214344, G.J.T., D.L.S.),

Wellcome Investigator Award 220863 G.J.T

MRC award (MR/S023380/1 G.J.T.)

Wellcome trust principal investigator award (217112/Z/19/Z, A.T.)

The Rosetrees Trust (ID2020/100020, LGT., G.J.T., D.L.S.)

NIHR UCLH/UCL Biomedical research centre (BRC) (BRC1191/III/101350, D.A.)

UCL TechFund (UTF2-21-007 Towers & Selwood / UCL, G.J.T., D.L.S., G.S.)

## Author contributions

Conceptualization: DA GS LGT DLS GJT AT KLM

Methodology: VP DLS LGT DA GJT KLM GS MR KP

Investigation: DA KLM BJC VP BG JW LGT NK MW ST TN AH DL TEW

Visualization: DA KLM BG LGT

Funding acquisition: GJT DLS GS LGT DA AT

Project administration: GJT

Supervision: GJT DLS GS

Writing – original draft: DA LGT VP

Writing – review & editing: DA LGT GJT KLM DLS BJC HA LSN NK

## Data and materials availability

All data are available in the main text or the supplementary materials.

