## Supplementary figures and Chemistry Materials and Methods for "A modified cyclosporine enhances lentivector transduction *ex vivo* and *in vivo* by degrading IFITM3"

**Fig. S1 to 8**

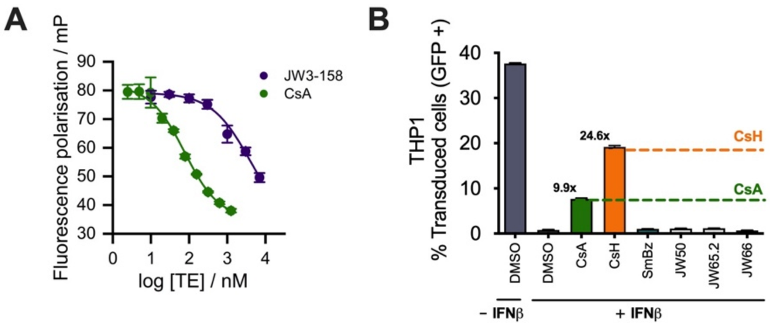


**Fig. S1: Screen of CsA-SmBz-like compounds (A)** Fluorescence polarization of JW3-158 and CsA binding to CypA. **(B)** LV-GFP transduction of THP-1 pretreated with +/- 10 ng/ml IFNβ, with 5 μM CsA-SmBz analogues added at the time of infection. % GFP positive cells measured at 48 hpi, n=3 +/-SEM.

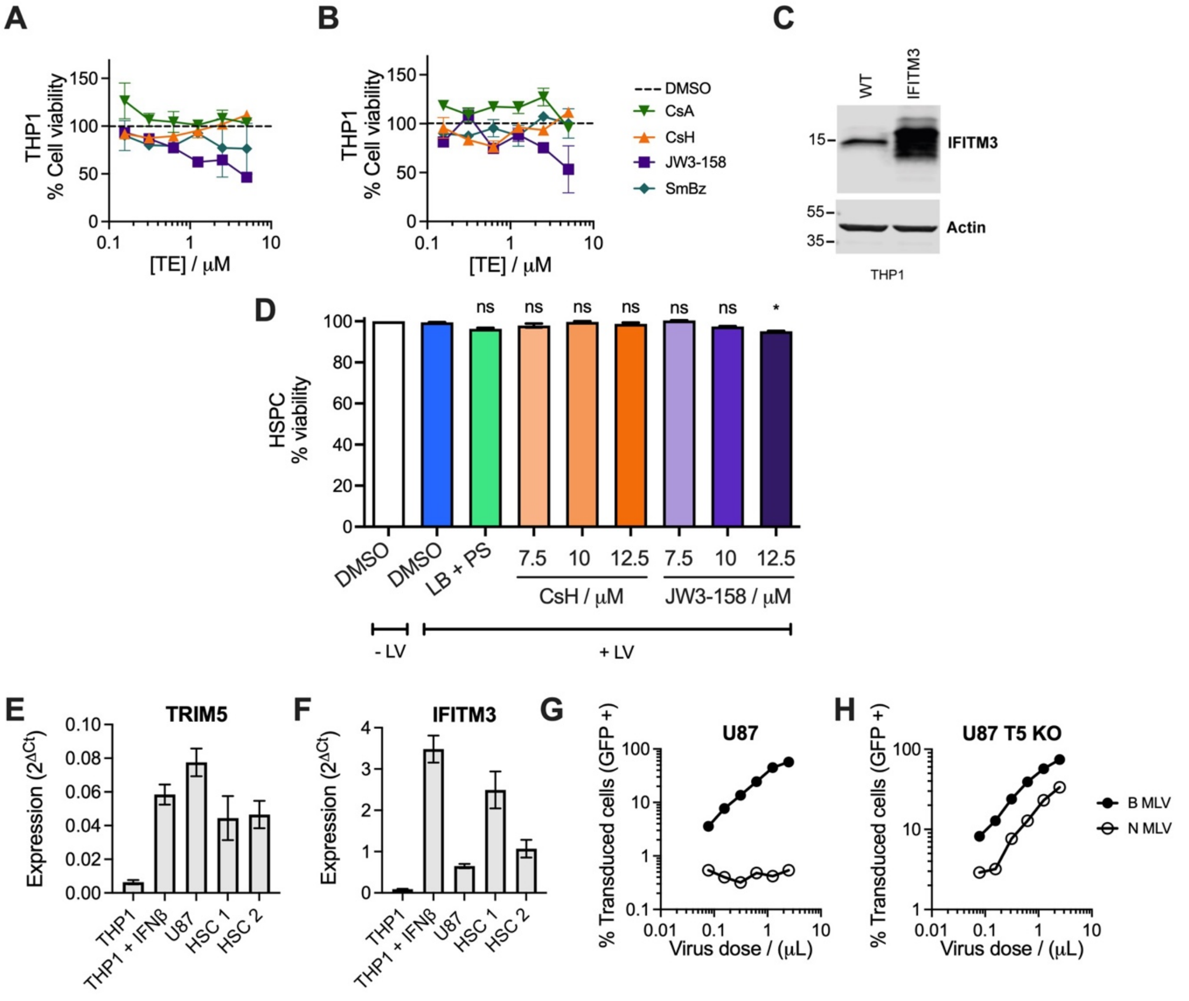


**Fig. S2. Cyclosporin-like transduction enhancers degrade IFITM3 and inhibit Cyclophilin A (A-B)** MTT cell viability assay of THP-1 cells without (A) and with (B) 10 ng/ml IFNβ pre-treatment in the presence of escalating doses of TEs, n= 3, +/- SEM. **(C)** Overexpression of IFITM3 in THP-1 cells. **(D)** HSPC treated with 1% LB and 4 μg/ml PS, or novel TE as shown, and infected with LV-GFP at an MOI of 10 IU/cell, percentage cell viability by trypan blue staining, n=2, +/- SEM. One way ANOVA with Dunnett's test compared to DMSO+LV, **; $p < 0.0332$, *; $p < 0.1234$. **(E-F)** TRIM5 and IFITM3 expression in THP-1 cells (+/- 20 ng/ml IFNβ pre-treatment), U87 cells and two HSPC donors measured by RT-qPCR, normalised to the housekeeping gene OAZ1 and transformed to 2-ΔCt. **(G-H)** WT and TRIM5 KO U87 cells infected with B- or N-tropic MLV-GFP over a range of viral doses. % GFP-positive cells measured at 48 hpi, n=3, +/- SEM.

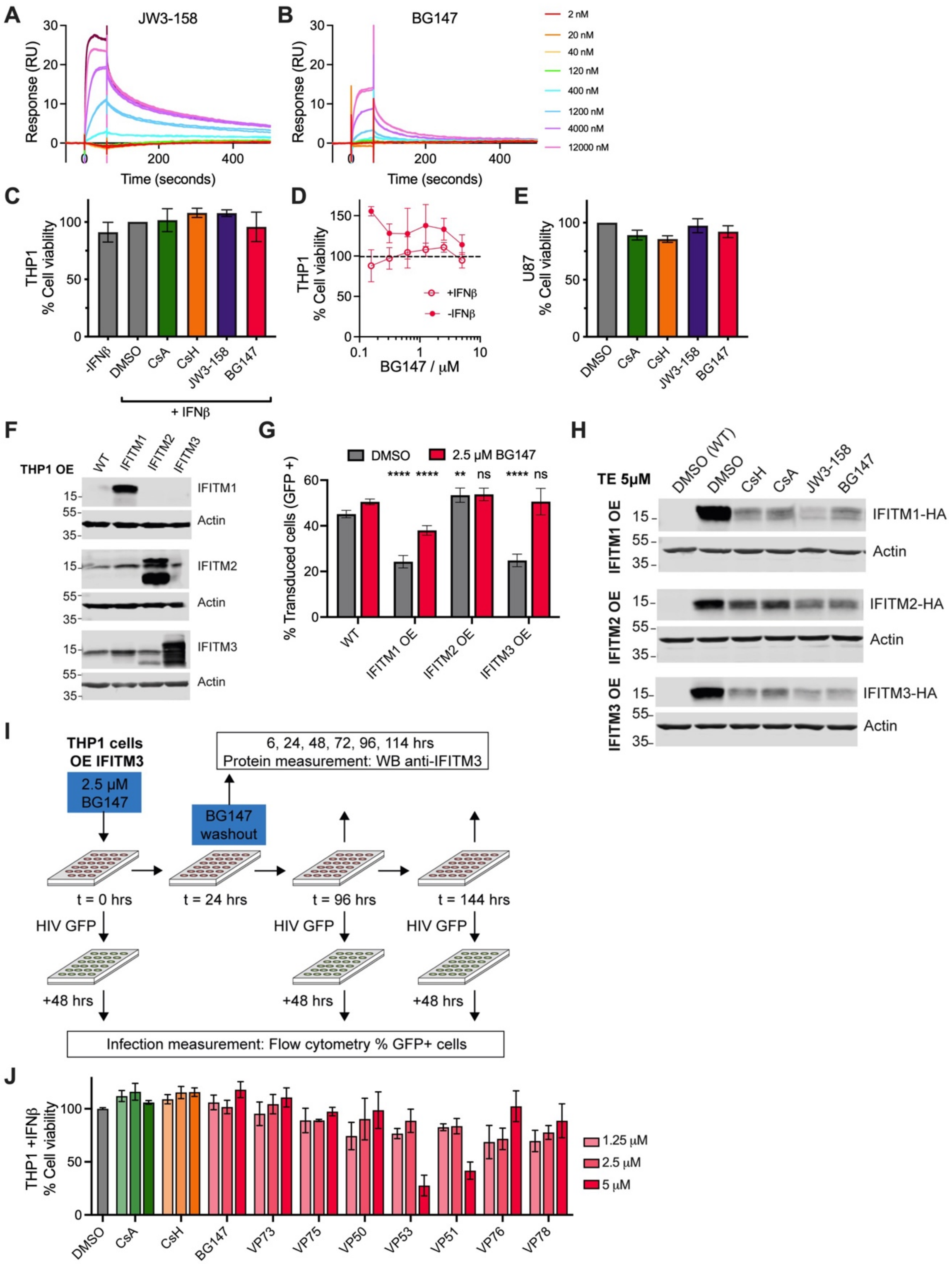


**Fig. S3: Structure-based design of a BG147. (A-B)** JW3-158 (A) or BG147 (B) (flow) and CypA (immobilised on chip) in representative SPR experiments. **(C-E)** MTT cell viability assay of THP-1 cells with and without IFNβ (C-D) and U87 cells (E) in the presence of 5 μM TEs (C, E) or different concentrations of BG147 (D), n=3, +/- SEM. **(F)** Overexpression of IFITM1-HA, IFITM2-HA or IFITM3-HA in THP-1 cells, n=1. **(G)** WT THP-1 cells and THP-1 cells overexpressing IFITM1-HA, IFITM2-HA or IFITM3-HA were treated with 2.5 μM BG147 or DMSO and transduced with LV-GFP. % GFP positive cells measured at 48 hpi, n=3, +/- SEM. Two-way ANOVA with Tukey's test compared to corresponding DMSO or BG147 treated WT cells. $p \leq 0.0001$, ****; $p \leq 0.0002$, ***; $p \leq 0.0021$, **; $p < 0.0332$, *; $p < 0.1234$ ns. **(H)** Levels of IFITM1-HA, IFITM2-HA and IFITM3-HA in THP-1s treated with 5 µM TEs or DMSO for 24 h. n=1. **(I)** Schematic of BG147 washout experiment. THP-1 cells overexpressing IFITM3-HA were treated with 5 µM BG147 or DMSO. After 24 h media was changed, and cells were washed 3 times with PBS. Samples were taken to measure IFITM3-HA levels at 6, 24, 48, 72, 96 and 144 h. **(J)** MTT cell viability assay of Fig. 3M, THP-1 cells from, pretreated with +/- 10 ng/ml IFNβ treated with 1.25, 2.5 and 5 µM TE or BG147 derivatives, n=2, +/-SEM.

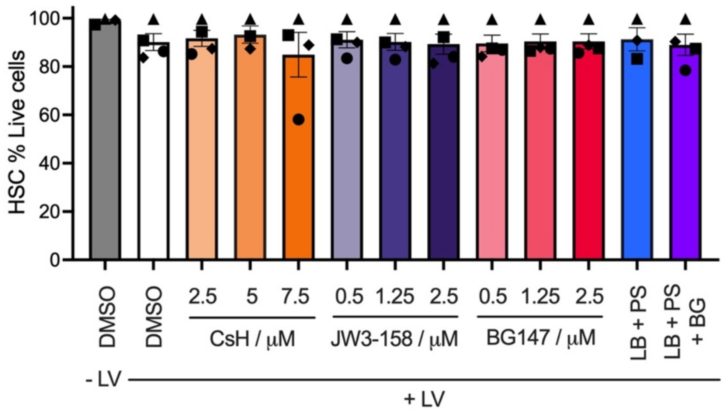


**Fig. S4: BG147 is novel, efficient transduction enhancer in HSPCs.** HSPC treated with 1% LB and 4ug/ml PS, or novel TE as shown, and infected with LV-GFP (LV) at an MOI of 10 GC/cell, % Live cells measured at 48 hpi by trypan blue staining, n=4 HSPC donors, represented by different symbols, +/- SEM.

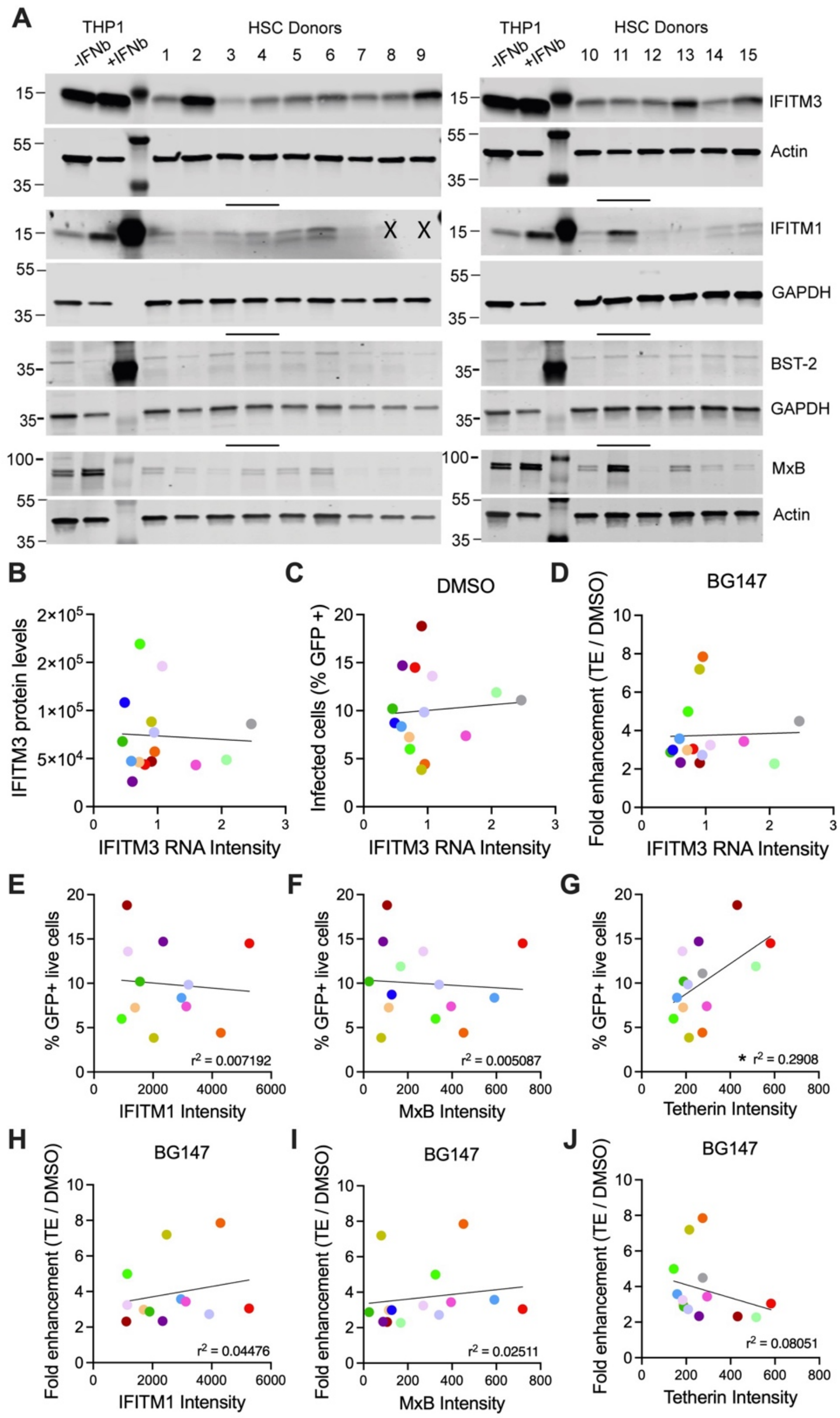


**Fig. S5: Interferon-stimulated genes in HSCs. (A)** Levels of IFITM3, IFITM1, tetherin and MxB protein, lanes that did not transfer well (indicated by an X) or outliers with very high IFITM1, tetherin or MxB levels were excluded from further analysis. **(B-D)** Correlation of 2-ΔCq IFITM3 expression from qPCR TaqMan assay, normalised to *oaz*1 with (B) IFITM3 protein levels measured by WB (A) for each donor, normalized against actin, (C) base level transduction and (D) fold enhancement with 2.5 µM BG (Fig. 4B-C). Simple linear regression, Pearson correlations insignificant. **(E-J)** Correlation of IFITM1, MxB and tetherin protein levels measured by WB (C), normalized against actin or GAPDH with base level transduction (E-G) and fold enhancement (H-J) with 2.5 µM BG (Fig.4 B-C). Simple linear regression, only (G) Pearson significant.

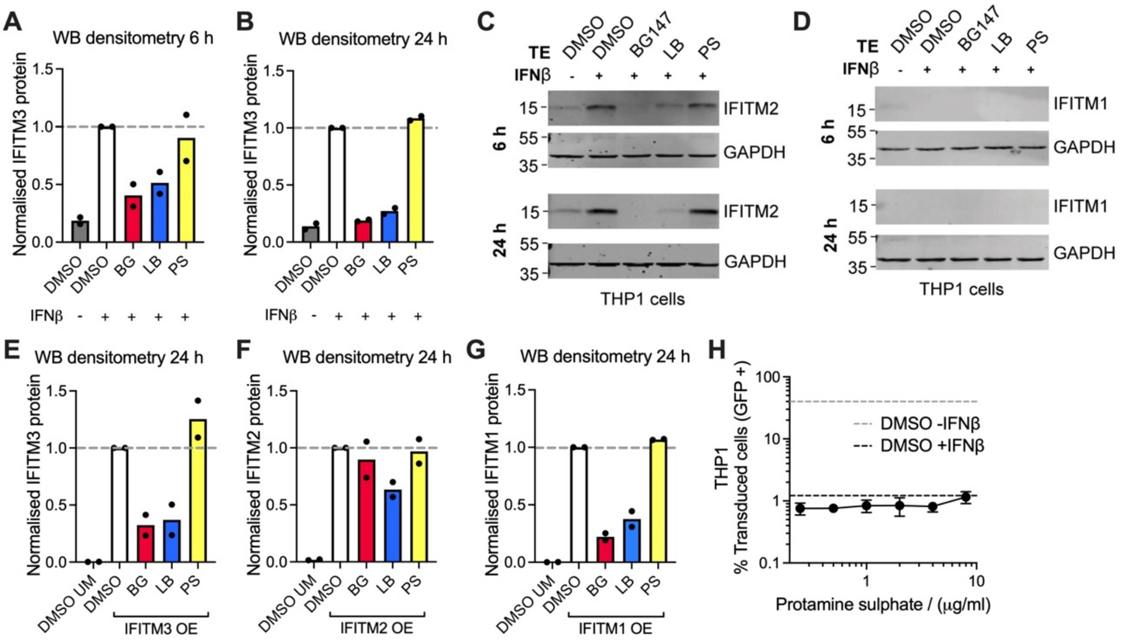


**Fig. S6: Transduction enhancer Lentiboost causes IFITM3 degradation. (A)** IFITM3 protein quantification from Fig. 5a normalised to actin and expressed as a percentage of DMSO+IFNβ, n=2. **(C-D)** IFITM2 (C) and IFITM1 (D) levels at 6 and 24 h post treatment with 1% LB and 4 μg/ml PS, 2.5 μM BG147 (BG), or a combination, in unmodified THP-1 pretreated with +/- 20 ng/ml IFNβ. n=2. **(E-G)** IFITM3 protein quantification from Fig. 5B normalised to actin and expressed as a percentage of DMSO+IFNβ, n=2. **(H)** Titration of PS on THP-1 pretreated with +/- 10 ng/ml IFNβ, or in THP-1 over-expressing IFITM3, infected with LV-GFP for 48 h, +/-SEM 3 replicates, n = 2.

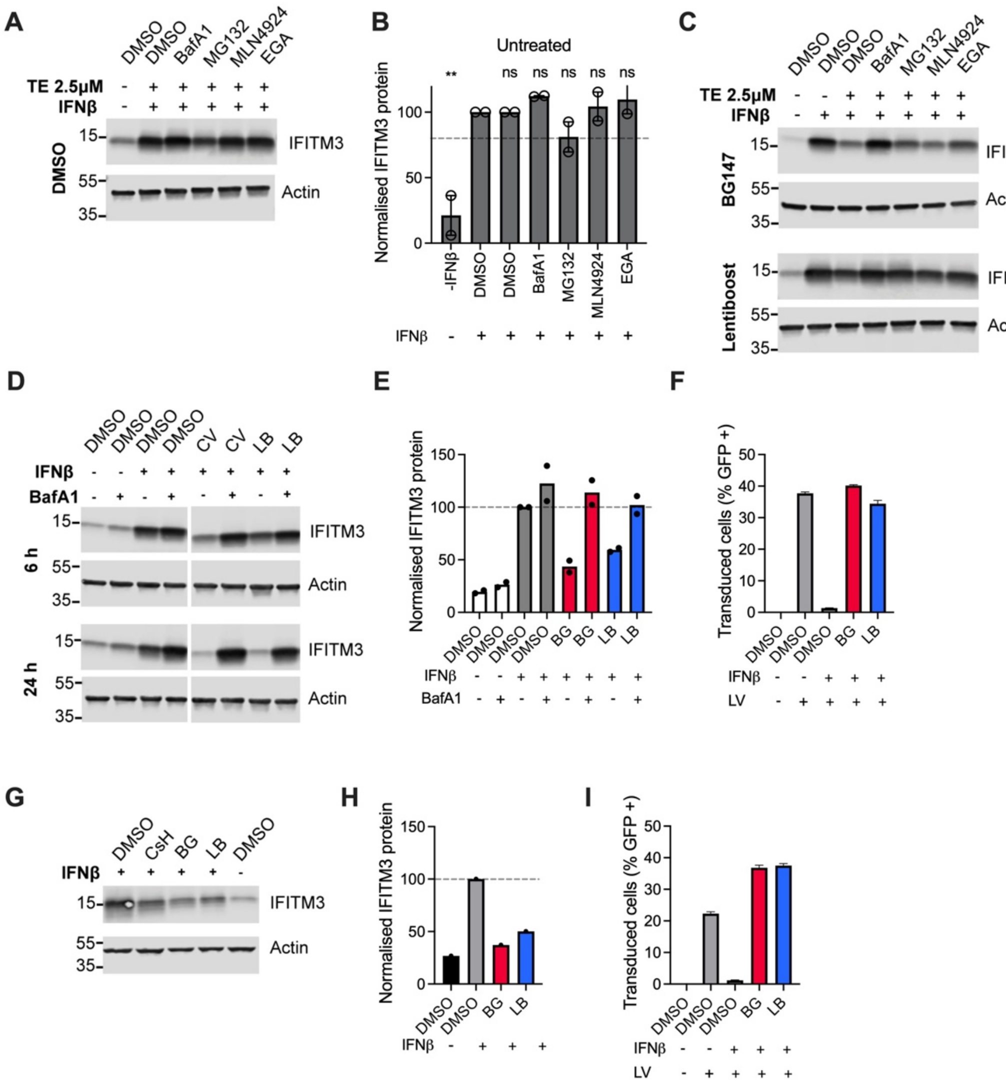


**Fig. S7: TE-dependent depletion of IFITM3 protein is caused by aberrant trafficking**. **(A-C)** IFITM3 levels at 6 h post treatment with DMSO (A-B), 1% LB or 2.5 µM BG147 (BG) (C), and cellular process inhibitors (MG132 proteasome inhibitor 20 µM, MLN4924 NEDDylation inhibitor 1 µM, BafA1 lysosome inhibitor 0.5 µM, EGA late endosome trafficking inhibitor 20 µM) in THP-1 pretreated with +/- 20 ng/ml IFNβ. n=3. **(D-F)** THP-1s from microscopy assay Fig. 5H were infected with LV-GFP (% GFP positive cells measured at 48 hpi) and IFITM3 levels measured after 6 or 24 h. IFITM3 protein density normalised to actin and plotted relative to THP-1 + IFNβ + DMSO, n=2. **(G-I)** THP-1s from microscopy assay Fig. 5J were infected with LV-GFP (% GFP positive cells measured at 48 hpi) and IFITM3 levels measured after 6 or 24 h. IFITM3 protein density normalised to actin and plotted relative to THP-1 + IFNβ + DMSO, n=2.

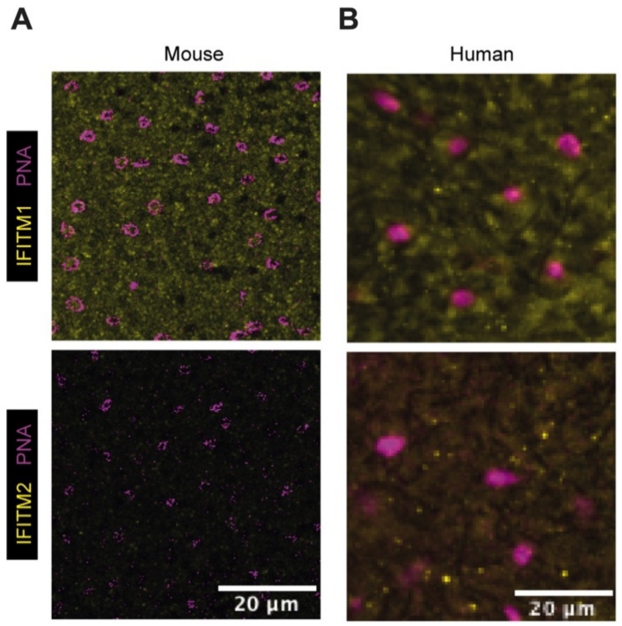


**Fig. S8: IFITM proteins in mouse and human retina. (A-B)** Confocal microscopy and quantification of IFITM1 or IFITM2 levels (yellow) in the outer nuclear layer of mouse (A) and human (B) retina. Photoreceptor cone cells are detected by Peanut agglutinin lectin (PNA) marker (pink).

### Chemistry Materials and Methods

- All **commercially available** solvents and reagents were used without further treatment as received unless otherwise noted.
- Solvent ***evaporation*** were performed under reduced pressure (40-60 °C) by using Buchi Rotavapor R-210.
- ***Microwave-assisted reactions*** were carried out in Biotage Initiator$^{+}$ apparatus for synthesis.
- Reactions under ***anhydrous atmosphere*** were performed using commercial $N_2$.
- **Thin layer chromatography (TLC)** was performed with qualitative purposes on aluminium silica gel plates (Alugram Sil G/UV 254) with detection by UV light (λ 254 nm) and by staining with [$(NH_4)_6MoO_4$, $Ce(SO_4)_2$, $H_2SO_4$, $H_2O$].
- **Purifications** were carried in a Biotage Isolera Four Flash Chromatography System. The chromatography column used were: a) normal phase Biotage Sfär Silica (5g, 10g, 25g), and b) reverse phase Biotage Snap Bio C4 300 Å 25g or Biotage Sfar C18 D-Duo 100 Å (12g, 30g, 60g). PE-AX 10g, strong base ion exchange resin was used when indicated.
- **$^{1}$H- and $^{13}$C-NMR spectra** were performed at UCL Chemistry NMR Facility and Bruker DRX 500, 600 or 700 MHz spectrometer were used. Chemical shifts (δ) are expressed in ppm relative to TMS as an internal standard and coupling constants (*J*) in Hz. *J* are assigned and not repeated. $CDCl_3$ and $CD_3OD$ were used as solvents at room temperature except when indicated. All the assignments were confirmed by 2D spectra (COSY and HSCQ).
- **Accurate mass measurements** using an ASAP-HESI ionisation connected to the Q Exactive Plus mass spectrometer were performed at UCL Chemistry Mass Spectrometry Facility.
- **LC-MS spectra** were obtained using a single quadrupole LC/MSD XT mass spectrometer with electrospray ionisation (ESI), using an analytical C4 column (Symmetry 300 , 50 x 4.6 mm, 3.5 μm) and C18 column (Kinetex 5 μm 100 Å, 50 x 4.6 mm). FA: Formic acid.The gradients used were:
  a) 10% → 95% of MeCN + 0.1% FA[1] in $H_2O$ + 0.1% FA (6.5 min)
  b) 30% → 95% of MeCN + 0.1% FA in $H_2O$ + 0.1% FA (9 min)
  c) 30% → 95% of MeCN in $H_2O$ 10 mM $NH_3$ (9 min)

Software used:

- **LC-MS spectra analysis:** Agilent MassHunter Software , MestraNova Software Version 12.0.3 - 22023
- **LC-MS Chromatography Method Development:** Agilent ChemStation Software
- **NMR spectra analysis:** MestreNova Software Version 12.0.3 - 22023

### Synthesis and characterization of novel compounds.

#### 1. Synthesis of the first generation of transduction enhancers (JW series).

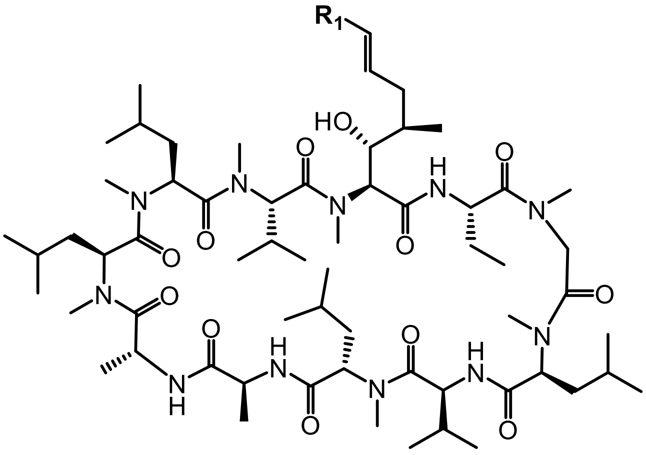


**CsA-R1**

All CsA modifications at the R1 position were obtained following the general method A described below.

**General Method A**

To a solution of Cyclosporin A (1 eq) in DCM (2 mL) was added the corresponding alkene ( 1.2 eq) and Hoveyda-Grubbs 2nd generation catalyst (17 mol%). The reaction was stirred in the microwave at 90 °C for 30 minutes and then allowed to cool to r.t. Triethylamine was added to the mixture and then stirred overnight with excess $P(CH_2OH)_3$ to coordinate the ruthenium catalyst. This was then washed away with brine and water before the mixture was passed through a Stratospheres PL Thiol MP SPE cartridge (polymer Lab, Varian Inc) to remove any remaining catalyst.

**JW47**, prepared as previously described *(42)***.**

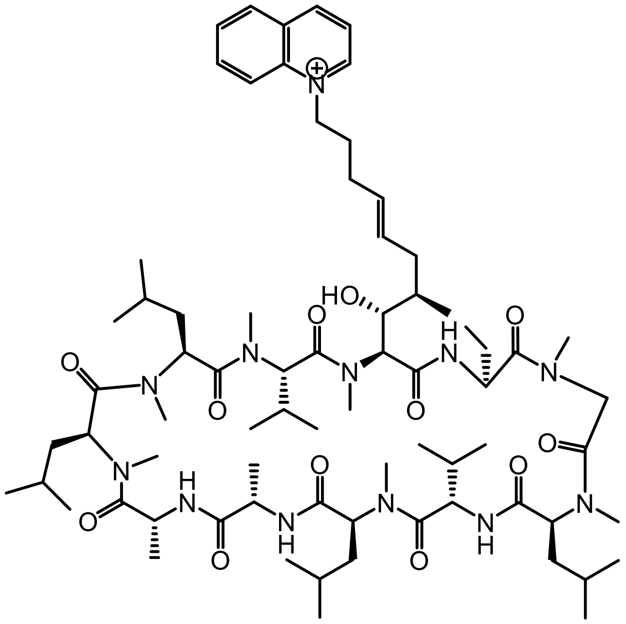

**JW60-2** prepared as in *(63)*.

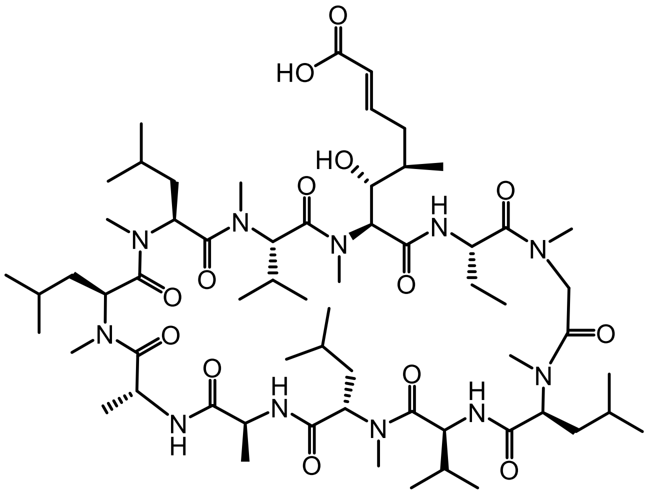

The crude product obtained following general procedure A and was subsequently purified by flash silica chromatography (5% MeOH in DCM). Repurified by reverse-phase column (10 – 100% MeCN-0.1% FA in water-0.1% FA) to give JW60-2 as a brown solid.

**JW115**

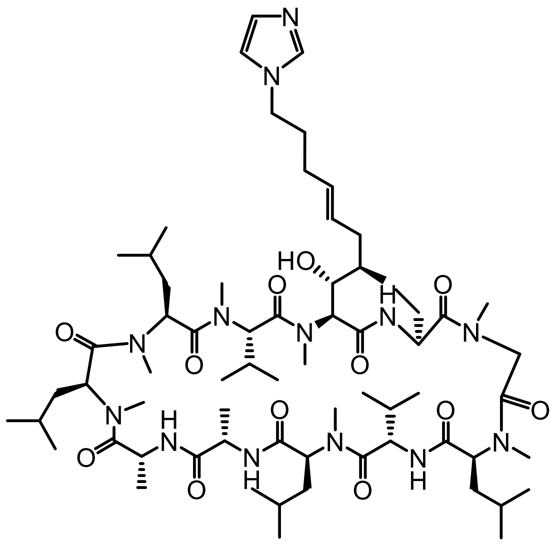

The crude product was purified by flash silica chromatography (5% MeOH in DCM) to give JW115 as a white solid (11.4 mg, 11%).

$^1$H NMR (700 MHz, $CDCl_3$) δ 3.49 (s, NMe, 3H), 3.39 (s, NMe, 3H), 3.23 (s, NMe, 3H), 3.11 (s, NMe, 3H), 3.69 (s, NMe, 3H), 2.69 (s, NMe, 3H), 2.67 (s, NMe, 3H).

**JW2-65**

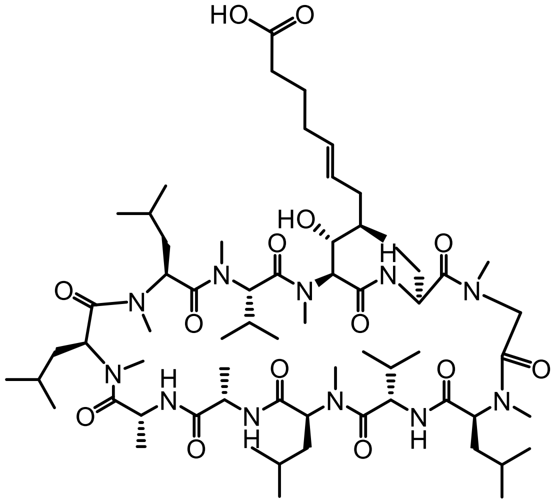

The crude product was purified by flash silica chromatography (5% MeOH in DCM). Repurified by reverse-phase column (10 – 100% MeCN-0.1% FA in water-0.1% FA) to give JW2-65 as a white powder (18 mg, 9%).

LCMS (*m/z*): $[MH]^+$ calcd. for $C_{65}H_{115}N_{11}O_{14}$, 1274.70; found 1274.80.

$^1$H NMR (600 MHz, $CDCl_3$) δ 3.47 (s, NMe, 3H), 3.38 (s, NMe, 3H), 3.23 (s, NMe, 3H), 3.10 (s, NMe, 3H), 3.09 (s, NMe, 3H), 2.70 (s, NMe, 3H), 2.68 (s, NMe, 3H).

**JW3-157**

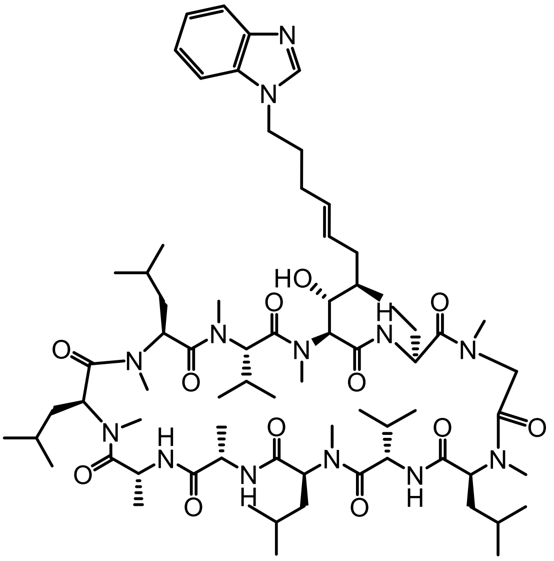

The crude product was purified by flash silica chromatography (5% MeOH in DCM) to give JW3-157 as an off-white powder (18 mg, 16% yield).

MS (m/z): [MH]+ calcd. for $C_{71}H_{119}N_{13}O_{12}$, 1346.81; found 1346.92.

$^{1}$H NMR (600 MHz, $CDCl_3$) δ 3.51 (s, NMe, 3H), 3.39 (s, NMe, 3H), 3.22 (s, NMe, 3H), 3.12 (s, NMe, 3H), 3.11 (s, NMe, 3H), 2.70 (s, NMe, 3H), 2.68 (s, NMe, 3H).

**JW3-158**

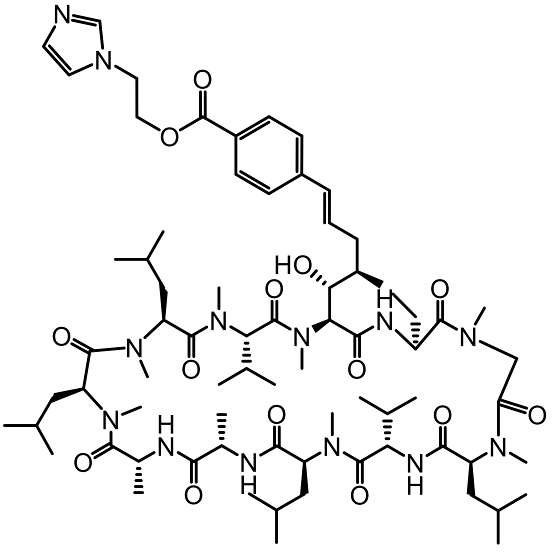

The crude product was purified by flash silica chromatography (5% MeOH in DCM). Repurified by reverse-phase column (10 – 100% MeCN-0.1% FA in water-0.1% FA) to give JW3-158 as an off-white powder (13 mg, 38% yield).

HRMS (*m/z*): $[MH]^+$ calcd. for $C_{73}H_{119}N_{13}O_{14}$, 1402.9072; found 1402.9072.

$^{1}$H NMR δ 3.49 (s, NMe, 3H), 3.35 (s, NMe, 3H), 3.23 (s, NMe, 3H), 3.12 (s, NMe, 3H), 3.07 (s, NMe, 3H), 2.72 (s, NMe, 3H), 2.69 (s, NMe, 3H).

**JW3-159**

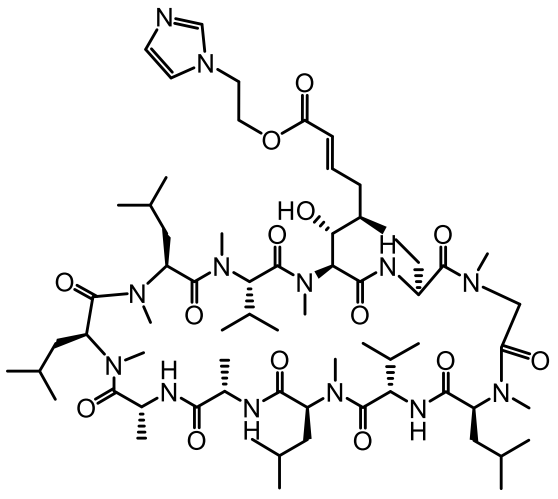

The crude product was purified by flash silica chromatography (5% MeOH in DCM). Repurified by reverse-phase column (10 – 100% MeCN-0.1% FA in water-0.1% FA) to give JW3-159 as an off-white powder (23 mg, 36% yield).

LCMS (*m/z*): $[MH]^+$ calcd. for $C_{67}H_{115}N_{13}O_{14}$, 1326.73; found 1326.88.

$^1$H NMR (600 MHz, $CDCl_3$) δ 3.52 (s, NMe, 3H), 3.32 (s, NMe, 3H), 3.16 (s, NMe, 3H), 3.15 (s, NMe, 3H), 3.07 (s, NMe, 3H), 2.96 (s, NMe, 3H), 2.71 (s, NMe, 3H).

**JW3-173**

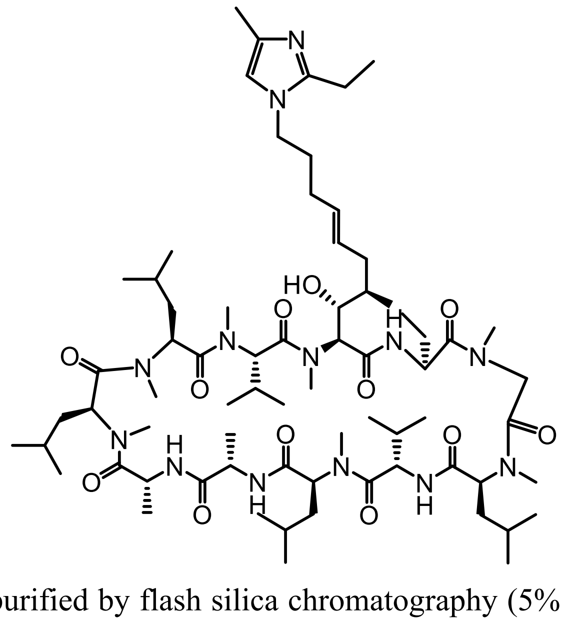

The crude product was purified by flash silica chromatography (5% MeOH in DCM) to give JW3-173 as an off-white powder (49 mg, 11% yield).

MS (m/z): [MH]+ calcd. for $C_{70}H_{123}N_{13}O_{12}$, 1338.83; found 1338.95.

$^1$H NMR (600 MHz, CDCl3) δ 3.46 (s, NMe, 3H), 3.36 (s, NMe, 3H), 3.20 (s, NMe, 3H), 3.09 (s, NMe, 3H), 3.08 (s, NMe, 3H), 2.67 (s, NMe, 3H), 2.65 (s, NMe, 3H).

**JW3-177**

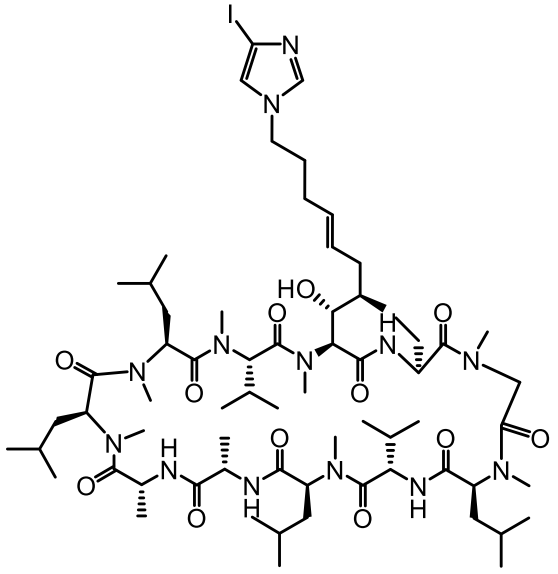

The crude product was purified by flash silica chromatography (5% MeOH in DCM) to give JW3-177 as an off-white powder (92 mg, 39% yield).

LCMS (m/z): [MH]+ calcd. for $C_{67}H_{116}IN_{13}O_{12}$, 1422.65; found 1422.80.

$^{1}$H NMR (600 MHz, $CDCl_3$) δ 3.49 (s, NMe, 3H), 3.38 (s, NMe, 3H), 3.22 (s, NMe, 3H), 3.12 (s, NMe, 3H), 3.11 (s, NMe, 3H), 2.69 (s, NMe, 3H), 2.67 (s, NMe, 3H).

**JW3-175** and **JW3-180**

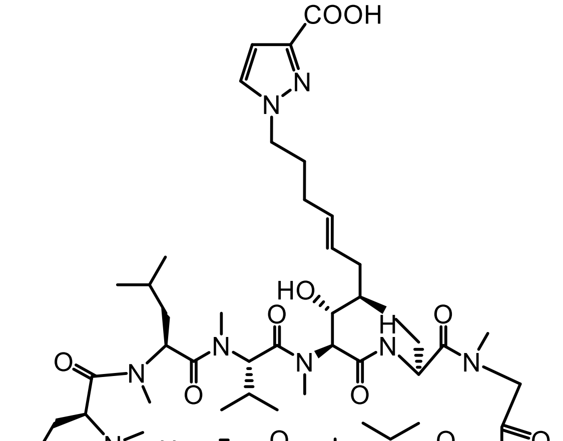

**JW3-180**

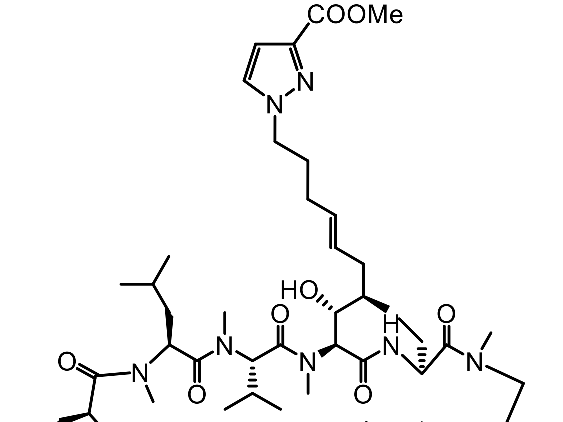

**JW3-175**

The crude product of JW3-175 was purified by flash silica chromatography (5% MeOH in DCM) to give crude JW3-175 as an off-white powder (50 mg, 0.037 mmol). The crude was then dissolved in THF before the addition of lithium hydroxide monohydrate (3 mg, 0.074

mmol) and water. This mixture was stirred at room temperature overnight. Purification was carried out by PE-AX column primed with 10cvs of 10% $NH_4OH$ in MeOH solution, then 5cvs of MeOH and 10cvs of 5% MeOH in DCM solution. Solution of crude product in MeOH was loaded and the column eluted with 5cvs of 5% MeOH in DCM solution. Column then eluted with 0-100% of 5% AcOH/ 5% MeOH in DCM solution in 5% MeOH in DCM solution.

JW3-180 was isolated as an off-white solid, 21mg (5% yield).

Data for JW3-180

LCMS (m/z): [MH]+ calcd. for $C_{68}H_{117}N_{13}O_{14}$, 1340.76; found 1340.90.

$^1$H NMR (600 MHz, $CDCl_3$) δ 3.49 (s, NMe, 3H), 3.38 (s, NMe, 3H), 3.23 (s, NMe, 3H), 3.11 (s, NMe, 3H), 3.10 (s, NMe, 3H), 2.69 (s, NMe, 3H), 2.68 (s, NMe, 3H).

**JW3-183B**

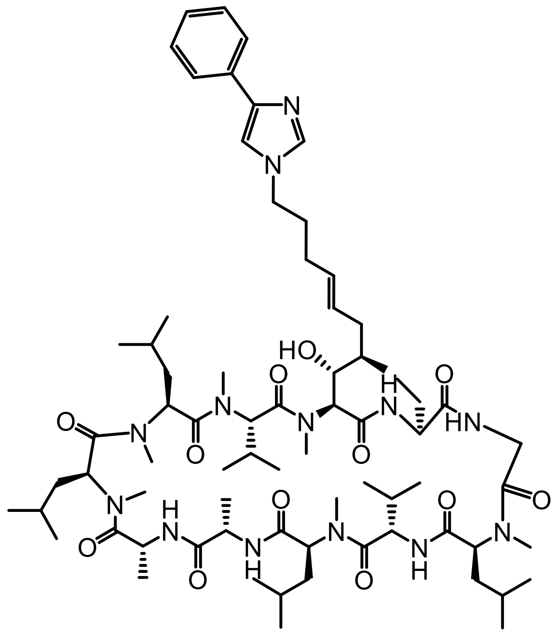

The crude product was purified by flash silica chromatography (5% MeOH in DCM) to give JW3-183B as an off-white powder (17 mg, 8% yield).

LCMS (m/z): [MH]+ calcd. for $C_{72}H_{119}N_{13}O_{12}$, 1358.82; found 1358.92.

$^1$H NMR (600 MHz, $CDCl_3$) δ 3.48 (s, NMe, 3H), 3.39 (s, NMe, 3H), 3.25 (s, NMe, 3H), 3.12 (s, NMe, 3H), 3.11 (s, NMe, 3H), 2.68 (s, NMe, 3H), 2.67 (s, NMe, 3H).

**JW3-190**

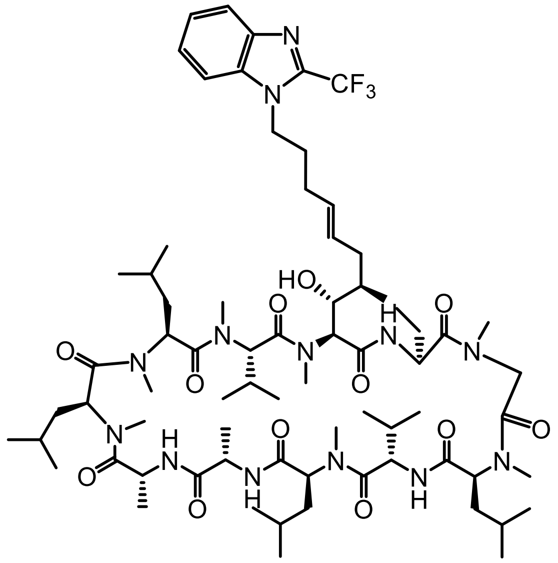

The crude product was purified by flash silica chromatography (5% MeOH in DCM) to give JW3-190 as an off-white powder (15 mg, 6% yield).

LCMS (m/z): [MH]+ calcd. for $C_{72}H_{118}F_3N_{13}O_{12}$, 1414.81; found 1414.90.

$^1$H NMR (600 MHz, $CDCl_3$) δ 3.50 (s, NMe, 3H), 3.39 (s, NMe, 3H), 3.22 (s, NMe, 3H), 3.11 (s, NMe, 3H), 3.10 (s, NMe, 3H), 2.69 (s, NMe, 3H), 2.67 (s, NMe, 3H).

**JW3-191**

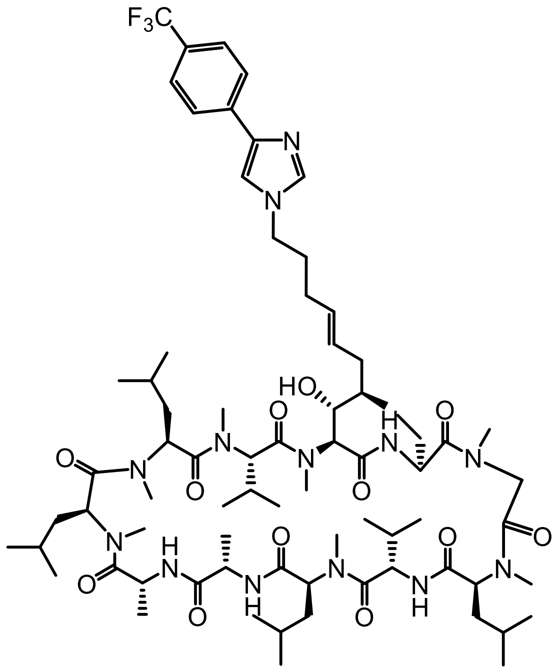

The crude product was purified by flash silica chromatography (5% MeOH in DCM) to give JW3-191 as an off-white powder (18 mg, 29% yield).

LCMS (m/z): [MH]+ calcd. for $C_{74}H_{120}F_3N_{13}O_{12}$, 1440.85; found 1140.92.

$^1$H NMR (600 MHz, $CDCl_3$) δ 3.50 (s, NMe, 3H), 3.39 (s, NMe, 3H), 3.23 (s, NMe, 3H), 3.11 (s, NMe, 3H), 3.10 (s, NMe, 3H), 2.69 (s, NMe, 3H), 2.67 (s, NMe, 3H).

**JW4-2**

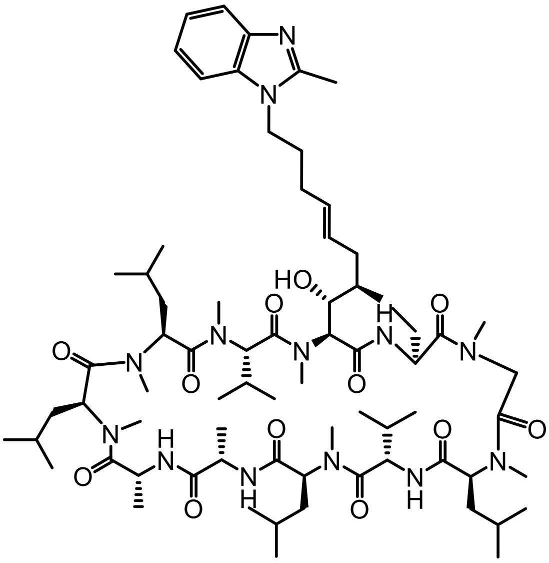

The crude product was purified by flash silica chromatography (5% MeOH in DCM) to give JW4-2 as an off-white powder (34 mg, 15%).

LCMS (m/z): [MH]+ calcd. for $C_{72}H_{121}N_{13}O^{12}$, 1360.84; found 1360.93.

$^{1}H$ NMR (600 MHz, $CDCl_3$) δ 3.50 (s, NMe, 3H), 3.39 (s, NMe, 3H), 3.23 (s, NMe, 3H), 3.11 (s, NMe, 3H), 3.10 (s, NMe, 3H), 2.69 (s, NMe, 3H), 2.68 (s, NMe, 3H).

**JW4-3**

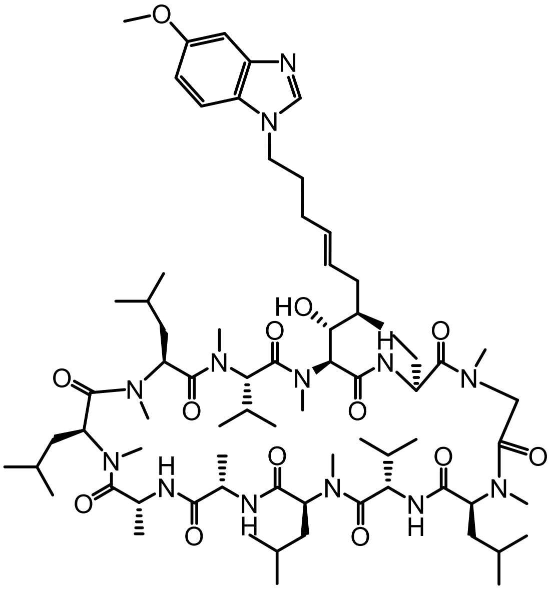

The crude product was purified by flash silica chromatography (5% MeOH in DCM) to give JW4-3 as an off-white powder (47 mg, 21% yield).

LCMS (m/z): [MH]+ calcd. for $C_{72}H_{121}N_{13}O_{13}$, 1376.84; found 1376.93.

$^{1}H$ NMR (600 MHz, $CDCl_3$) δ 3.49 (s, NMe, 3H), 3.38 (s, NMe, 3H), 3.20 (s, NMe, 3H), 3.10 (s, NMe, 3H), 3.09 (s, NMe, 3H), 2.68 (s, NMe, 3H), 2.66 (s, NMe, 3H).

**JW4-4**

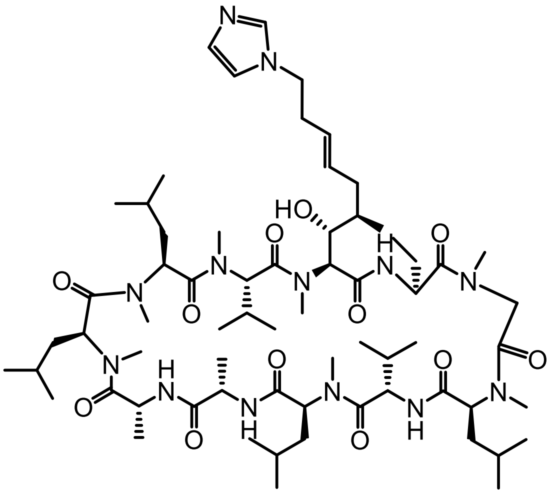

The crude product was purified by flash silica chromatography (5% MeOH in DCM) to give JW4-4 as an off-white solid (22 mg, 10% yield).

MS (m/z): [MH]+ calcd. for $C_{66}H_{115}N_{13}O_{12}$, 1282.73; found 1282.89.

$^{1}$H NMR (600 MHz, $CDCl_3$) δ 3.48 (s, NMe, 3H), 3.39 (s, NMe, 3H), 3.25 (s, NMe, 3H), 3.13 (s, NMe, 3H), 3.12 (s, NMe, 3H), 2.69 (s, NMe, 3H), 2.67 (s, NMe, 3H).

**JW4-35**

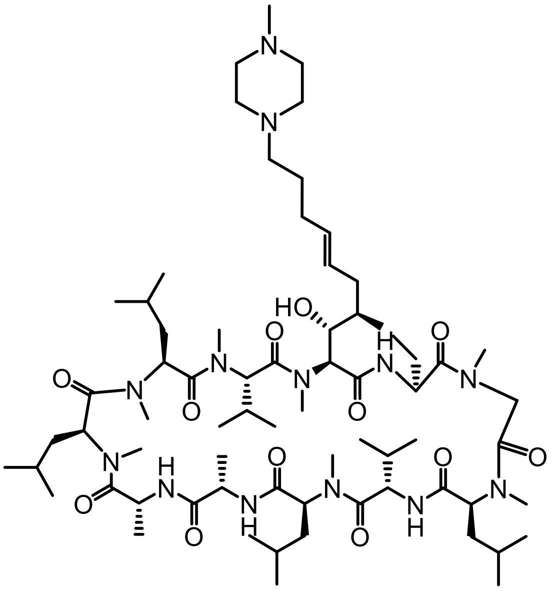

The crude product was purified by flash silica chromatography (5% MeOH in DCM) to give JW4-35 as an off-white powder (10 mg, 9% yield).

MS (*m/z*): $[MH]^+$ calcd. for $C_{69}H_{125}N_{13}O_{12}$, 1328.84; found 1328.96.

$^{1}$H NMR (600 MHz, $CDCl_3$) δ 3.49 (s, NMe, 3H), 3.39 (s, NMe, 3H), 3.24 (s, NMe, 3H), 3.12 (s, NMe, 3H), 3.10 (s, NMe, 3H), 2.70 (s, NMe, 3H), 2.68 (s, NMe, 3H).

**JW4-12**

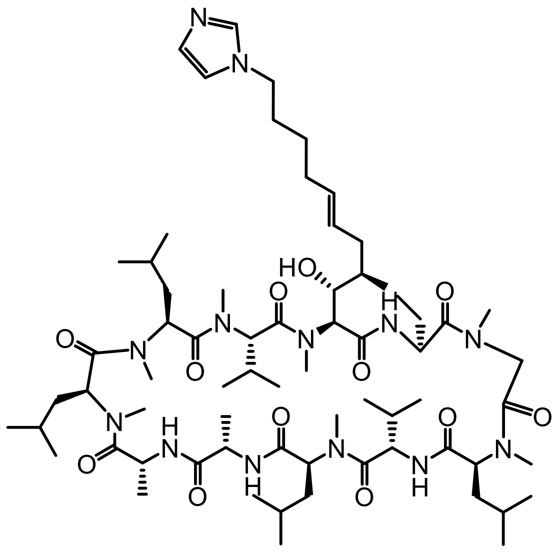

The crude product was purified by flash silica chromatography (5% MeOH in DCM) to give JW4-12 as an off-white powder (24 mg, 11% yield).

LCMS (m/z): [MH]+ calcd. for $C_{68}H_{119}N_{13}O_{12}$, 1310.78; found 1310.92.

$^{1}$H NMR (600 MHz, CDCl3) δ 3.49 (s, NMe, 3H), 3.38 (s, NMe, 3H), 3.23 (s, NMe, 3H), 3.11 (s, NMe, 3H), 3.10 (s, NMe, 3H), 2.69 (s, NMe, 3H), 2.68 (s, NMe, 3H).

**JW4-13**

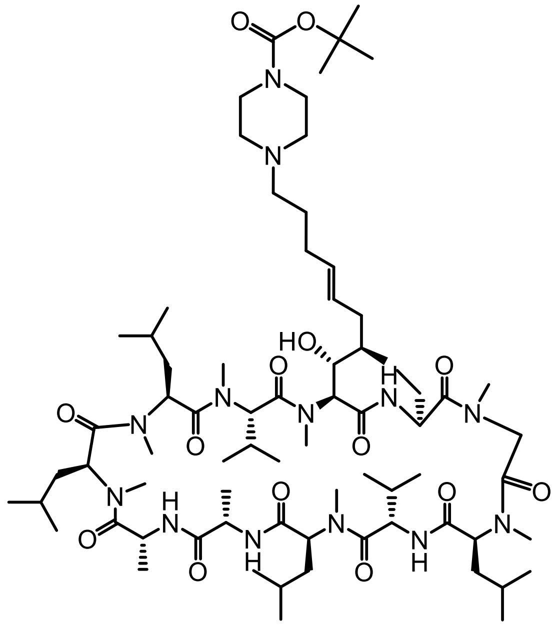

The crude product was purified by flash silica chromatography (5% MeOH in DCM) to give JW4-13 as a pale brown solid (108 mg, 30%).

LCMS (m/z): [MH]+ calcd. for $C_{73}H_{131}N_{13}O_{14}$, 1414.93; found 1415.00.

$^{1}$H NMR (600 MHz, $CDCl_3$) δ 3.48 (s, NMe, 3H), 3.37 (s, NMe, 3H), 3.23 (s, NMe, 3H), 3.11 (s, NMe, 3H), 3.08 (s, NMe, 3H), 2.69 (s, NMe, 3H), 2.67 (s, NMe, 3H)

**JW4-16**

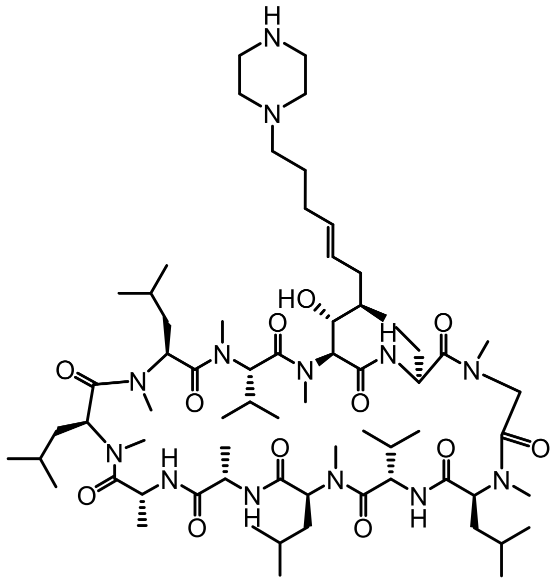

The crude product was purified by flash silica chromatography (5% MeOH in DCM) to give JW4-16 as an off-white powder (14 mg, 11% yield).

LCMS (m/z): [MH]+ calcd. for $C_{68}H_{123}N_{13}O_{12}$, 1314.81; found 1314.95.

$^1$H NMR (600 MHz, $CDCl_3$) δ 3.49 (s, NMe, 3H), 3.39 (s, NMe, 3H), 3.25 (s, NMe, 3H), 3.12 (s, NMe, 3H), 3.10 (s, NMe, 3H), 2.70 (s, NMe, 3H), 2.69 (s, NMe, 3H).

**JW4-17**

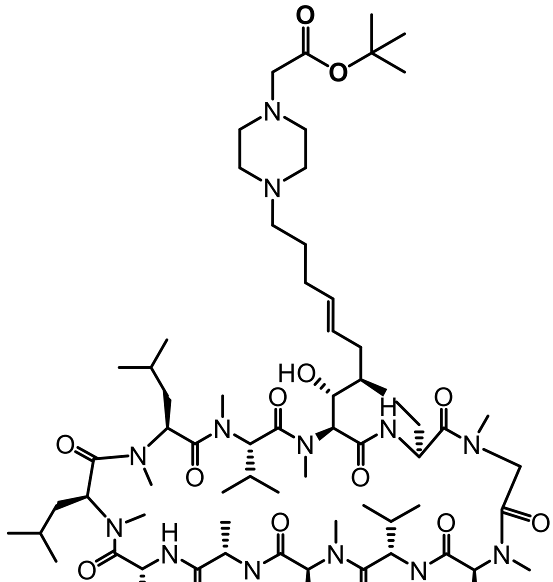

The crude product was purified by flash silica chromatography (5% MeOH in DCM) to give JW4-17 as a brown solid (25 mg, 21% yield).

MS (*m/z*): [MH]+ calcd. for $C_{74}H_{133}N_{13}O_{14}$, 1428.96; found 1429.02.

$^{1}$H NMR (600 MHz, $CDCl_3$) δ 3.48 (s, NMe, 3H), 3.38 (s, NMe, 3H), 3.23 (s, NMe, 3H), 3.11 (s, NMe, 3H), 3.09 (s, NMe, 3H), 2.70 (s, NMe, 3H), 2.68 (s, NMe, 3H).

**TWH246**

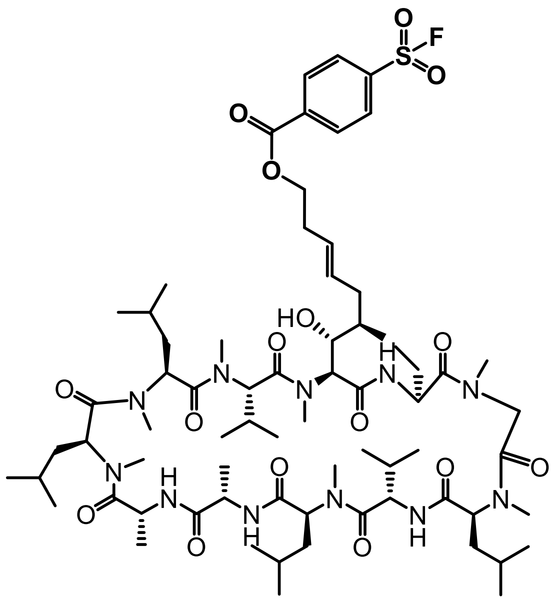

The crude product was purified by flash silica chromatography (5% MeOH in DCM) to give TWH246.

LCMS (*m/z*): [MH]+ calcd. for $C_{70}H_{116}FN_{11}O_{15}S$, 1401.84; found 1402.84.

**JW3-163A**

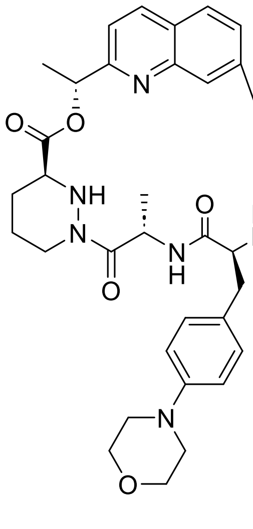


(R)-1-(7-bromoquinolin-2-yl)ethyl(S)-1-(((S)-2-(but-3-enamido)-3-(4-morpholinophenyl)propanoyl)-L-alanyl)hexahydropyridazine-3-carboxylate was disolved in dry MeCN (20 mL) followed by Palladium(II) acetate (0.1 eq), Tri(o-tolyl)phosphine (0.2 eq), triethylamine (2 eq). The reaction mixture was stirred in a MW at 150 °C for 2.5h. The solution was then filtered through celite and purified by chromatography column on solica gel (MeOH 0% to 10% in DCM). The final product was isolated as an off-white solid (24%).

LCMS (*m/z*): $[MH]^+$ calcd. for $C_{36}H_{43}N_6O_6$, 655.77; found 655.40.

**JW3-185**

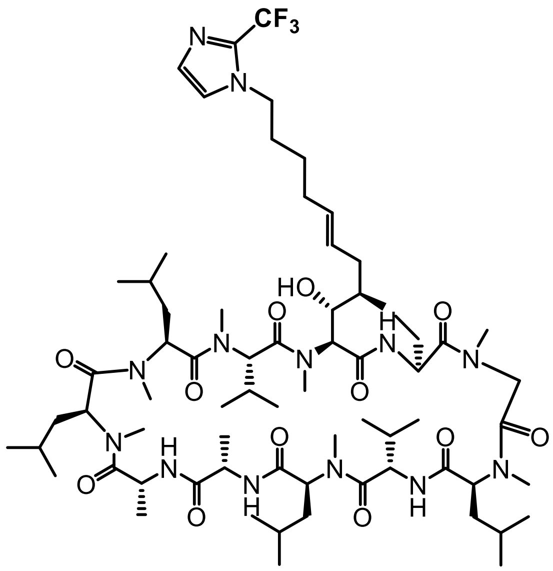


The crude product was purified by flash silica chromatography (5% MeOH in DCM) to give JW3-185 as an off-white powder (40 mg, 18% yield).

HRMS (*m/z*): $[MH]^+$ calcd. for $C_{68}H_{116}F_3N_{13}O_{12}$, 1364.8891; found 1364.8924.

$^1H$ NMR (600 MHz, $CDCl_3$) δ 3.50 (s, NMe, 3H), 3.40 (s, NMe, 3H), 3.24 (s, NMe, 3H), 3.12 (s, NMe, 3H), 3.11 (s, NMe, 3H), 2.69 (s, NMe, 3H), 2.68 (s, NMe, 3H).

**JW72**

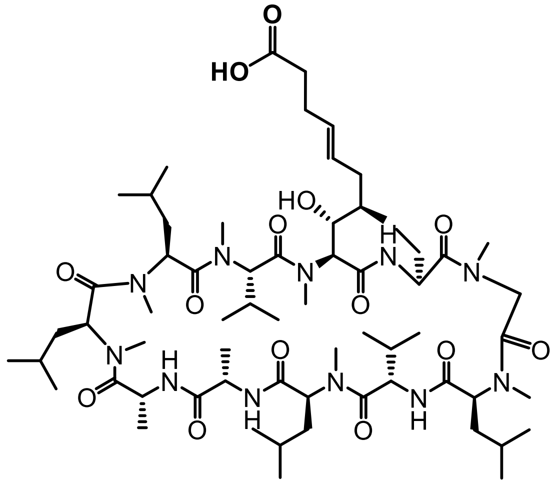

The crude product was purified by flash silica chromatography (5% MeOH in DCM) to give JW72 as an off-white powder (322 mg, 62%).

MS (m/z): [MH]+ calcd. for $C_{64}H_{113}N_{11}O_{14}$, 1260.65; found 1261.84.

$^1$H NMR (600 MHz, $CDCl_3$) δ 3.39 (s, NMe, 3H), 3.33 (s, NMe, 3H), 3.21 (s, NMe, 3H), 3.13 (s, NMe, 3H), 3.06 (s, NMe, 3H), 2.72 (s, NMe, 3H), 2.65 (s, NMe, 3H).

**CsA-R3**

All CsA modifications at the R3 position (see below) were obtained following the general method B described below.

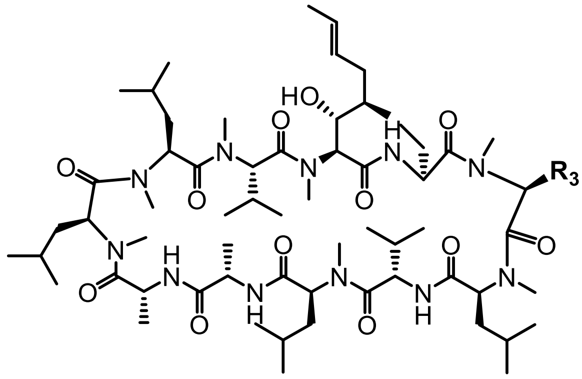



**General Method B**

To a stirred solution of CsA (1.00g, 0.83 mmol) in dry THF (25 ml) under nitrogen at 0°C was added dropwise fresh LDA (4.6 mmol, 2.3 ml, 2 M in THF). To the resultant deep-brown suspension was added, dropwise, trimethylsilylchloride (0.83 mmol, 0.1 ml) to give a clear brown solution. The mixture was stirred at 0°C for 10 min. Then further LDA (7.1 mmol, 3.5 ml, 2 M in THF) was added dropwise and the reaction stirred for 30 min at 0°C. A solution of the corresponding methyl bromo carboxylate (7 eq) in dry THF (10 ml) was added dropwise to give a pale yellow solution which was stirred for a further 1 h at room temperature (22°C). The reaction was quenched with saturated $NH_4Cl$ (aq) (10 ml) followed by 2M HCl and then diluted with DCM (dichloromethane) (20 ml). The separated aqueous layer was extracted with DCM (2x20 ml). The combined organic phases were washed with 2 M HCl (aq) (2x20 ml), saturated $NH_4Cl$ (aq) (2x20 ml) and saturated NaCl (2x20 ml) and then dried ($MgSO_4$). The volatiles were removed *in vacuo* to leave a dark brown oil residue. Purification by flash chromatography eluting with 6% (v/v) methanol in DCM gave the final product as a yellow residue.

The free carboxylic acid version of the previous product was obtained by dissolving the previous yellow solid residue (0.93 g) in THF/methanol (1:1, 20 ml) at 0◦C followed by a solution of $LiOH\cdot H_2O$ (500 mg) in water (10 ml). The reaction was allowed to gradually warm-up to room temperature over 18 h. Then DCM (20 ml) was added. The resultant solution was acidified with 2 M HCl aq (pH 3). The separated aqueous layer was extracted with DCM (3x30 ml). The combined organic extracts were washed with saturated 2 M HCl (aq) (2x30 ml) and saturated NaCl (2x30 ml) and then dried ($MgSO_4$). The volatiles were removed *in vacuo* to leave a solid residue as a mixture of the acid and unreacted CsA. The acid was separated from the CsA by flash column chromatography through an amine column eluting with a mixture of methanol/DCM/NH3 aq (1:8:1) to give the acid as a salt. After stirring the salt in DCM (20 ml) and 2 M HCl (aq) (20ml) for 10min, extraction by DCM(3x20 ml), concentration, and purification by flash column chromatography gave the final product.

**JW50**

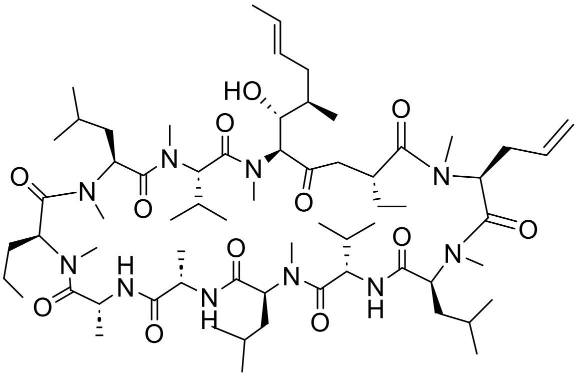


The crude product was purified by flash silica chromatography (5% → 30% acetone in DCM) to give the final product as an off-white powder (1.144 g, 18%).

$^{1}H$ NMR (600 MHz, $CDCl_3$) δ 3.50 (s, NMe, 3H), 3.26 (s, NMe, 3H), 3.25 (s, NMe, 3H), 3.09 (s, NMe, 3H), 3.07 (s, NMe, 3H), 2.69 (s, NMe, 3H), 2.67 (s, NMe, 3H).

13C NMR (600 MHz, CDCl3, δ ppm) δ 174.75 (C=O), 174.55 (C=O), 173.95 (C=O), 173.75 (C=O), 173.57 (C=O), 171.77 (C=O), 171.27 (C=O), 170.53 (C=O), 170.45 (C=O), 170.31 (C=O), 170.26 (C=O).

**JW65.2**

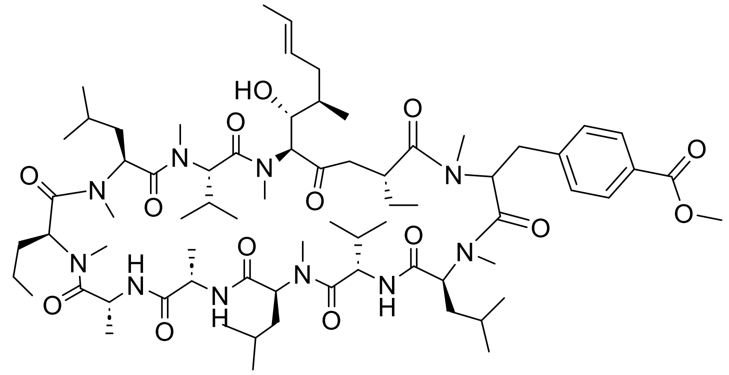

The crude product was purified by flash silica chromatography (0% → 25% acetone in DCM) to give the final product as pale yellow solid (110 mg, 16%).

$^1$H NMR (600 MHz, $CDCl_3$) δ 3.90 (s, NMe, 3H), 3.51 (s, NMe, 3H), 3.34 (s, NMe, 3H), 3.23 (s, NMe, 3H), 3.09 (s, NMe, 3H), 2.68 (s, NMe, 3H), 2.67 (s, NMe, 3H).

13C NMR (600 MHz, $CDCl_3$, δ ppm) δ 174.55 (C=O), 173.97 (C=O), 173.92 (C=O), 173.79 (C=O), 173.58 (C=O), 171.73 (C=O), 171.26 (C=O), 170.55 (C=O), 170.49 (C=O), 170.32 (C=O), 170.17 (C=O).

**JW66**

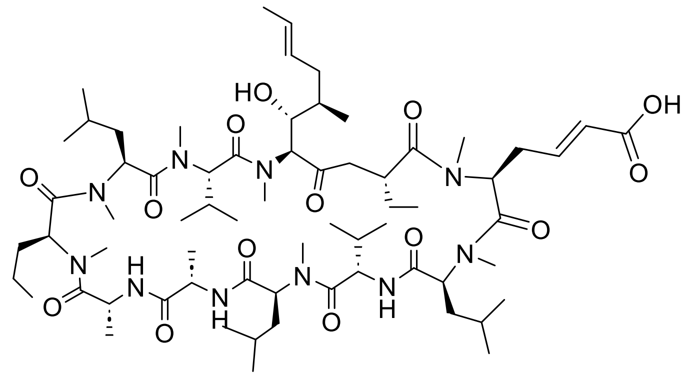

The crude product was purified by flash silica chromatography (10% → 20% MeOH in DCM) to give the final product as off white solid (45 mg, 53%).

$^1$H NMR (600 MHz, $CDCl_3$) δ 3.50 (s, NMe, 3H), 3.26 (s, NMe, 3H), 3.25 (s, NMe, 3H), 3.11 (s, NMe, 3H), 3.06 (s, NMe, 3H), 2.69 (s, NMe, 3H), 2.68 (s, NMe, 3H).

$^{13}$C-NMR (600 MHz, $CDCl_3$, δ ppm) δ 174.55 (C=O), 173.97 (C=O), 173.92 (C=O), 173.79 (C=O), 173.58 (C=O), 171.73 (C=O), 171.26 (C=O), 170.55 (C=O), 170.49 (C=O), 170.32 (C=O), 170.17 (C=O).

### 2.0 Synthesis of CsA-$BF_3K$ and the final compounds.

### BG158

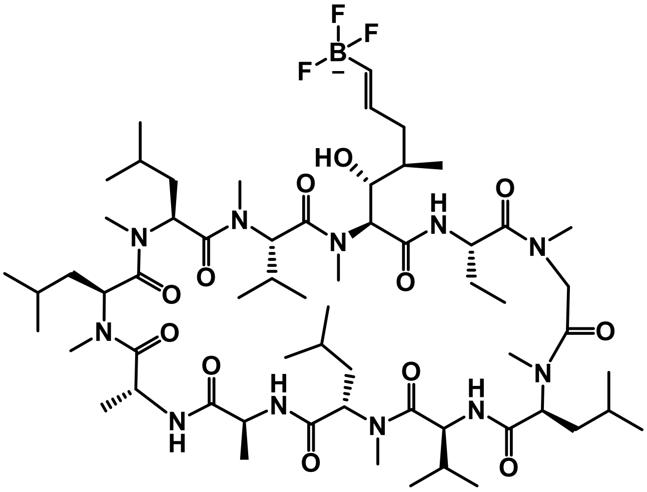


To a solution of Cyclosporin A (1 g, 0.83 mmol) in DCE (10 mL) was added vinylboronic acid pinacol ester (58 mg, 0.38 mmol) and Hoveyda-Grubbs 2nd generation catalyst (6 mol%). The reaction mixture was stirred in a MW reactor (70 ºC, 30 min) under $N_2$ atmosphere and then allowed to cool. The solvent was removed under reduce pressure and the row material was dissolved in MeOH, passed through a Stratospheres PL Thiol MP SPE cartridge (polymer Lab, Varían Inc) to remove the catalyst and concentrated under vacuum. The crude product (1.1 g, 0.83 mmol) was dissolved in MeOH (10 mL) and potassium hydrogen fluoride solution (4.5 M, 1 mL) was added. The reaction mixture was allowed to stir at r.t. overnight, concentrated under reduced pressure, dissolved in acetone (10 mL x 3) and filtered. The filtrate was then precipitated into $Et_2O$ (50 mL) to give an orange solid which was collected by filtration, washed with $Et_2O$ (50 mL) and concentrated under reduced pressure to give the final product (50 mg, 0.04 mmol, 5%).

**The compound was used as crude and not further characterized.**

### First Route for the synthesis of the compounds of the BG series:

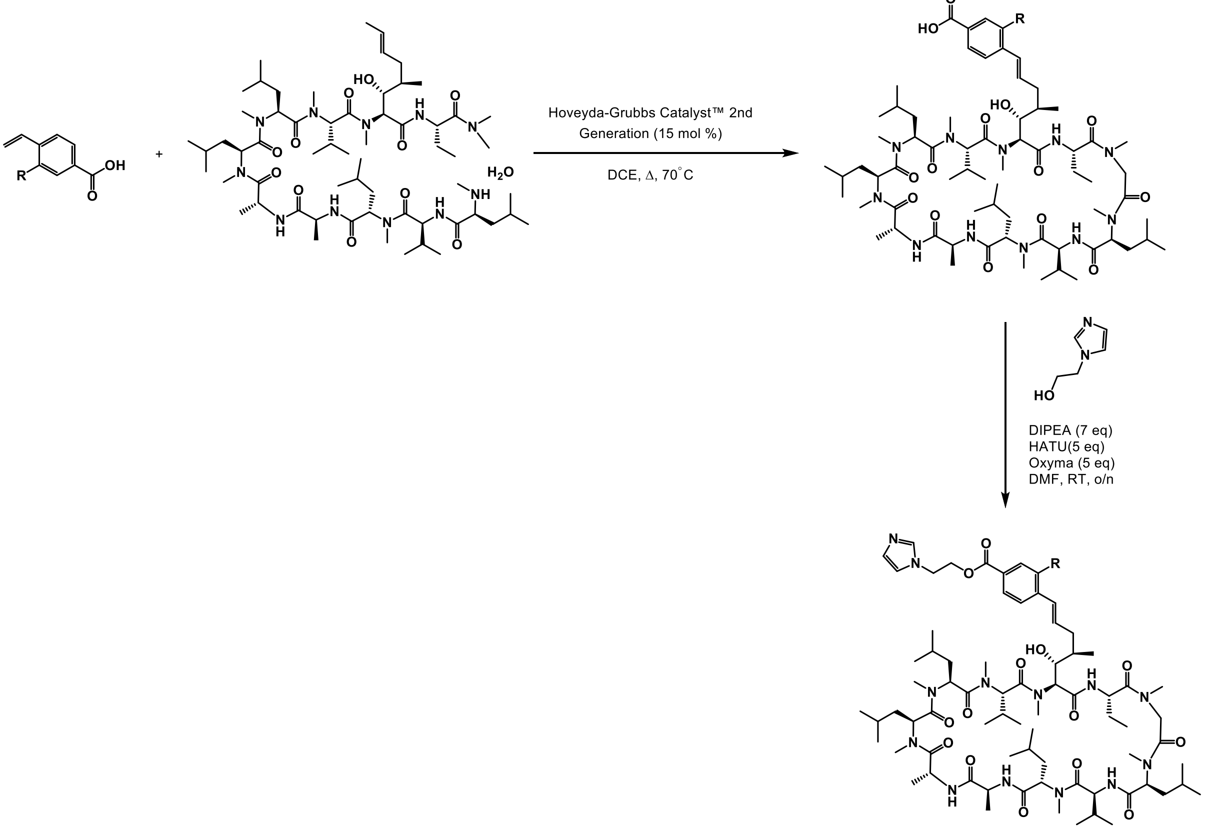


The corresponding acid (144.8 mg, 0.893 mmol, 3 eq) was dissolved in anhydrous DCE (12.0 mL) and Hoveyda-Grubbs Catalyst$^{TM}$ (27.2 mg, 0.043 mmol, 0.150 eq) was added under $N_2$ atmosphere. The reaction mixture was sparged with $N_2$ for 15 minutes, before being heated overnight at 70°C. After this time has lapsed, the solvent was removed in *vacuo* and the crude reaction mixture was placed on a small SiO2 plug which is eluted with EtOAc (100 mL). The organic solvent was removed and redissolved in DMF (0.5 mL) and wet loaded onto a reverse phase column (C18, MeCN 10% → 100% in $H_2O$ + 10 mM $NH_3$).

The presence of the corresponding product was confirmed *via* LCMS. Due to solubility issues the products were characterized after the next synthetic steps.

### General method for the esterification of the final compounds of the BG series.

The corresponding carboxylic acid (1 eq) was dissolved in a solution THF:Water = 9:1 (1 mL) followed by the corresponding 2-(lH-imidazol-1-yl)ethyl4-bromo-arylbenzoate (2 eq.), caesium carbonate (6 eq) and the catalyst Pd(dppf)Cb.DCM (10 mol%). The reaction mixture was stirred at 90 ºC for 30 min under $N_2$ atmosphere, warmed to r.t. and diluted with $H_2O$. The organic layer was separated and the aqueous phase extracted with DCM (x3). The combined organic layers were washed with Brine (x3), dried over $Mg_2SO_4$, filtered and concentrated under reduced pressure. The crude product was purified by reverse phase C18 (0-100% MeCN 0.1% FA+ 15 % MeOH in $H_2O$ 0.1% FA) obtaining the final product.

**BG147**

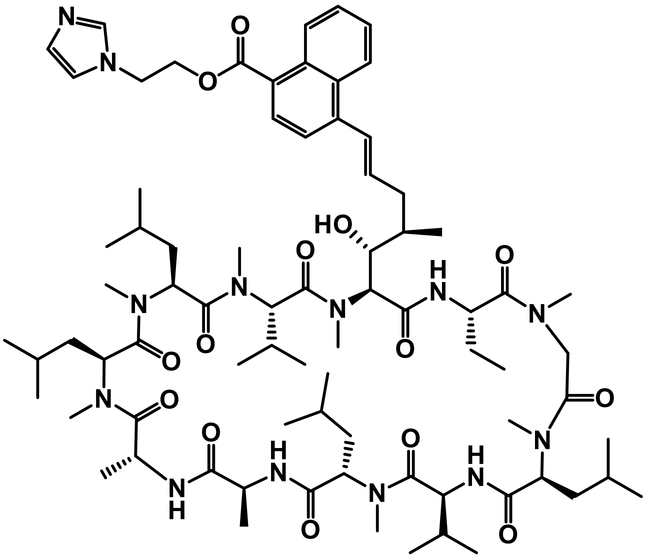

7% ; off-white solid.

LCMS (m/z): [MH]+ calc. for $C_{77}H_{121}N_{13}O_{14}$ 1452.9, found 1452.9.

**$^{1}$H-NMR (600 MHz, $CDCl_3$, δ ppm)** 3.55 (s, NMe, 3H), 3.39 (s, NMe, 3H), 3.27 (s, NMe, 3H), 3.10 (s, NMe, 3H), 3.09 (s, NMe, 3H), 2.70 (s, NMe, 3H), 2.66 (s, NMe, 3H).

**$^{13}$C-NMR (500 MHz, $CDCl_3$, δ ppm)** δ 173.92 (C=O), 173.89 (C=O), 173.66 (C=O), 171.71 (C=O), 171.50 (C=O), 171.38 (C=O), 171.34 (C=O), 170.63 (C=O), 170.56 (C=O), 170.45 (C=O), 170.37 (C=O).

**BG149**

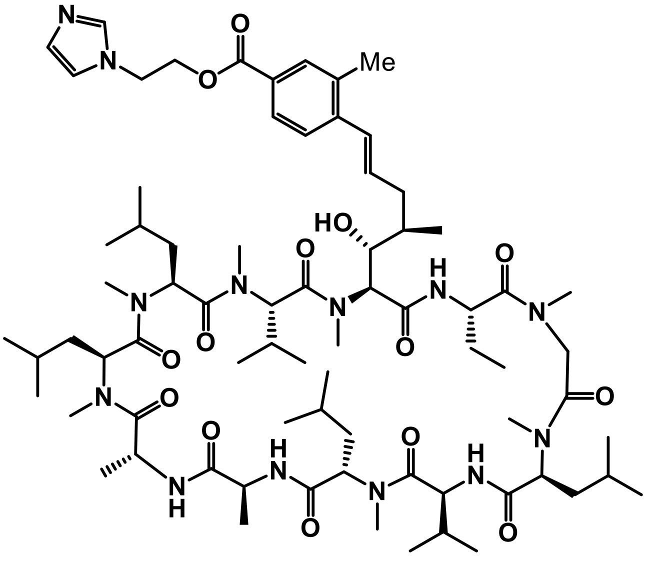

7%; orange oil

LCMS (m/z): [M+2H/2] calc. for $C_{74}H_{121}N_{13}O_{14}$ 1416.9, found 709.1.

**$^{1}$H-NMR (700 MHz, CDCl$_3$, δ ppm)** 3.53 (s, NMe, 3H), 3.39 (s, NMe, 3H), 3.26 (s, NMe, 3H), 3.11 (s, NMe, 3H), 3.10 (s, NMe, 3H), 2.71 (s, NMe, 3H), 2.67 (s, NMe, 3H), 2.34 (s, ArCH3, 3H).

**$^{13}$C-NMR (500 MHz, CDCl$_3$, δ ppm)** 173.95 (C=O), 173.94 (C=O), 173.94 (C=O), 173.92 (C=O), 173.63 (C=O), 171.69 (C=O), 171.54 (C=O), 171.32 (C=O), 170.55 (C=O), 170.49 (C=O), 170.32 (C=O).

#### 2.1. Synthesis and characterization of BG147 its analogues (VP series).

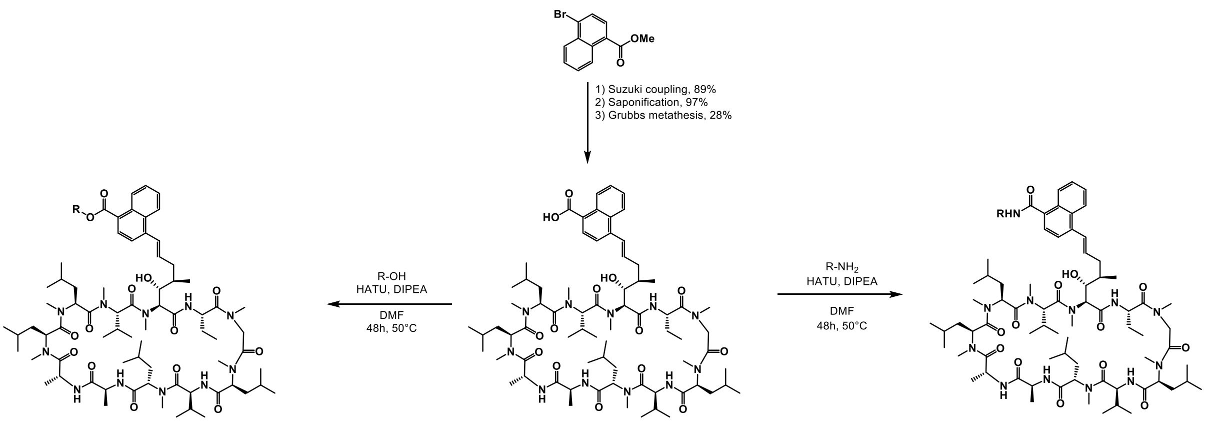


***Scheme 2.1.*** Synthesis of **BG147** and its analogues.

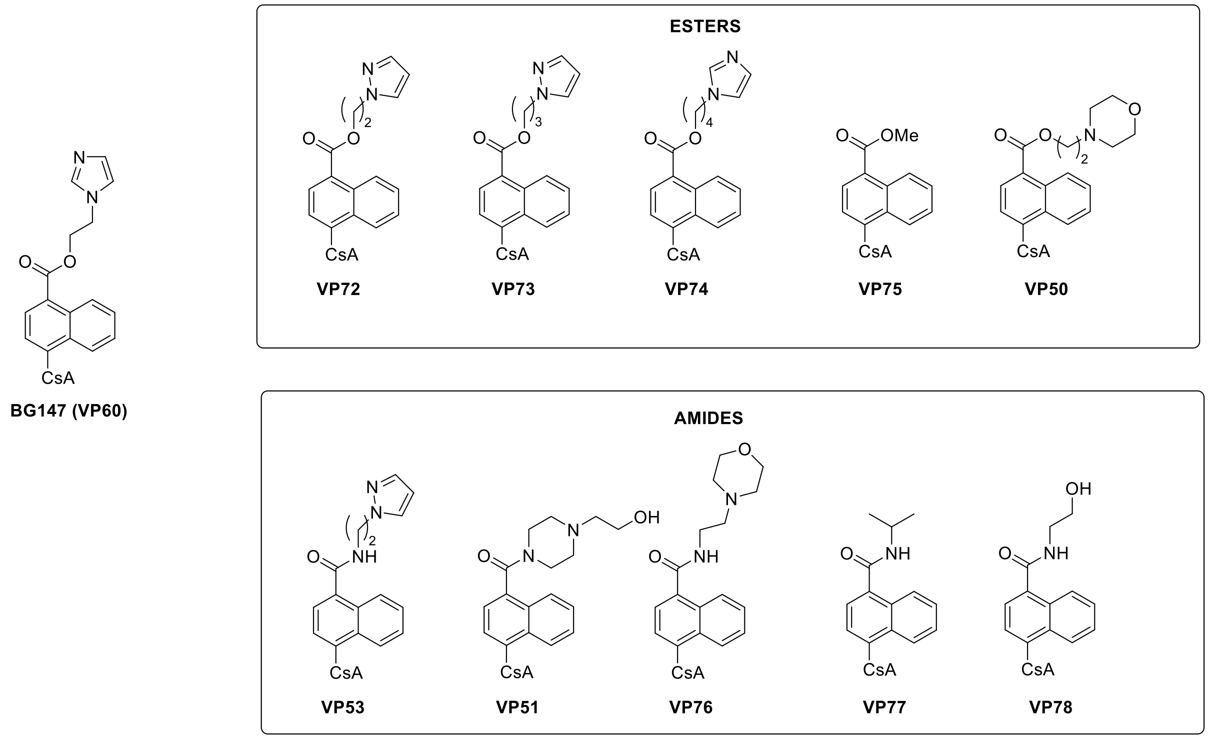


***Figure 1.*** Structures of **BG147** and its esters and amides analogues.

**VP43**

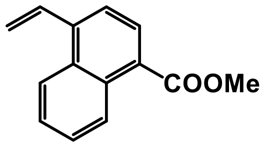


Methyl 4-bromonaphthalene-1-carboxylate (530 mg, 2 mmol) was dissolved in THF:Water= 9:1 (28 mL) followed by potassium vinyltrifluoroborate (1.3 g, 10 mmol),Cesium carbonate (1.9 g, 6 mmol), and [1,1′Bis(diphenylphosphino)ferrocene]dichloropalladium(II) (1:1) (5 mol%). After stirring overnight at 70 °C, the reaction mixture was cooled and condensed *in vacuo*. The residue was diluted with $CH_2Cl_2$ (10 mL), washed with water (3x10 mL) and brine, dried over $MgSO_4$ and concentrated under reduced pressure. The crude product was purified by chromatography column on silica gel (EtOAc 0% → 100% in Cyclohexane) to afford VP43 (378 mg, 1.779 mmol, 89%) as a yellow oil.

The data matches previous literature*(64)*.

**VP57**

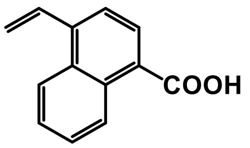


VP43 (2.2 g, 10 mmol) was dissolved in THF (12 mL) and treated with a solution 5M of LiOH (25 mL). After stirring overnight at 50 °C, THF removed under vacuum and the residue was acidified with HCl 1M up to pH 2, extracted with EtOAc (30x3), dried over $MgSO_4$, filtered and concentrated under reduced pressure to afford VP57 (2 g, 10 mmol, 97%) as a yellow solid.

The data matches previous literature*(65)*.

**VP47**

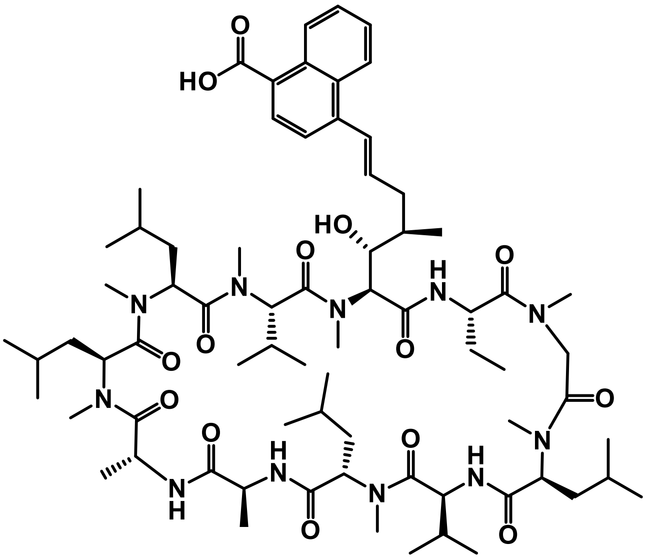

VP57 (270 mg, 1.36 mmol) was dissolved into anhydrous DCE (4.5 mL) followed by Cyclosporine A (360 mg, 0.299 mmol) and Hoveyda-Grubbs 2nd generation catalyst (15 mol%). The reaction mixture was stirred under $N_2$ atmosphere in a MW reactor (70 °C, 30 min)

and then cooled to r.t. The solvent was removed under reduced pressure and the row material was redissolved in MeOH, passed through a Stratospheres PL Thiol MP SPE cartridge (polymer Lab, Varian Inc) to remove the catalyst. The unreacted CsA was removed by Biotage Isolute PE-AX 10g (MeOH + 10% $NH_4OH$ → 100% MeOH → 5% MeOH in DCM→ 5% MeOH + 5% AcOH in DCM). The excess of VP57 was removed by reverse phase chromatography (C18, MeCN 10% → 100% in $H_2O$ + 10 mM $NH_3$) obtaining VP47 (115 mg, 0.085 mmol, 28%, trans/cis=3/1) as an off- white solid.

$^1$H-NMR (500 MHz, CDl3, δ ppm) δ 3.56 (s, NMe, 3H), 3.40 (s, NMe, 3H), 3.27 (s, NMe, 3H), 3.11 (s, NMe, 3H), 3.10 (s, NMe, 3H), 2.72 (s, NMe, 3H), 2.67 (s, NMe, 3H).

$^{13}$C-NMR (500 MHz, $CDCl_3$, δ ppm) δ 173.92 (C=O), 173.89 (C=O), 173.66 (C=O), 171.71 (C=O), 171.50 (C=O), 171.38 (C=O), 171.34 (C=O), 170.63 (C=O), 170.56 (C=O), 170.45 (C=O), 170.37 (C=O).

LCMS (*m/z*): $[MH]^+$ calcd. for $C_{72}H_{115}N_{11}O_{14}$, 1358.9; found 1358.4

**General procedure for the synthesis of BG147 and its ester and amide analogues.**

To a solution of VP47 (1 eq) in DMF (1 mL) were added DIPEA (7 eq) and HATU (6 eq). After stirring for 10 min at r.t., a solution of the appropriate alcohol or amine (10 eq) in DMF (0.5 mL) was added and the reaction mixture was stirred overnight at 50 °C. Then, the solvent was removed under reduced pressure and the crude product was purified by chromatography column on silica gel (MeOH 0% →10% in DCM) and by reverse phase C4 (MeCN 30% → 100% in $H_2O$ 10 mM $NH_3$) obtaining the final product.

**BG147**

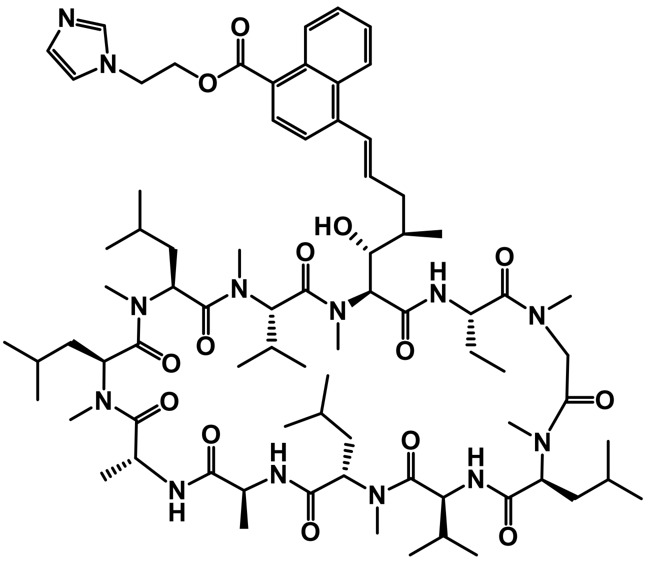

BG147 was obtained as an off-white solid (17 mg, 0.01 mmol, 20%, trans/cis=3/1) following the general procedure described above and using 2-(1H-imidazol-1-yl)ethan-1-ol as appropriate alcohol.

$^1$H-NMR (600 MHz, CDl3, δ ppm) δ 3.55 (s, NMe, 3H), 3.39 (s, NMe, 3H), 3.27 (s, NMe, 3H), 3.10 (s, NMe, 3H), 3.09 (s, NMe, 3H), 2.70 (s, NMe, 3H), 2.66 (s, NMe, 3H).

$^{13}$C-NMR (600 MHz, $CDCl_3$, δ ppm) δ 173.97 (C=O), 173.88 (C=O), 173.63 (C=O), 173.13 (C=O), 172.80 (C=O), 172.16 (C=O), 171.54 (C=O), 171.43 (C=O), 171.26 (C=O), 170.53 (C=O), 170.28 (C=O).

HRMS (*m/z*): $[MH]^+$ calcd. for $C_{77}H_{121}N_{13}O_{14}$, 1452.91560; found 1452.92287

**VP50**

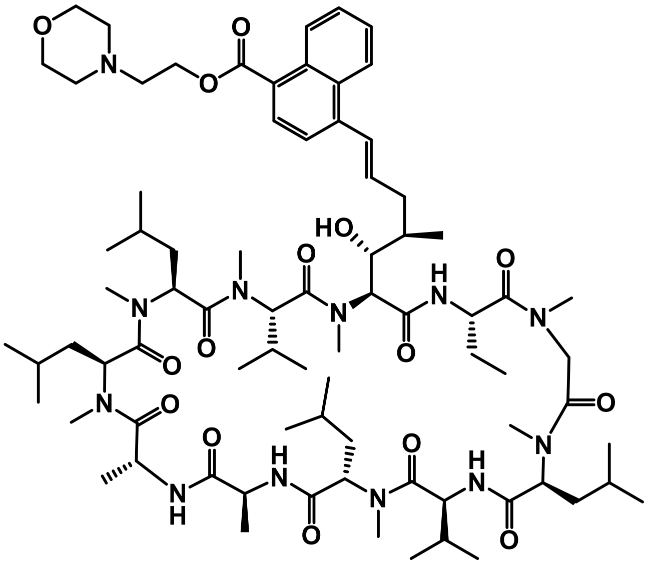

VP50 was obtained as an off-white solid (3.0 mg, 0.002 mmol, 17%, trans/cis=3/1) following the general procedure described above and using 2-morpholinoethan-1-ol as appropriate alcohol.

$^1$H-NMR (600 MHz, $CDCl_3$, δ ppm) δ 3.56 (s, NMe, 3H), 3.41 (s, NMe, 3H), 3.28 (s, NMe, 3H), 3.11 (s, NMe, 3H), 3.10 (s, NMe, 3H), 2.70 (s, NMe, 3H), 2.66 (s, NMe, 3H).

LCMS (*m/z*): $[MH]^+$ calcd. for $C_{78}H_{126}N_{12}O_{15}$. 1471.9; found 1471.6

**VP51**

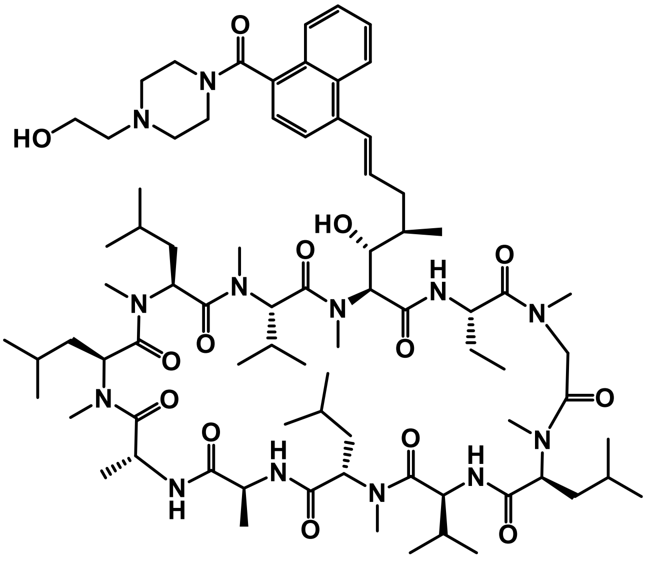

VP51 was obtained as an off-white solid (3.0 mg, 0.002 mmol, 18%, trans/cis=3/1) following the general procedure described above and using 2-(piperazin-1-yl)ethan-1-ol as appropriate amine.

$^{1}$H-NMR (600 MHz, $CDCl_3$, δ ppm) δ 3.55 (s, NMe, 3H), 3.41 (s, NMe, 3H), 3.27 (s, NMe, 3H), 3.12 (s, NMe, 3H), 3.00 (s, NMe, 3H), 2.71 (s, NMe, 3H), 2.67 (s, NMe, 3H).

$^{13}$C-NMR (600 MHz, $CDCl_3$, δ ppm) δ 173.92 (C=O), 173.62 (C=O), 171.64 (C=O), 171.37 (C=O), 171.27 (C=O), 170.61 (C=O), 170.56 (C=O), 170.44 (C=O), 170.30 (C=O), 170.20 (C=O), 169.71.(C=O).

LCMS (*m/z*): $[MH]^+$ calcd. for $C_{78}H_{127}N_{13}O_{14}$, 1,471.0; found 1471.2

**VP53**

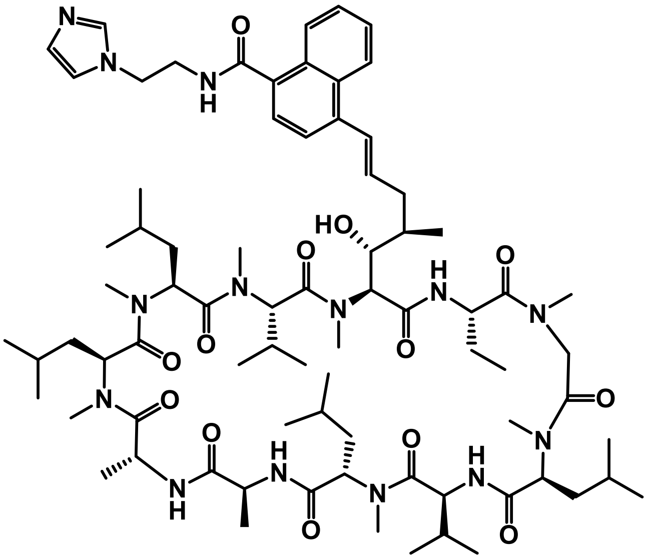


VP53 was obtained as a yellow solid (8.0 mg, 0.006 mmol, 47%, trans/cis=3/1) following the general procedure described above and using 2-(1H-imidazol-1-yl)ethan-1-amine as appropriate amine.

$^{1}$H-NMR (600 MHz, $CDCl_3$, δ ppm) δ 3.55 (s, NMe, 3H), 3.37 (s, NMe, 3H), 3.29 (s, NMe, 3H), 3.08 (s, NMe, 3H), 3.03 (s, NMe, 3H), 2.71 (s, NMe, 3H), 2.67 (s, NMe, 3H).

$^{13}$C NMR (600 MHz, $CDCl_3$, δ ppm) δ 173.99 (C=O), 173.94 (C=O), 173.89 (C=O), 173.76 (C=O), 173.67 (C=O), 173.63 (C=O), 173.57 (C=O), 171.66 (C=O), 171.40 (C=O), 171.22 (C=O), 171.21 (C=O), 171.19 (C=O).

LCMS (*m/z*): $[MH]^+$ calcd. for $C_{77}H_{122}N_{14}O_{13}$, 1451.9; found 1452.2

**VP72**

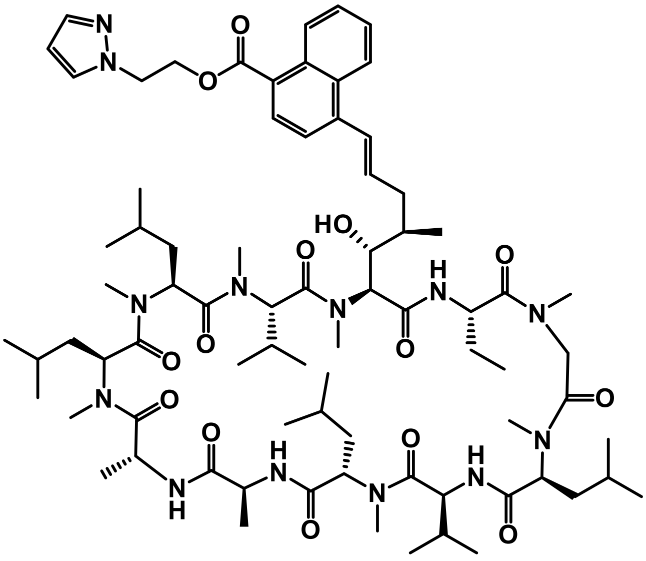

VP72 was obtained as an yellow solid (3.0 mg, 0.002 mmol, 14%, trans/cis=3/1) following the general procedure described above and using 2-(1H-pyrazol-1-yl)ethan-1-ol as appropriate alcohol.

$^1$H-NMR (500 MHz, $CDCl_3$, δ ppm) δ 3.56 (s, NMe, 3H), 3.41 (s, NMe, 3H), 3.27 (s, NMe, 3H), 3.11 (s, NMe, 3H), 3.10 (s, NMe, 3H), 2.71 (s, NMe, 3H), 2.67 (s, NMe, 3H).

LCMS (*m/z*): $[MH]^+$ calcd. for $C_{77}H_{121}N_{13}O_{14}$, 1451.9; found 1453.0

**VP73**

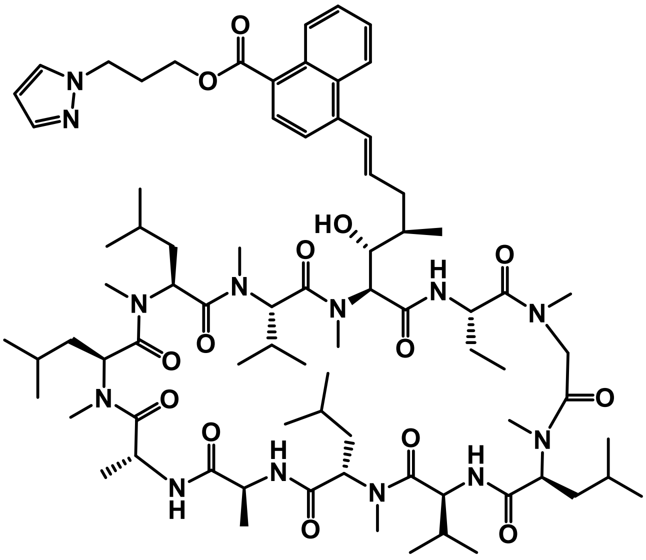

VP73 was obtained as a yellow solid (3.0 mg, 0.002 mmol, 14%, trans/cis=3/1) following the general procedure described above and using 3-(1H-pyrazol-1-yl)propan-1-ol as appropriate alcohol.

$^1$H-NMR (500 MHz, $CDCl_3$, δ ppm) δ 3.56 (s, NMe, 3H), 3.41 (s, NMe, 3H), 3.28 (s, NMe, 3H), 3.11 (s, NMe, 3H), 3.10 (s, NMe, 3H), 2.71 (s, NMe, 3H), 2.67 (s, NMe, 3H).

LCMS (*m/z*): $[MH]^+$ calcd. for $C_{78}H_{123}N_{13}O_{14}$, 1465.9; found 1467.1

**VP74**

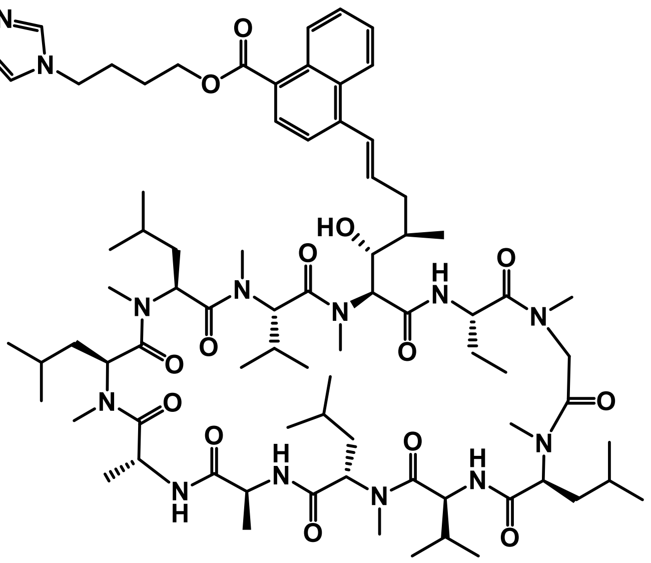


VP74 was obtained as a yellow solid (4.0 mg, 0.003 mmol, 18%, trans/cis=1/1) following the general procedure described above and using 4-(1H-imidazol-1-yl)butan-1-ol as appropriate alcohol.

$^1$H-NMR (500 MHz, $CDCl_3$, δ ppm) δ 3.55 (s, NMe, 3H), 3.40 (s, NMe, 3H), 3.28 (s, NMe, 3H), 3.11 (s, NMe, 3H), 3.10 (s, NMe, 3H), 2.71 (s, NMe, 3H), 2.67 (s, NMe, 3H).

LCMS (*m/z*): $[MH]^+$ calcd. for $C_{79}H_{125}N_{13}O_{14}$, 1479.9; found 1481.1

**VP75**

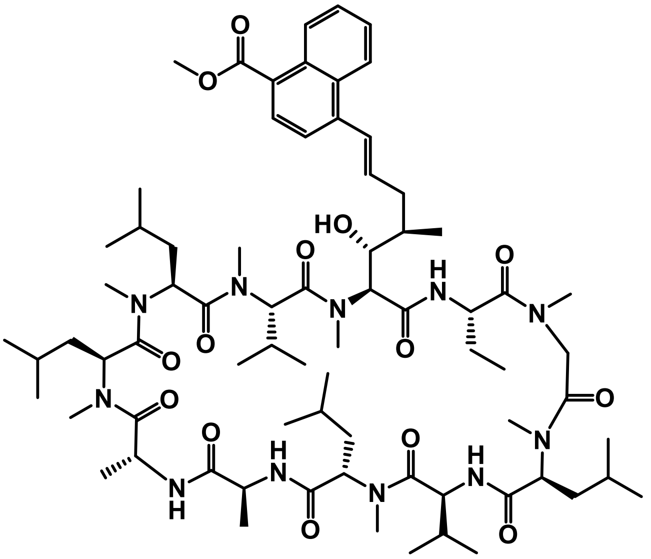

**VP75** was obtained as a yellow solid (11.0 mg, 0.008 mmol, 54%, trans/cis=3/1) following the general procedure described above and using methanol as appropriate alcohol.

$^{1}$H-NMR (500 MHz, $CDCl_3$, δ ppm) δ 3.56 (s, NMe, 3H), 3.40 (s, NMe, 3H), 3.27 (s, NMe, 3H), 3.11 (s, NMe, 3H), 3.10 (s, NMe, 3H), 2.71 (s, NMe, 3H), 2.67 (s, NMe, 3H).

LCMS (*m/z*): $[MH]^+$ calcd. for $C_{73}H_{117}N_{11}O_{14}$, 1371.9; found 1373.0

**VP76**

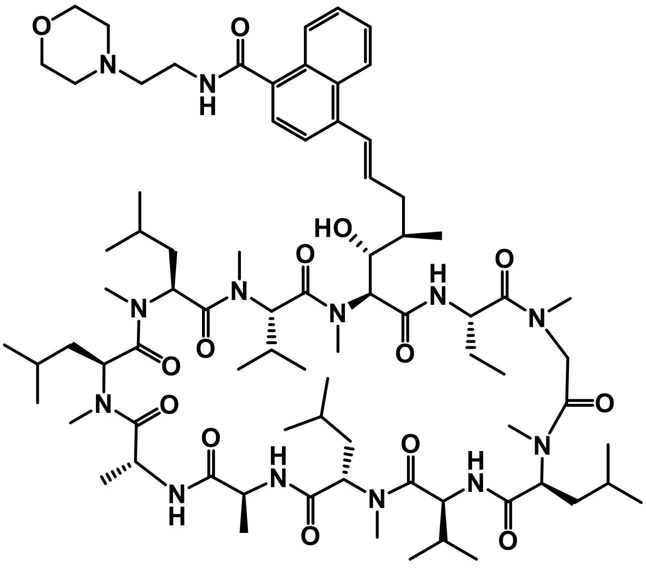

VP76 was obtained as an white solid (10.0 mg, 0.007 mmol, 46%, trans/cis=3/1) following the general procedure described above and using 2-morpholinoethan-1-amine as appropriate amine.

$^{1}$H-NMR (500 MHz, $CDCl_3$, δ ppm) δ 3.55 (s, NMe, 3H), 3.38 (s, NMe, 3H), 3.27 (s, NMe, 3H), 3.10 (s, NMe, 3H), 3.08 (s, NMe, 3H), 2.72 (s, NMe, 3H), 2.67 (s, NMe, 3H).

LCMS (*m/z*): $[MH]^+$ calcd. for $C_{78}H_{127}N_{13}O_{14}$, 1470.0; found 1471.3

**VP77**

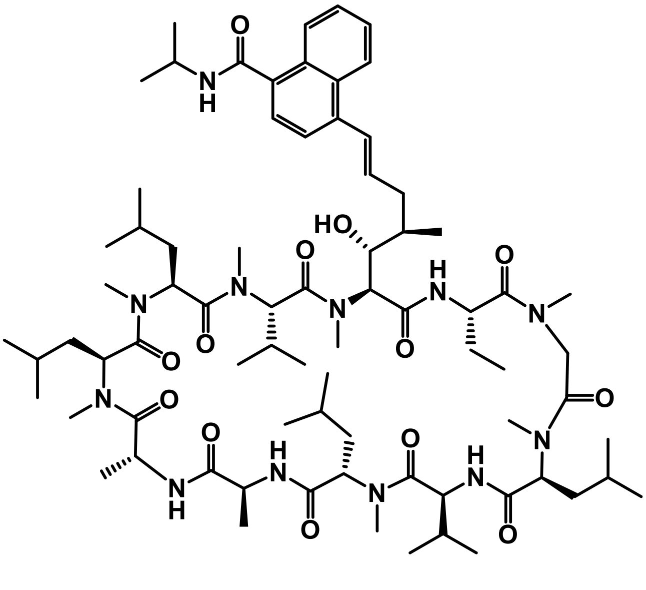

VP77 was obtained as an white solid (17.0 mg, 0.0012 mmol, 83%, trans/cis=3/1) following the general procedure described above and using propan-2-amine as appropriate amine.

$^{1}$H-NMR (500 MHz, $CDCl_3$, δ ppm) δ 3.55 (s, NMe, 3H), 3.38 (s, NMe, 3H), 3.27 (s, NMe, 3H), 3.09 (s, NMe, 3H), 3.06 (s, NMe, 3H), 2.71 (s, NMe, 3H), 2.67 (s, NMe, 3H).

$^{13}$C-NMR (500 MHz, $CDCl_3$, δ ppm) δ 173.94 (C=O), 173.91 (C=O), 173.76 (C=O), 171.61 (C=O), 171.54 (C=O), 171.50 (C=O), 171.25 (C=O), 170.55 (C=O), 170.45 (C=O), 170.39 (C=O), 170.26 (C=O).

LCMS (*m/z*): $[MH]^+$ calcd. for $C_{75}H_{122}N_{12}O_{13}$, 1398.9; found 1400.00

**VP78**

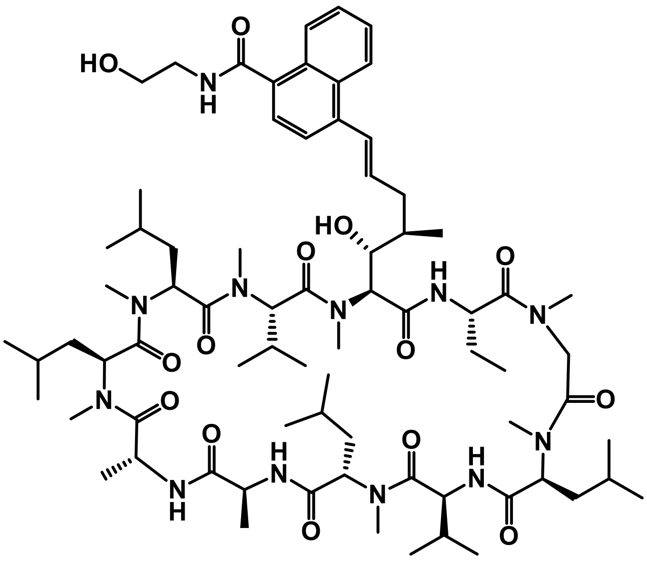


VP78 was obtained as an white solid (13.0 mg, 0.009 mmol, 63%, trans/cis=3/1) following the general procedure described above and using 2-aminoethan-1-ol as appropriate amine.

$^{1}$H-NMR (500 MHz, $CDCl_3$, δ ppm) δ 3.56 (s, NMe, 3H), 3.37 (s, NMe, 3H), 3.29 (s, NMe, 3H), 3.08 (s, NMe, 3H), 3.04 (s, NMe, 3H), 2.71 (s, NMe, 3H), 2.67 (s, NMe, 3H).

$^{13}$C-NMR (500 MHz, $CDCl_3$, δ ppm) δ 173.94 (C=O), 173.90 (C=O), 173.66 (C=O), 173.57 (C=O), 171.58 (C=O), 171.47 (C=O), 171.09 (C=O), 170.95 (C=O), 170.89 (C=O), 170.48 (C=O), 170.27 (C=O).

LCMS (*m/z*): $[MH]^+$ calcd. for $C_{74}H_{120}N_{12}O_{14}$, 1400.9; found 1402.0

### 1H-NMR, 13C-NMR and HRMS of the key compounds

### JW3-158

HRMS

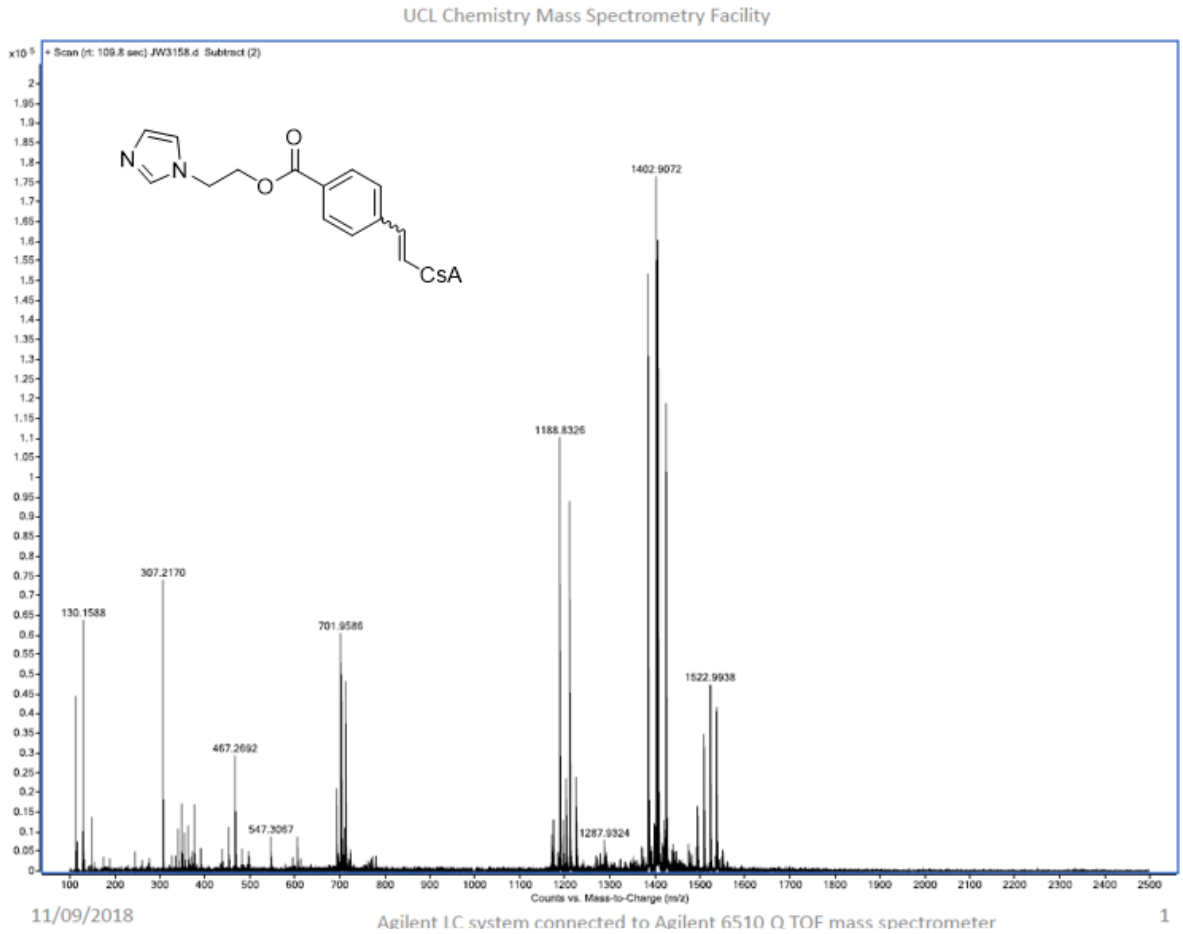

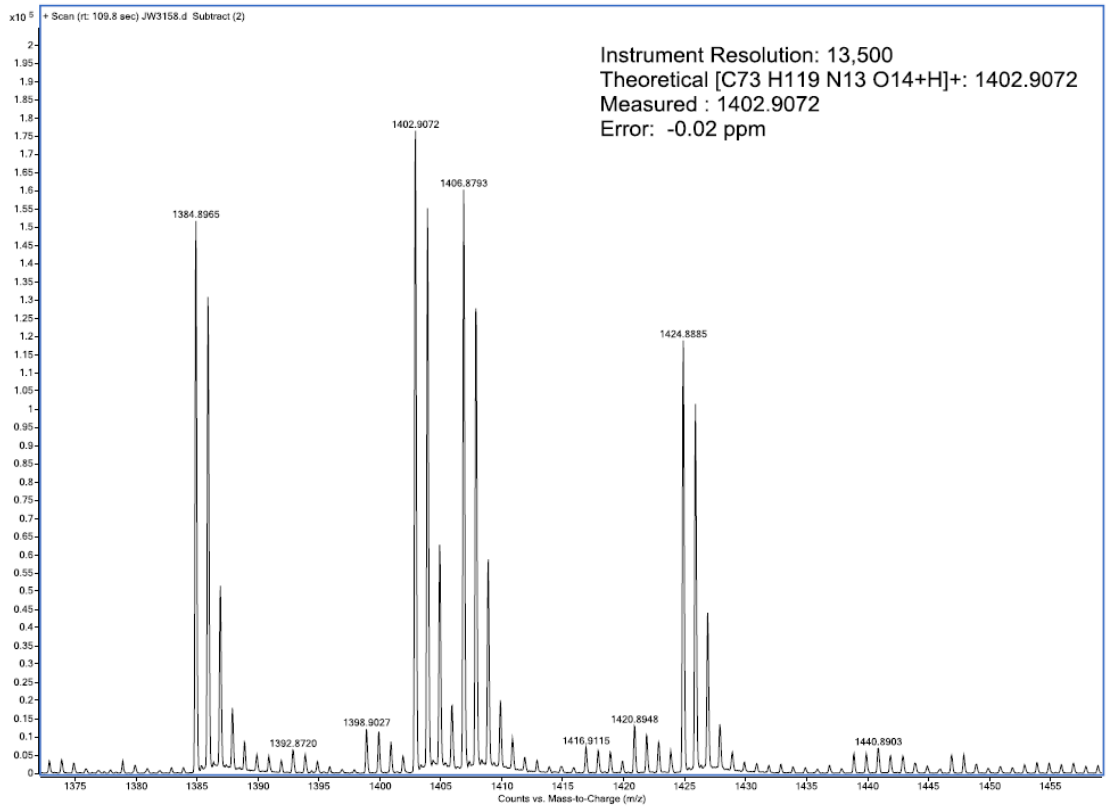


11/09/2018

Agilent LC system connected to Agilent 6510 Q TOF mass spectrometer

2

**BG147**

$^1$H-NMR

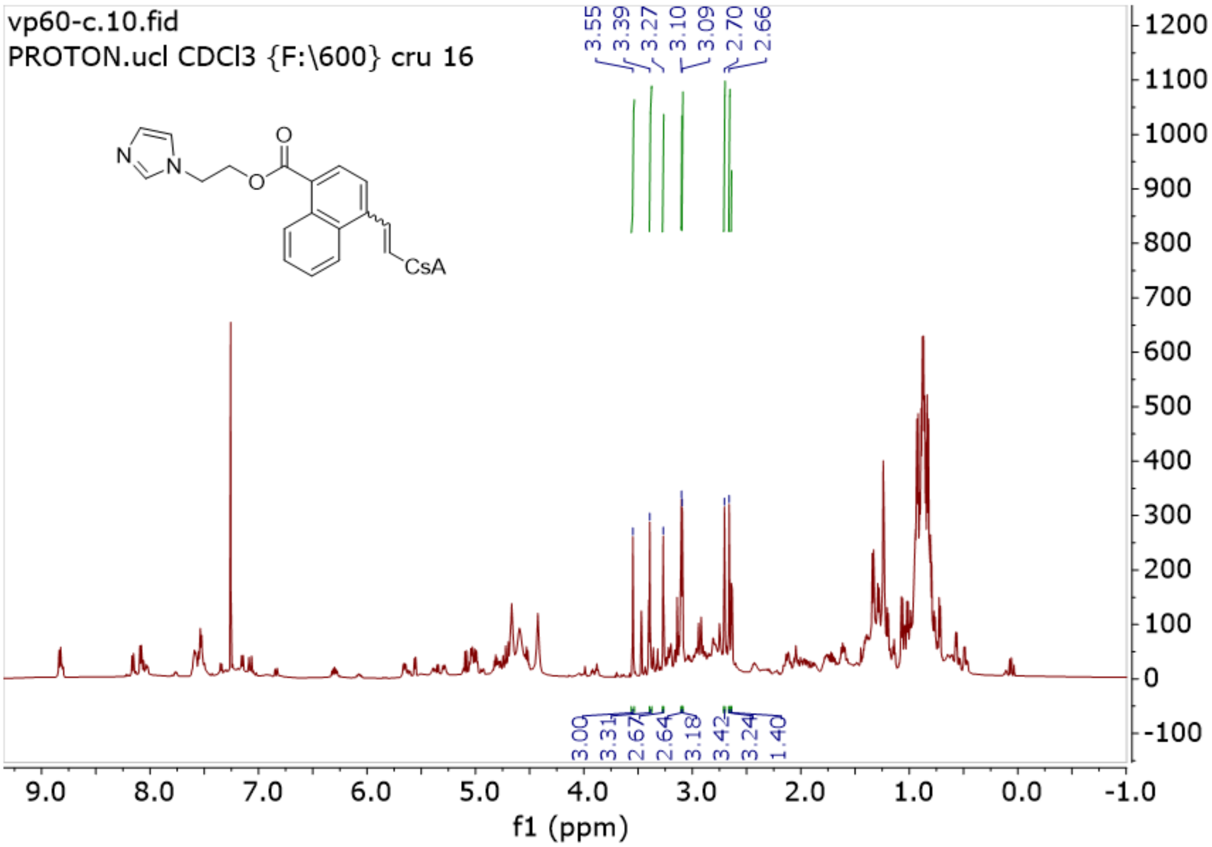
vp60-c.10.fid
PROTON.ucl CDCl3 {F:\600} cru 16
3.55 3.39 3.27 3.10 3.09 2.70 2.66
3.00 3.31 2.67 2.64 3.18 3.42 3.24 1.40
f1 (ppm)

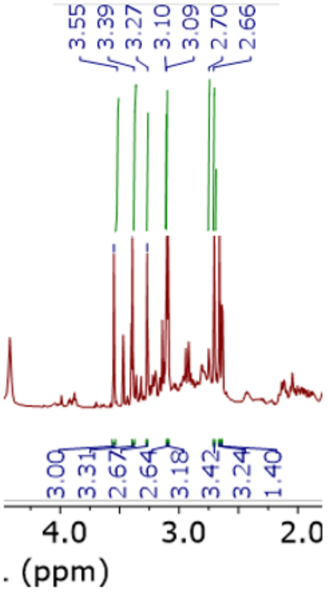
3.55 3.39 3.27 3.10 3.09 2.70 2.66
3.00 3.31 2.67 2.64 3.18 3.42 3.24 1.40
(ppm)

**BG147**

$^{13}$C-NMR

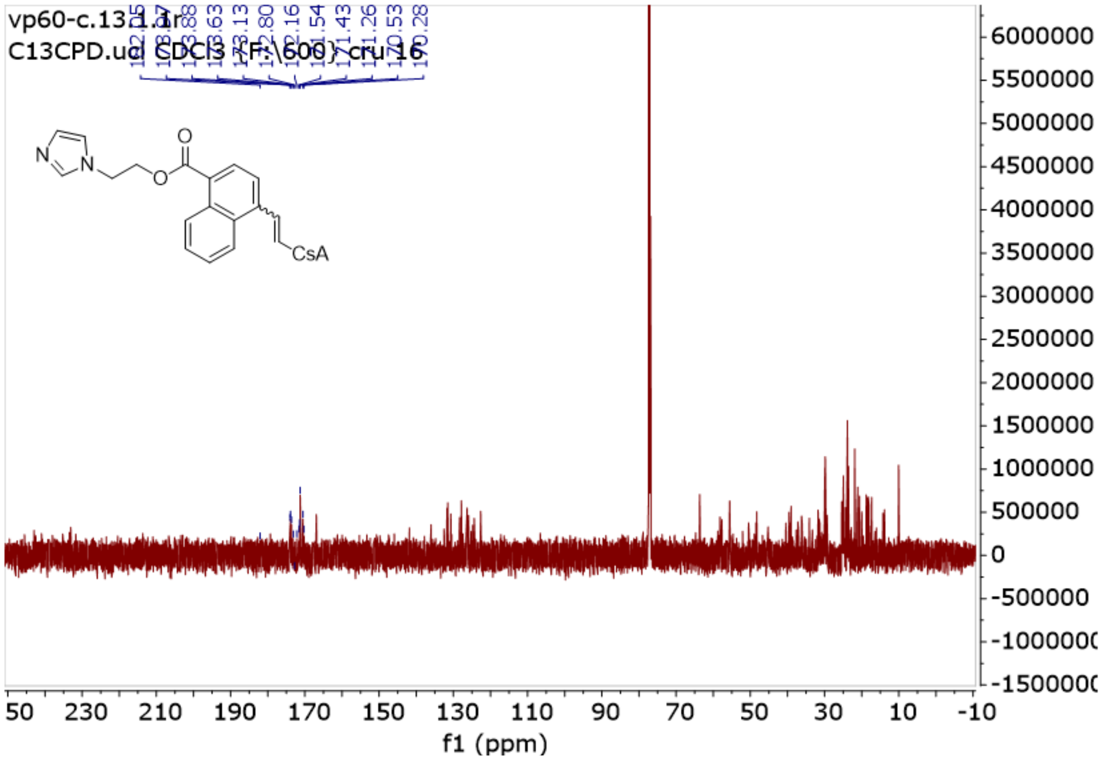

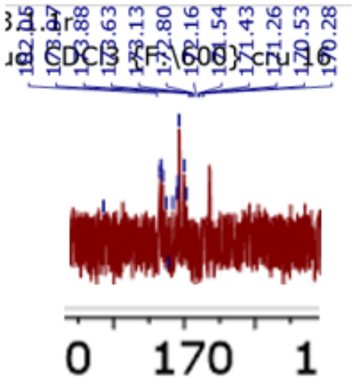

**BG147**

HRMS

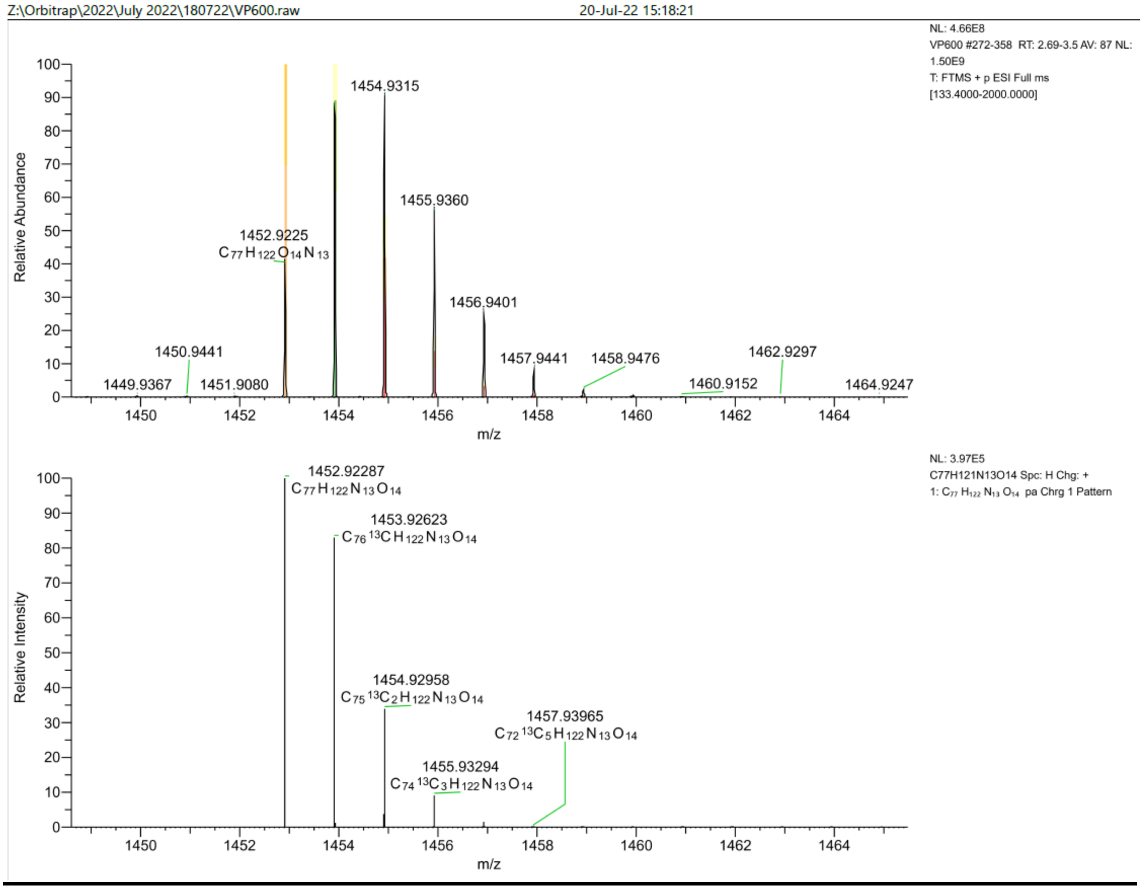
Z:\Orbitrap\2022\July 2022\180722\VP600.raw
20-Jul-22 15:18:21
NL: 4.66E8
VP600 #272-358 RT: 2.69-3.5 AV: 87 NL: 1.50E9
T: FTMS + p ESI Full ms [133.4000-2000.0000]
Relative Abundance
1454.9315
1455.9360
1452.9225
C77 H122 O14 N13
1456.9401
1450.9441
1457.9441
1458.9476
1462.9297
1449.9367
1451.9080
1460.9152
1464.9247
m/z
NL: 3.97E5
C77H121N13O14 Spc: H Chg: +
1: C77 H122 N13 O14 pa Chrg 1 Pattern
Relative Intensity
1452.92287
C77 H122 N13 O14
1453.92623
C76 13C H122 N13 O14
1454.92958
C75 13C2 H122 N13 O14
1457.93965
C72 13C5 H122 N13 O14
1455.93294
C74 13C3 H122 N13 O14
m/z

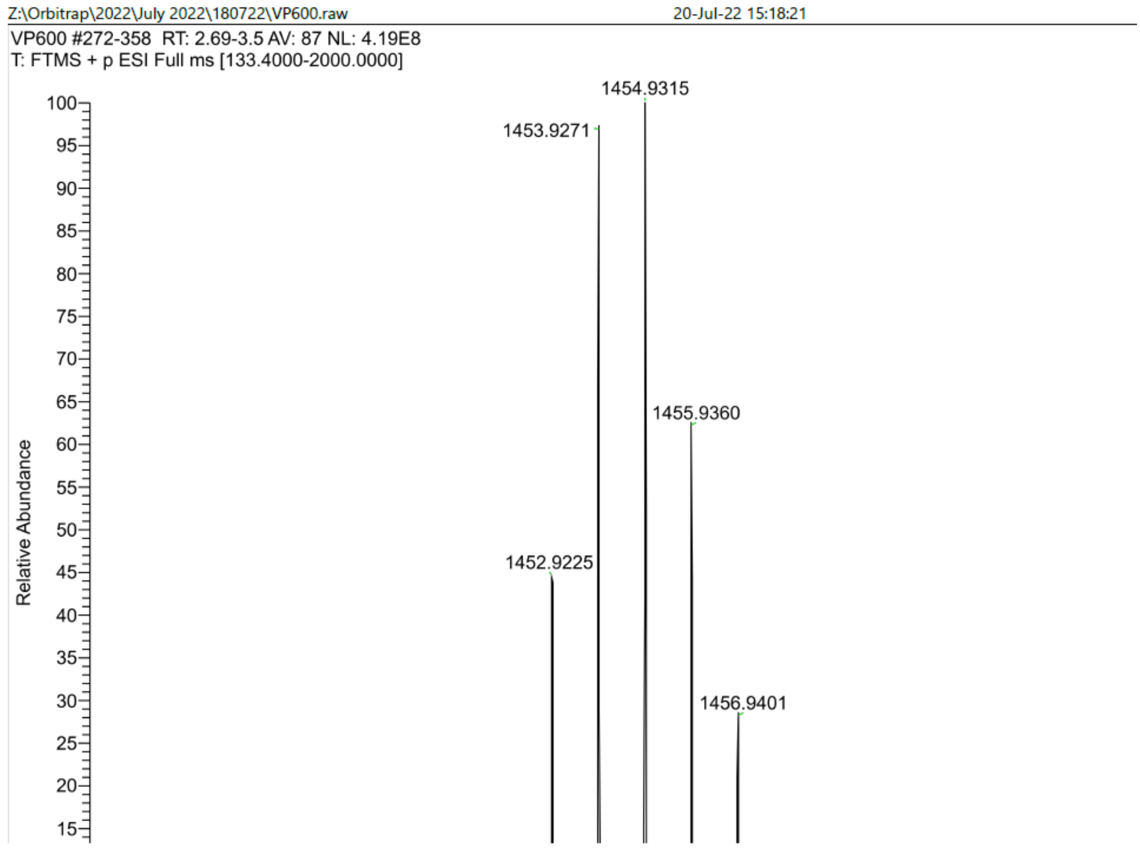
Z:\Orbitrap\2022\July 2022\180722\VP600.raw
20-Jul-22 15:18:21
VP600 #272-358 RT: 2.69-3.5 AV: 87 NL: 4.19E8
T: FTMS + p ESI Full ms [133.4000-2000.0000]
Relative Abundance
1454.9315
1453.9271
1455.9360
1452.9225
1456.9401

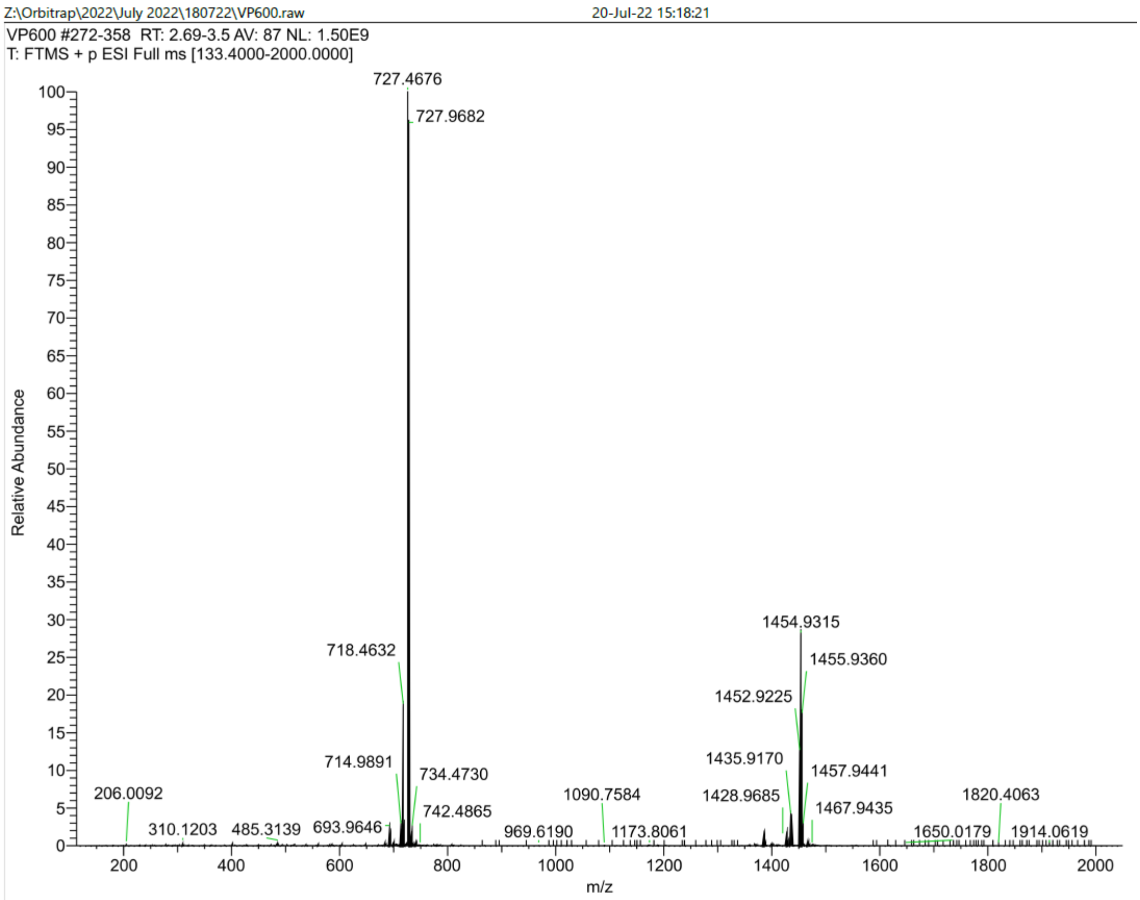
Z:\Orbitrap\2022\July 2022\180722\VP600.raw
20-Jul-22 15:18:21
VP600 #272-358 RT: 2.69-3.5 AV: 87 NL: 1.50E9
T: FTMS + p ESI Full ms [133.4000-2000.0000]
Relative Abundance
m/z
727.4676
727.9682
718.4632
714.9891
734.4730
742.4865
206.0092
310.1203
485.3139
693.9646
969.6190
1090.7584
1173.8061
1454.9315
1455.9360
1452.9225
1435.9170
1457.9441
1428.9685
1467.9435
1820.4063
1650.0179
1914.0619
